# Dynamic atP5CS2 Filament Facilitates Substrate Channeling

**DOI:** 10.1101/2023.09.07.556688

**Authors:** Chen-Jun Guo, Tianyi Zhang, Qingqing Leng, Xian Zhou, Jiale Zhong, Ji-Long Liu

## Abstract

In plants, the rapid accumulation of proline is a common response to combat abiotic stress. Delta-1-pyrroline-5-carboxylate synthase (P5CS) is a rate-limiting enzyme in proline synthesis, catalyzing the initial two-step conversion from glutamate to proline. Here, we determine the first structure of plant P5CS. Our results show that Arabidopsis thaliana P5CS2 (atP5CS2) can form enzymatic filaments in a substrate-sensitive manner. The destruction of atP5CS filaments by mutagenesis leads to a significant reduction in enzymatic activity. Furthermore, separate activity tests on two domains reveals that filament-based substrate channeling is essential for maintaining the high catalytic efficiency of atP5CS. Our study demonstrates the unique mechanism for the efficient catalysis of P5CS, shedding light on the intricate mechanisms underlying plant proline metabolism and stress response. Therefore these findings provide potential avenues for crop genetically modified breeding.

## Introduction

Delta-1-pyrroline-5-carboxylate synthase (P5CS) is a key enzyme in the conserved proline biosynthesis pathway. This enzyme catalyzes a two-step reaction that converts glutamate to L-glutamate-5-phosphate (G5P), and then to L-glutamate-5-semialdehyde (GSA). GSA will spontaneously cyclize and reach equilibrium with delta-1-pyrroline-5-carboxylate (P5C). The final product P5C is an essential precursor for proline biosynthesis^1^.

As a bifunctional enzyme, P5CS has two structural domains: the glutamate kinase (GK) domain and the glutamyl phosphate reductase (GPR) domain. In prokaryotes, GK and GPR are two separate enzymes. For example, in *E. coli*, GK and GPR are encoded by proB and proA genes, respectively^2,3^. These two genes belong to the same operon, in the order of “promoter-proB-proA”^4^. During evolution, there may be gene fusion events between proB and proA before last eukaryotic common ancestor (LECA). Nowadays, most eukaryotes have bifunctional P5CS^5^.

We have found that *Drosophila melanogaster* P5CS (dmP5CS) can form cytoophidium, and purified dmP5CS can also form filament in vitro^6^. In dmP5CS filaments, GK tetramers are assembled as the “core” of the filaments, while GPR dimers surround GK tetramers in a double helix pattern^7^. The ordered arrangement of GK and GPR in dmP5CS may be encouraged during evolution, as the intermediate product G5P is highly unstable^8^. Some studies have suggested that there are some interactions between prokaryotic GK and GPR, which may have significant implications for substrate channeling of G5P^9,10^. For dmP5CS, mutations that disrupt filamentation can also impair overall activity dramatically^7^. It has been assumed that dmP5CS filaments promote G5P channeling, but the detailed mechanism between filamentation and enzyme activity is still unclear.

We are not sure if filamentation of P5CS is a conserved phenomenon or a special case of a few species. Except for dmP5CS, the full-length structure of P5CS has not been reported. Investigating the P5CS structure in another major eukaryotic group, plants, may be a good starting point. Despite the lack of structural information, many studies have emphasized the importance of P5CS in plants stress resistance. Proline accumulation has been observed when plants facing different abiotic conditions^11–13^, and as the key enzyme in proline biosynthesis, P5CS is upregulated under water stress^14,15^, salt stress^14,16,17^, cold stress^17,18^, and heavy metal stress^19,20^. *Arabidopsis thaliana* has two P5CSs (atP5CS1 and atP5CS2) with different functions^14,16^. atP5CS2 is considered to be housekeeping and has high expression level in actively dividing cells^14^. Knocking out atP5CS2 can lead to embryo lethality and can be rescued by exogenous proline supplementation^14,21^. On the other hand, atP5CS1 expression can be induced by light and stresses^14,22^. The induced expression of atP5CS1 is important for proline accumulation under stress conditions. Knocking out P5CS1 will decrease proline levels under water and salt stresses, and reduce salt tolerance^14^.

Climate change is a global concern today. In this context, crop breeding focuses more on the ability to resist abiotic stress. Although the exact role of proline in stress resistance is still unclear^23^, many researchers have attempted to use P5CS for transgenic breeding. Overexpressing *Vigna aconitifolia* P5CS (vaP5CS) can increase drought or salt tolerance in transgenic tobacco^24^, rice^25,26^, wheat^27^, potato^28^, and chickpea^29^. vaP5CS has a moderate feedback inhibition effect on proline (IC50 = 6 mM), and the F129A mutation of vaP5CS will remove the feedback inhibition effect (IC50 = 960 mM)^30^. Compared with tobacco expressing wild-type vaP5CS, transgenic tobacco expressing vaP5CS-F129A mutant protein accumulates more proline and is more tolerant to osmotic stress^31^. These studies show the potential value of P5CS or modified P5CS in crop breeding. However, little is known about the biochemical and structural properties of plant P5CS. Besides vaP5CS, only rice P5CS2 has been well studied at the biochemical level^32^. The structure of plant P5CS has not been reported yet. These gaps in plant P5CS studies limit the P5CS toolbox for transgenic breeding.

In this study, we choose atP5CS as the model to characterize the structure of plant P5CS. We find that atP5CS2 can also form filament, although in a different manner from dmP5CS. Stable interface between dmP5CS GPR domains is not found atP5CS2, and atP5CS2 GK domain is sufficient for filament formation. Site-directed mutagenesis of key residues on the helical interface of atP5CS2 disrupts atP5CS2 filament and impair the activity of atP5CS2. Interestingly, this mutation reduces the G5P utilization efficiency of G5P. We report the first structure of plant P5CS, showing that filamentation of P5CS is conserved in distinct eukaryotic organisms. Enzyme activity assays also indicate that there is substrate channeling between the GK domain and the GPR domain, and filamentation of P5CS is important for this channeling. The filament-based substrate channeling of atP5CS2 provides a new model for the intermediate transfer in multi-domain enzymes.

## Results

### The helical interface of P5CS is not conserved in plants

We have constructed a global landscapeof P5CS evolution. As previously pointed out, the bifunctional P5CS in eukaryotes evolved from the fusion of GK and GPR^5,33^. The GK and GPR domains in P5CS catalyze the same reaction as their ancestors. In the first step of proline biosynthesis, GK catalyzes the transfer of phosphorate from ATP to glutamate, producing G5P. Then, GPR catalyzes the reduction of G5P with NADPH to produce GSA. GSA will spontaneously cyclize in solvent and reach equilibrium with P5C (**Figure 1 A**). It should be noted that the intermediate product G5P is labile and will spontaneously cyclize to pyroglutamic acid (**Figure 1 A)**.

**Figure 1.**
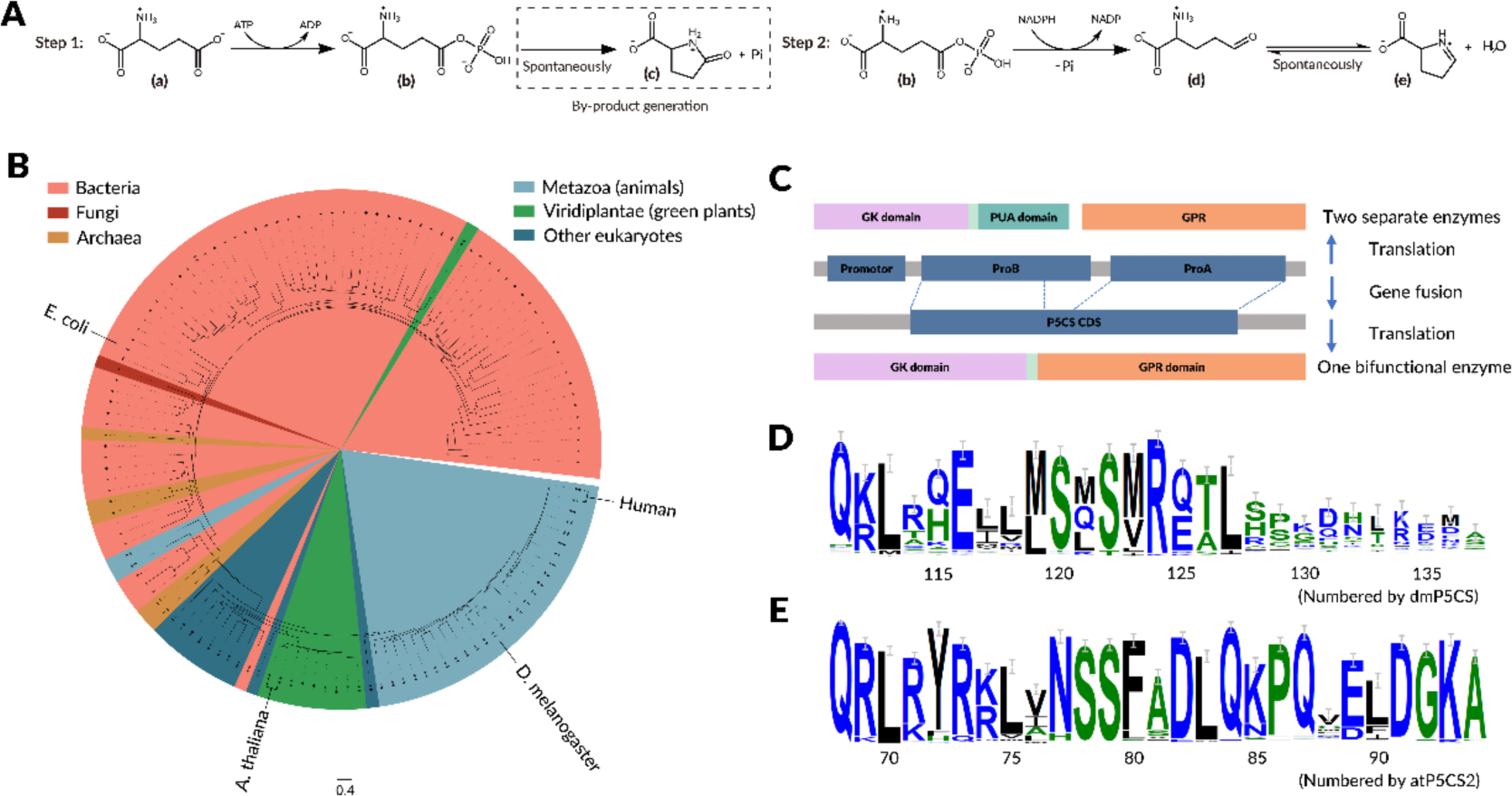
Domain organization and evolution of P5CS gene. (A) Two-step reaction catalyzed by P5CS. A possible side reaction is shown in the dashed box. a: L-glutamate. b: G5P. c: GSA. d: L-Pyroglutamic acid. e: Delta-1-pyrroline-5-carboxylate. (B) Phylogenetic tree of GK and P5CS proteins. Domain organization of these GK and P5CS is represented with marker at the extension of each clade, where “-” represent GK without PUA domain, “+” represent GK with PUA domain, and “++” represent P5CS. (C) Scheme of the fusion of GK and GPR during evolution. (D) Part of sequence logo of aligned animal P5CS. The reference residue number at the bottom is numbered based on dmP5CS. E116 and R124 are conserved in animal P5CS. (E) Part of sequence logo of aligned plant P5CS. The reference residue number at the bottom is numbered based on atP5CS2.

We visualized the evolution of P5CS by reconstructing phylogenetic trees including GK and P5CS. We downloaded protein sequences from the UniProt database, including all P5CS sequences and only the reviewed (Swiss-Prot) GK sequences. Only UniRef50 representative sequences were used to reduce data size. Finally, a phylogenetic tree was constructed using 77 GK clusters and 54 P5CS clusters (**Figure 1 B**). Most prokaryotes do not have P5CS, and some rare cases can be explained by horizontal gene transfer (HGT). Prokaryotic GK proteins typically have additional pseudouridine synthase and archaeosine transglycosylase (PUA) domains, which are lost in P5CS (**Figure 1 C**). The functions of these domains are not yet clear, with 17 out of 77 analyzed GK clusters lacking PUA domains (**Figure 1 B**).

Interestingly, all eukaryotes except fungi have P5CS. This is a bit strange because fungi have a relatively close relationship with animals, and they are classified into a group called Opisthokonta, which is separated from other eukaryotes. From the phylogenetic tree perspective, all available fungal GKs are grouped into a UniRef50 cluster. This clade is located in other prokaryotes GKs, and the closest sisters group belongs to cyanobacteria (**Figure S1**). We have not established a phylogenetic tree for GPR and P5CS, but according to the previous results, the closest fungal sisters group is also cyanobacteria^5^. Based on these results, it seems that after the separation of fungi and animals, the common ancestor of fungi lost P5CS and regained GK and GPR through HGT.

In this phylogenetic tree, plants and animals are clearly separated into two clusters. To investigate the detailed differences between plant P5CS and animal P5CS, we aligned all plant P5CS and animal P5CS from UniProt, respectively. From the structure of dmP5CS, we know that dmP5CS has two important filament assembly interfaces. The first one is the “hook” in the GK domain. The two necessary residues (from residue 105 to 127) E116 and R124 on the hook interface of dmP5CS are highly conserved in animal P5CS (**Figure 1 D**). In plant, the P5CS hooks (residues 62 to 84 in atP5CS2) correspond mainly to R and A (**Figure 1 E**). Loops following the plant hooks are shorter and more conserved than those in animal hooks (**Figure 1 D-E, Figure S2**). The second interface is located on the “contact loop” in the GPR domain. In dmP5CS, the “FGPP” motif in the middle of the loopis necessary for filament formation. This motif is conserved as “FXPX” (X refers to any amino acids) in animal P5CS (**Figure S3**). The same loop in plant P5CS is two residues shorter at the exact position of the motif (**Figure S2**). No similar motifs are found in plants (**Figure S3**). These sequence alignment results reveal significant differences between P5CS filament assembly interfaces in plants and animals.

### Substrates promote atP5CS2 filament formation

We choose atP5CS as the model to investigate the structure and function of plant P5CS. The two P5CSs in *Arabidopsis thaliana*, atP5CS1 and atP5CS2, share 89% identity in protein sequence. However, when recombinant proteins were expressed in *E. coli*, it was more difficult to get active atP5CS1 than atP5CS2. We focused on atP5CS2 for further experiment. During purification, atP5CS2 showed a wide range of molecular weight in size exclusion chromatography (**Figure S4**). Oligomerization state of atP5CS2 was checked with negative staining EM. At APO state (with 20 mM Tris-HCl, 150 mM NaCl, 10 mM MgCl2, pH = 8.0), no filament was observed. atP5CS2 filament will appear once there is ATP or glutamate in buffer (**Figure 2 A**). When both ATP and glutamate are present, which means GK domain is actively catalyzing, longer filaments were observed (**Figure 2 B-C**). Average length of atP5CS2 filament increased significantly when all substrates are added (mix state) compared with all other conditions. (**Figure 2 C**). With these negative staining and statistic data, we conclude atP5CS2 could form filament, and filamentation of atP5CS2 is regulated by its substrates.

**Figure 2.**
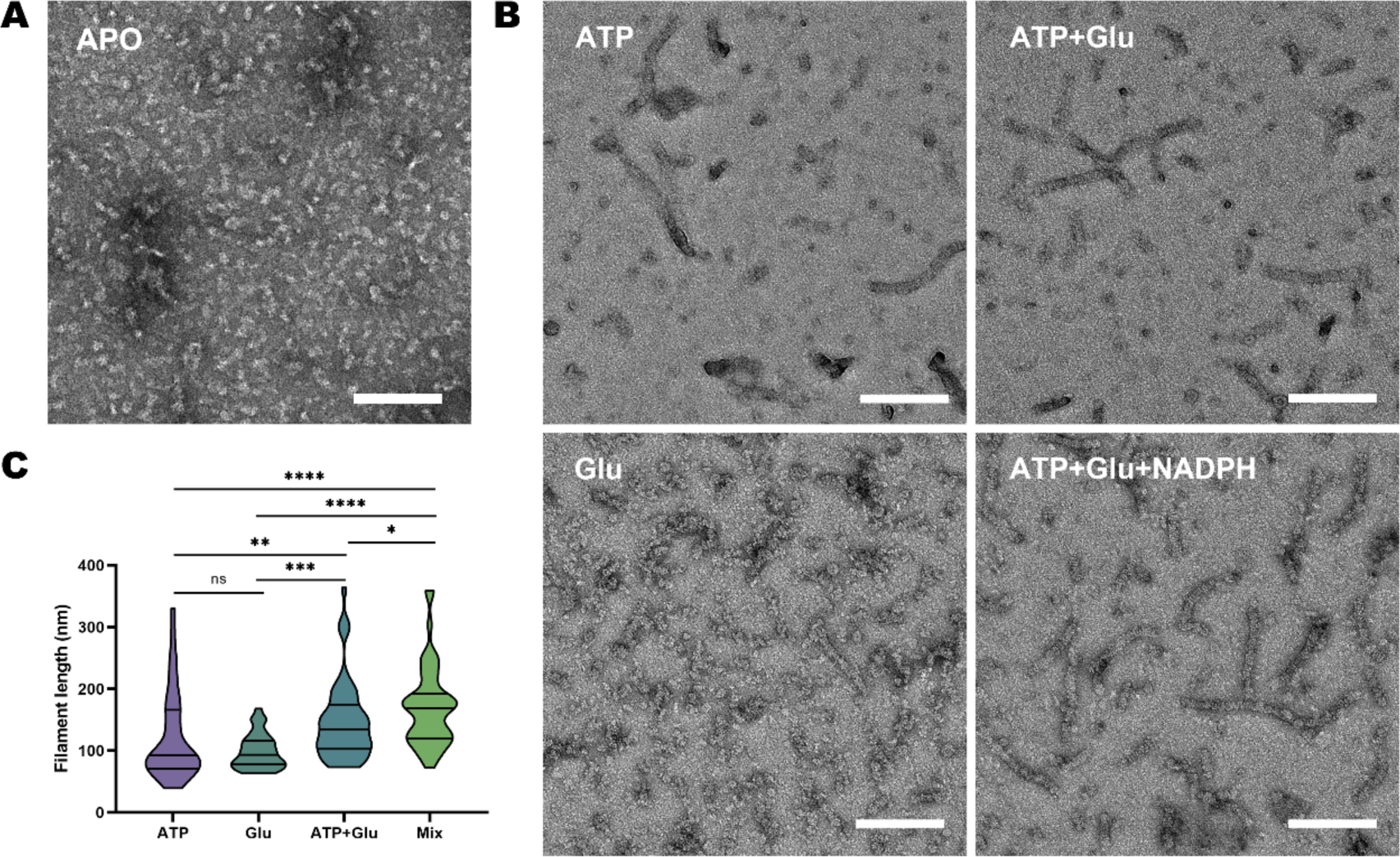
atP5CS forms active filament in vitro and responses to the ligand binding. (A) Negative staining EM micrographs of atP5CS2 at APO state. Scale bar: 160nm. (B) Negative staining EM micrographs of atP5CS2 incubated with different substrate combinations. Conditions are labeled at the top of each image. Scale bar: 160nm. (C) Statistical analysis of atP5CS2 filament length under different substrate conditions. Truncated violin plots show the medians and quartiles. ****P < 0.0001, ***P < 0.001, **P < 0.01, *P < 0.05, two-sided Mann–Whitney test.

Compared with dmP5CS, the regulation of atP5CS2 filament is different. For dmP5CS, the contribution of glutamate is greater than ATP^7^. For atP5CS2, both glutamate and ATP promote filament formation at similar degrees (**Figure 2 C**). For both P5CSs, mix state is the most suitable condition for filament formation.

### atP5CS2 tetramers are stackedperpendicularly to form filaments

To get more detailed understanding of the structural basis of atP5CS2 filament, we determined cryo-EM structure of mix state atP5CS2 (**Figure S5**). Using the single-particle analysis and focus refinement strategy, we obtained a map containing about 5 helical units of atP5CS2 (**Figure 3A**). Each helical unit is a tetramer, in which GK domain locates at the core as one tetramer, and GPR domain locates on both sides as two dimers. Helical units are arranged in a perpendicular way, and an obvious interface consists of hooks locates on the helical axis. Interestingly, in our map, there is no apparent interaction between adjacent GPRs, which is thought to be important for the activity of dmP5CS.

**Figure 3.**
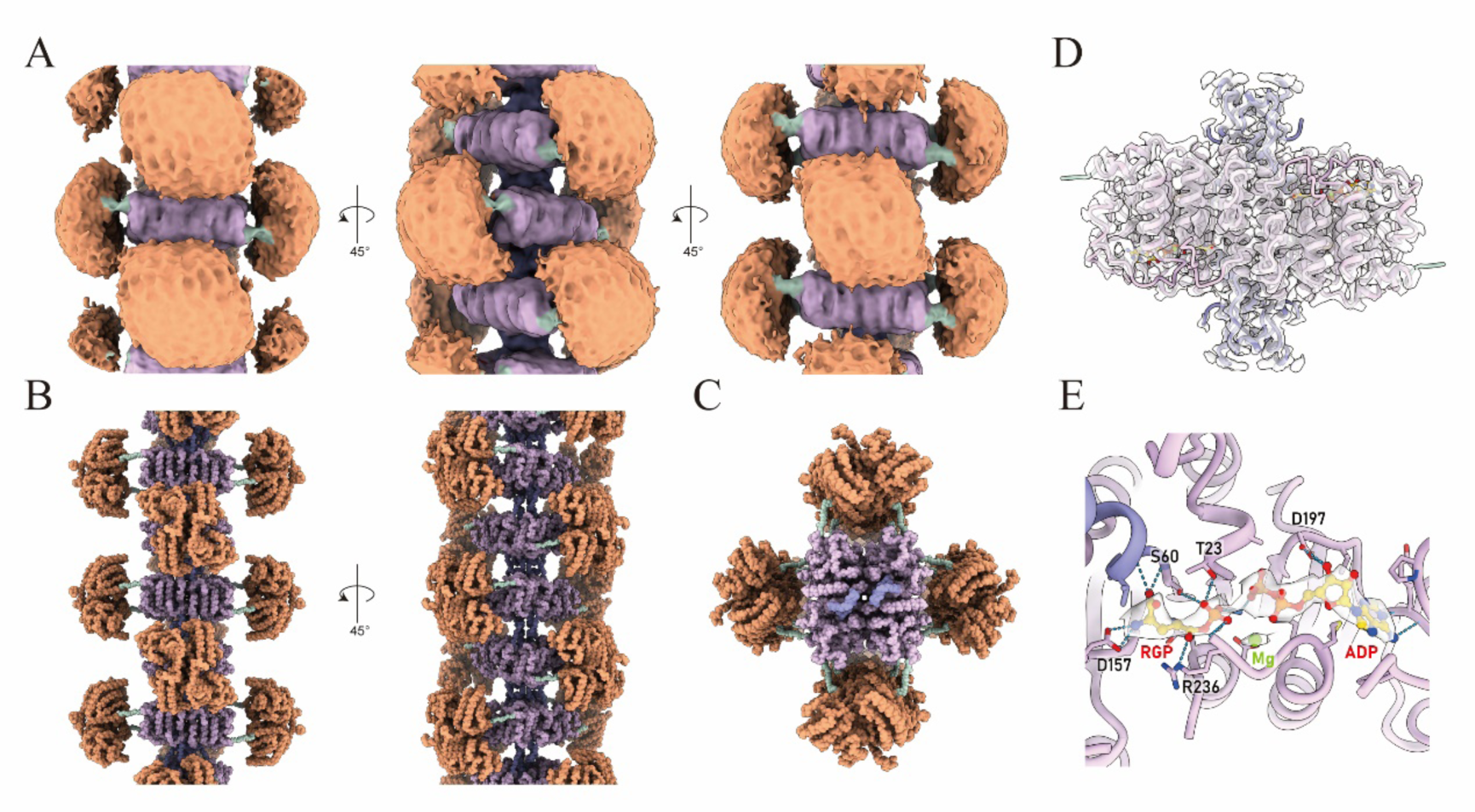
Overall structure of atP5CS filament. (A) Density map of atP5CS2 filament. GK domain, hook, GPR domain and linker of atP5CS2 filament are colored differently. (B) Proposed model for the atP5CS filament. This model was generated by using the predicted full-length model of atP5CS2 from AlphaFold2 and fitting it into our density map. (C) Zoom-in view of the GK domain. The map generated by focus-refinement is presented by a transparent surface. The “hook” region of GK domain is colored by purple and the others in pink. (D) The ligand binding pocket of GK domain. RGP and ADP, the products of GK domain are colored by gold. And their densities are displayed by the white transparent surface. Surrounding amino acids and potential interactions are indicated.

The only connection between GK domain and GPR domain is a flexible linker, and large motion is observed between GK tetramer and GPR dimer. In the overall reconstruction, only a blob can be seen at GPR domain. After focus refinement, the central GK tetramer could reach an average 3.3 Å resolution (**Figure 3D, S5-S6**), but no high-resolution reconstruction for GPR domain was made. By fitting the full-length model generated by AlphaFold2 into our map and manual adjustments, we built the model for atP5CS2 filament. The hook between adjacent helical units can be seen clearly in the side view of this model, and the whole filament appears as a cross in the top view due to the orthogonal assembly of atP5CS (**Figure 3B-C**).

In the focus refined GK tetramers, products binding mode was captured (**Figure 3E**). Compared to ADP, RGP (PDB code for G5P) is closer to the tetramer center, with the ammonia head of RGP stabilized by D157, and the carbohydrate group recognized by the peptide nitrogen on the backbone from the initial part of the hook. The gamma phosphate group of RGP has a specific interaction with surrounding amino acids, with the side chain of S60, T23, K20, and the backbone of R236 involved in stabilizing the gamma phosphate group of RGP.

Substrate ATP is stabilized by the ADP binding loop of atP5CS2. The triphosphate group of ATP is recognition and catalyzed by the highly conserved functional amino acids KKD of the kinase family. Besides, between the RGP and ADP, there is a magnesium ion that was previously observed in the model of dmP5CS. The conserved position for magnesium implies its potential function during the catalysis of glutamate kinase.

### The stable interface of atP5CS filament locates in the GK domain

The dmP5CS filament forms in a double helix pattern through the interaction of hooks and adjacent GPRs with high flexibility, which is considered to be the structural basis for its function. However, the GPR domain of atP5CS was found to be highly flexible based on 2D and 3D classification results. The representative classes of 2D classification show that the GK domain in the center of atP5CS remains in high resolution with clear details. A clear but small signal is resolved for the connection by the hook of adjacent helical units, whereas GPR on the side shows up in a blurred and light gray manner (**Figure 4 A**). These 2D classification results indicate the stable interface of atP5CS filament is located on the GK domain.

**Figure 4.**
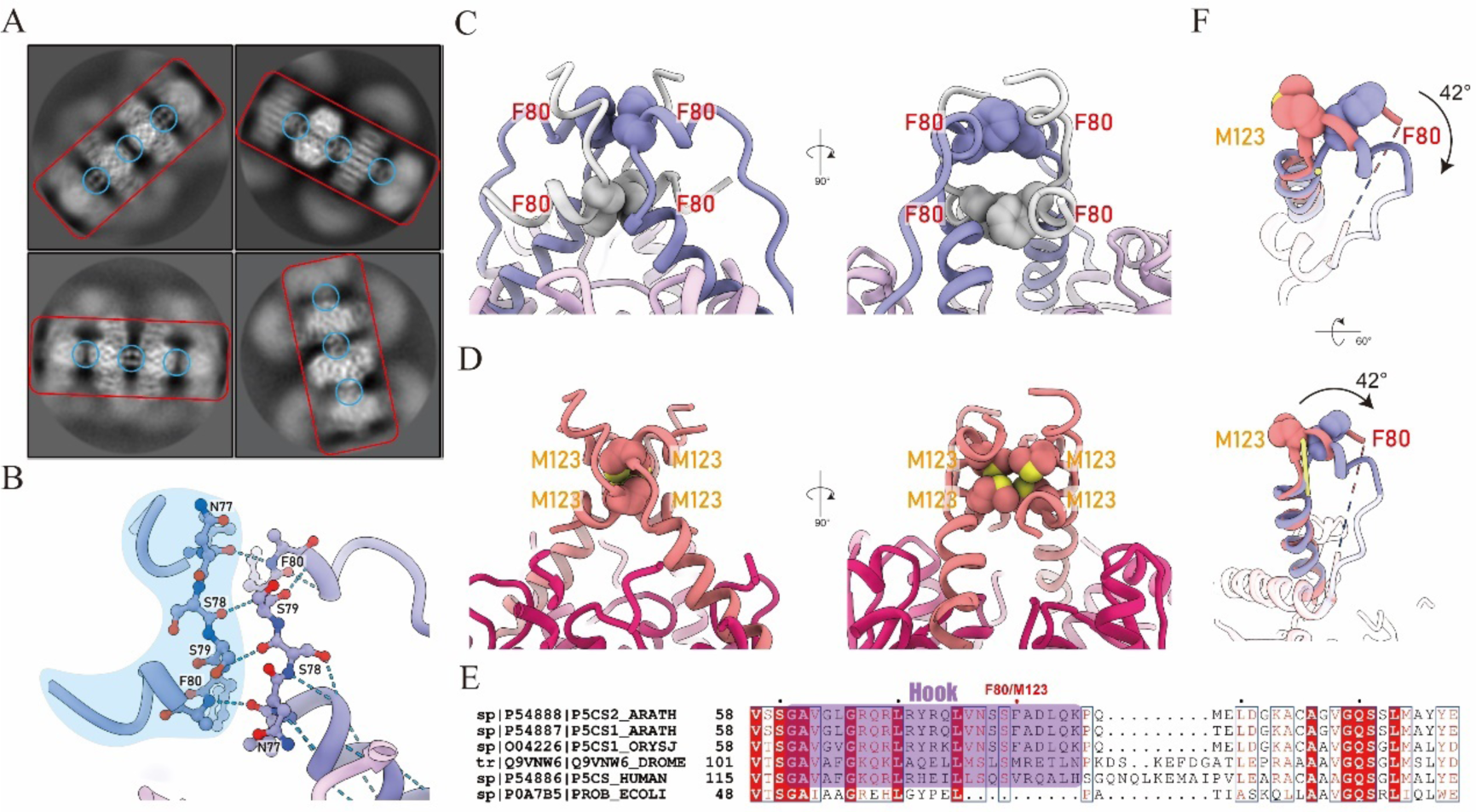
The stable interface of filament locates on the GK domain. (A) Representative 2D classification results of atP5CS filament. GK domain on the central of filament is indicated by the red box. Interfaces of GK domain composed of hook are marked by blue circle. (B) Hydrogen bond interaction inside a pair of hooks from different helical unit. For clearance, one of the hooks are highlighted by a blue background. (C) Front and side view of a hook tetramer in atP5CS filament. Hooks from one helical unit are colored by purple and the other by light grey. (D) Front and side view of a hook tetramer in dmP5CS filament. (E) Sequence alignment of P5CS from different species. The hook region is indicated by a purple box. (F) Comparison of hooks from atP5CS and dmP5CS filament. By aligning helix-3 of dmP5CS and atP5CS, the rotation of hook can be measured as 42°. The rotation axis is represented as a yellow stick in the figure.

We tried to capture potential transient stable GPR locations by 3D classification. After re-centered particles from GK tetramer to hook, the quality of 3D classification results was improved and density of GPR dimers became solid in many classes (**Figure S7**). None of these solid GPR dimers stays at the central line of atP5CS2 tetramer, i.e., the linkers between GK and GPR are not perpendicular to the helical axis. Comparing the locations of GPR dimers in different classes, we found the GPR dimer can rotate around the GK tetramer in a considerably wide range (**Figure S8 A-D**). Some transient interactions become possible during the rotation. Take class 11 as the example, the most possible interaction is between GPR dimer and a GK tetramer from adjacent helical unit. GK tetramer model and GPR dimer model were fitted into the map. The E347 in GPR and K131/135 in GK are quite close. A plausible interaction between E347 and K131/135 will be seen with proper rotamers (**Figure S8 E-F**). We also evaluated potential interactions between two GPR dimers. In class 11, two GPR dimers from adjacent atP5CS2 tetramers swing toward each other (**Figure S8 G**). Two pairs of Asn and Gln were found at the tip of GPR dimer. In the fitted models, the distances between side chains of N586 pair and Q557 pair are more than 10 Å (**Figure S8 H**), which is too far for direct interactions.

Combining the consensus map and classification results, we only find stable helical interface on GK domain. Some potential transient interactions on GPR domain are also revealed by 3D classification, and these interactions are all between two adjacent helical units (**Figure S8**). The stable interfaces built up by hooks maintain the filament and provide platform for other transient interactions.

### The helical interface of atP5CS2 assembles in a novel mode

The interface of the atP5CS filament is composed of four hooks from two adjacent helical units. Focusing on one side of the helical interface, four pairs of hydrogen bond form between the backbones of a pair of hooks from adjacent helical units. This interaction provides a direct connection between adjacent helical units and is conserved in the previously determined dmP5CS structures (**Figure 4 B**).

In the helical interface of atP5CS, the pi-pi interaction of F80 with another F80 from the same helical unit forms a bridge-like structure. Due to the symmetry, the other two protomers from adjacent helical units also form the same structure but in an inverted orientation. Together, the two bridges combine to form a lock structure that connects and stabilizes adjacent helical units along the helical axis (**Figure 4 C**).

Comparing the atP5CS and dmP5CS structures, it is observed that although the composition of the helical interface on the GK domain is still four hooks, the amino acid phenylalanine, which plays a key role in atP5CS, has been replaced by methionine (**Figure 4 D-E**). There is a distinct difference in the formation of the lock structure, as the four methionine gathers together and interact with each other, unlike the two bridges of a lock structure in atP5CS that maintain distance from one another.

Multiple sequence alignment reveals that the hook is highly conserved in both plants and animals. However, compared to animal p5cs, the following loop length of plant p5cs is noticeably shorter, and the amino acids in the relative position of F80 in plants are mainly phenylalanine, while in animals they are mainly methionine or valine (**Figure 4 E**).

The overall conformation of the hook structure is a helix-loop-helix and exhibits a similar angle in both dmP5CS and atP5CS. However, when aligning the long helix of the hook from atP5CS to dmP5CS, the short helix of the hook shows a rotation around the long helix of approximately 42 degrees (**Figure 4 F**). This results in M123 and F80 being located in different positions, and the change in the angle of the hook provides a structural basis for the difference in helical twist between dmP5CS and atP5CS.

### The GK domain is sufficient to form filaments

In dmP5CS, contact loop of GPR domain is necessary for filament formation. Mutation of key residues in dmP5CS contact loop will abolish filament formation. But in atP5CS2, whether GPR domain is necessary for filament formation is not clear. To explore this question, we expressed and purified truncated atP5CS2 protein consisting of residues 1-290 (referred to as atP5CS2-GK).

Similar to full length atP5CS2, atP5CS2-GK formed filaments quickly after incubated with substrates (**Figure 5 A**). Few filaments were observed for APO atP5CS2-GK (**Figure 5 A**). After incubated with glutamate, only a small number of tiny filaments could be found, and 3 minutes incubation was enough for longest filament formation. On the other hand, ATP would dramatically stimulate filament formation of atP5CS2-GK (**Figure 5 B**). When ATP was present in buffer, average length of filament would increase with incubation time, and maximum length was already reached after 7 minutes incubation (**Figure 5 B**).

**Figure 5.**
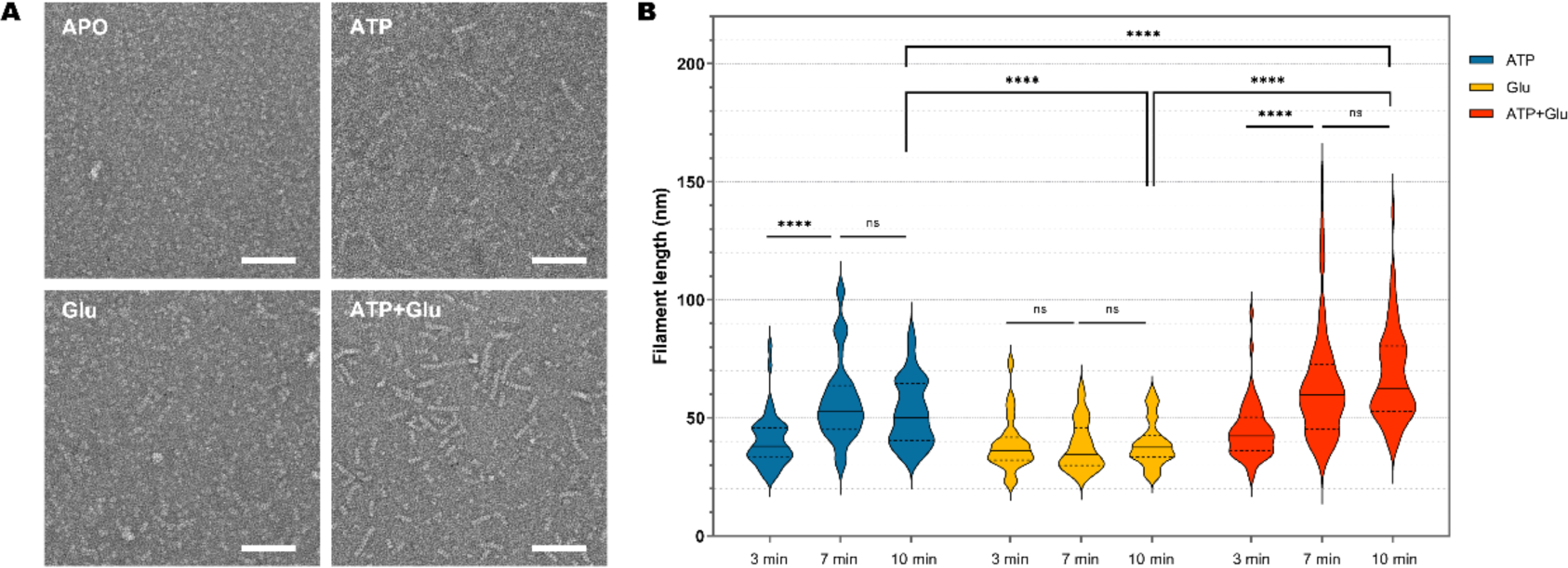
GK domain is sufficient to form filament. (A) Negative stain electron microscopy micrographs of atP5CS2 GK domain incubated with different substrates. Scale bar: 100 nm. (B) Statistical analysis of atP5CS2 filament length under different conditions. Violin plots show the medians (solid lines) and quartiles (dashed lines). ****P < 0.0001, two-sided Mann–Whitney test.

The longest filament of atP5CS2-GK appeared after 7 minutes incubation with both ATP and glutamate (**Figure 5 B**). We tested the activity of atP5CS2-GK using Pi assay. When incubated with ATP and glutamate, Pi increased linearly with incubation time, indicating this truncated protein is enzymatically active (**Figure S9**). So, like WT atP5CS2, atP5CS2-GK forms longest filament at reaction state.

### The destruction of atP5CS2 filament impairs catalysis

We use site mutagenesis to study the function of atP5CS2 filament. atP5CS2-F80A mutant was constructed according to structural analysis. At APO state, no filament was observed for both WT and F80A atP5CS2 (**Figure 6 A**). But after incubated with all substrates, WT and mutant atP5CS2 showed conspicuous difference (**Figure 6 A**). F80A mutation blocked the filament formation of atP5CS2. We measured the overall activity of WT and F80A atP5CS2 by decrease of absorbance at 340 nm (**Figure 6 B**). The decrease of A340 showed good linearity within 10 minutes. Vmax of F80A mutant is 7.07 μmol of NADPH min^−1^ μmol^−1^, only 30% of WT which is 23.28 μmol of NADPH min^−1^ μmol^−1^ (**Table 2**).

**Figure 6.**
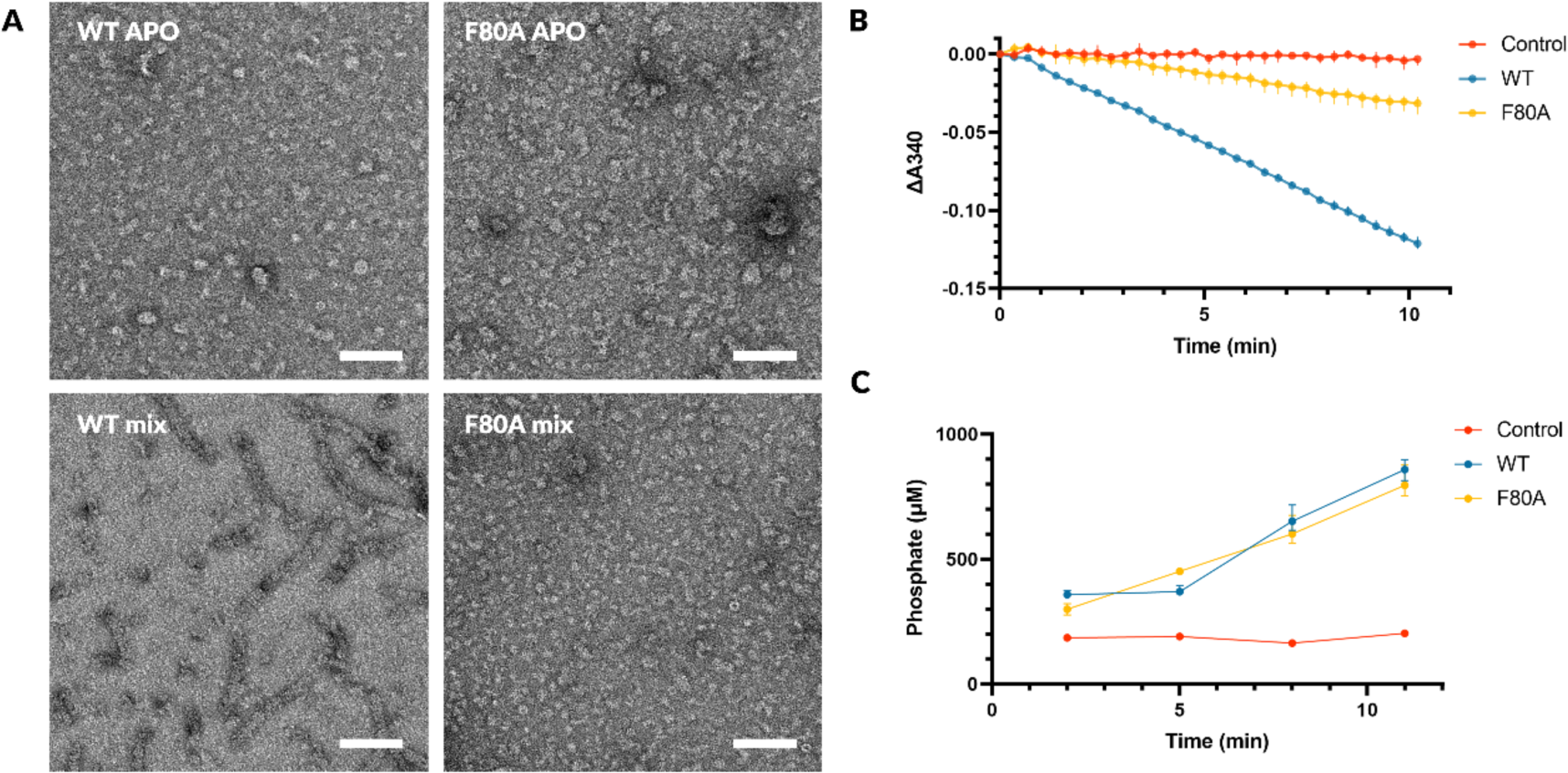
F80 is essential for the filamentation and function of atP5CS2. (A) Negative staining results of atP5CS2-WT and atP5CS2-F80A. Scale bar: 100 nm. (B) GPR activity of atP5CS2-WT and atP5CS2-F80A, measured by the change of A340. Error bars show the range of data (n = 3). (C) GK activity of atP5CS2-WT and atP5CS2-F80A, measured by phosphate generation. Error bars show the range of data (n = 3).

**Table 2.** Domain activity of atP5CS2-WT and atP5CS2-F80A.

|  | **WT** | **F80A** |
| --- | --- | --- |
| **GK activity <sup>a</sup>** | 396.24 | 364.27 |
| **GPR activity <sup>b</sup>** | 23.28 | 7.07 |
| **GPR/GK** | 5.9% | 1.9% |

a: $\mu\text{mol of Pi min}^{-1} \mu\text{mol}^{-1}$ . b: $\mu\text{mol of NADPH min}^{-1} \mu\text{mol}^{-1}$ .

We tried to find how F80A mutation affected the overall activity of atP5CS2. It has been assumed P5CS could transfer the intermediate G5P between two domains because G5P is labile^34^. Filamentation of P5CS may facilitates the channeling of G5P. To test this hypothesis, we measured the specific activity of GK domain in WT and F80A atP5CS2 by phosphate assay (**Figure 6 C**). Activity of atP5CS2-F80A GK domain is 364.27 μmol of Pi min^−1^ μmol^−1^, which is slightly slower than WT GK domain (396.24 μmol of Pi min^−1^ μmol^−1^). The results show that G5P generation rates are much higher than NADPH consumption rates in both WT and F80A enzymes, which is consistent with human P5CS^35^. Despite the similar specific activity of GK domain, the G5P utilization efficiency in WT enzyme is much higher than F80A enzyme (**Table 2**). Taken together, the F80A mutation impair the activity of atP5CS2 by reduce the efficiency of G5P channeling.

## Discussion

### Conservation of the P5CS filament

Some well-studied filamentous enzymes, like CTPS and PRPS, have shown that their filamentation is conserved across a wide range of species^36–41^. In previous studies, we found dmP5CS could form cytoophidia and filament, and the filament is critical for its activity^7^. But the conservation of P5CS filament was not clear. Here, we show that atP5CS2 also form filament, and this filament is regulated by ligands and is important for activity. With cryo-EM, we got the overall structure of atP5CS2 filament and reconstructed GK tetramer to high resolution. The stable helical interface was only found in GK domain, which is assembled by four hooks from two atP5CS2 tetramers. Multiple sequence alignment of plant P5CS shows that hook is highly conserved. On the other hand, hook of animal P5CS is also conserved in another manner. Thus, the filamentation of P5CS is likely to be conserved in both plants and animals.

### Divergence of P5CS filaments in plants and animals

The structure of atP5CS2 also reveals a novel filament assembly. In dmP5CS, two interfaces in GK and GPR are important for filamentation, and between them GPR is probably more important. While in atP5CS2, GK domain is sufficient to form filament. GPR domain of atP5CS2 may only have transient interactions with adjacent atP5CS2 tetramers. The primary interactions in atP5CS2 hook are also different from those in dmP5CS. Such dramatic changes on helical interfaces haven’t been reported in other filamentous enzymes. Multiple sequence alignments show that these properties are conserved in animals and plants, respectively. It is not clear whether these changes between plants and animals are due to special selection pressure, or just occasional mutations.

### Dynamics of atP5CS2 filaments and substrate channeling

Filamentation of atP5CS2 is important for its activity, especially for the channeling of G5P. However, no physical tunnel is observed. GK domain and GPR domain are connected with a flexible linker. No stable relative positions of GK and GPR exist in our mix state sample, while the flexibility of GPR seems to be important for its functions. We captured some possible transient interactions of GPR dimers by 3D classification. In some classes, GPR dimers can rotate a large angle so that the one GPR catalytic center is significantly closer to a GK catalytic center from an adjacent GK tetramer (**Figure S10**). This shorter distance may help G5P channeling through some unknown mechanisms.

### Catalysis and filamentation promote each other

Taken together, we propose a model connected the catalyzing and filamentation of atP5CS2. On one hand, catalyzing of atP5CS2 promotes filament formation. At APO state, some residues around hook may attract atP5CS2 tetramers together. After substrates binding or catalyzing, the atP5CS2 tetramers are able to overcome the energy barrier and form stable filaments. Repeating of the attraction and catalysis procedures elongate the atP5CS2 filament (**Figure 7A**). On the other hand, filamentation of atP5CS2 promotes substrate channeling of G5P. When stable filament forms through hooks on GK domain, a platform for other transient interactions is built. Some transient interactions will shorten the distance between activity center in GK domain and GPR domain, thus facilitates the channeling of G5P (**Figure 7B**).

**Figure 7.**
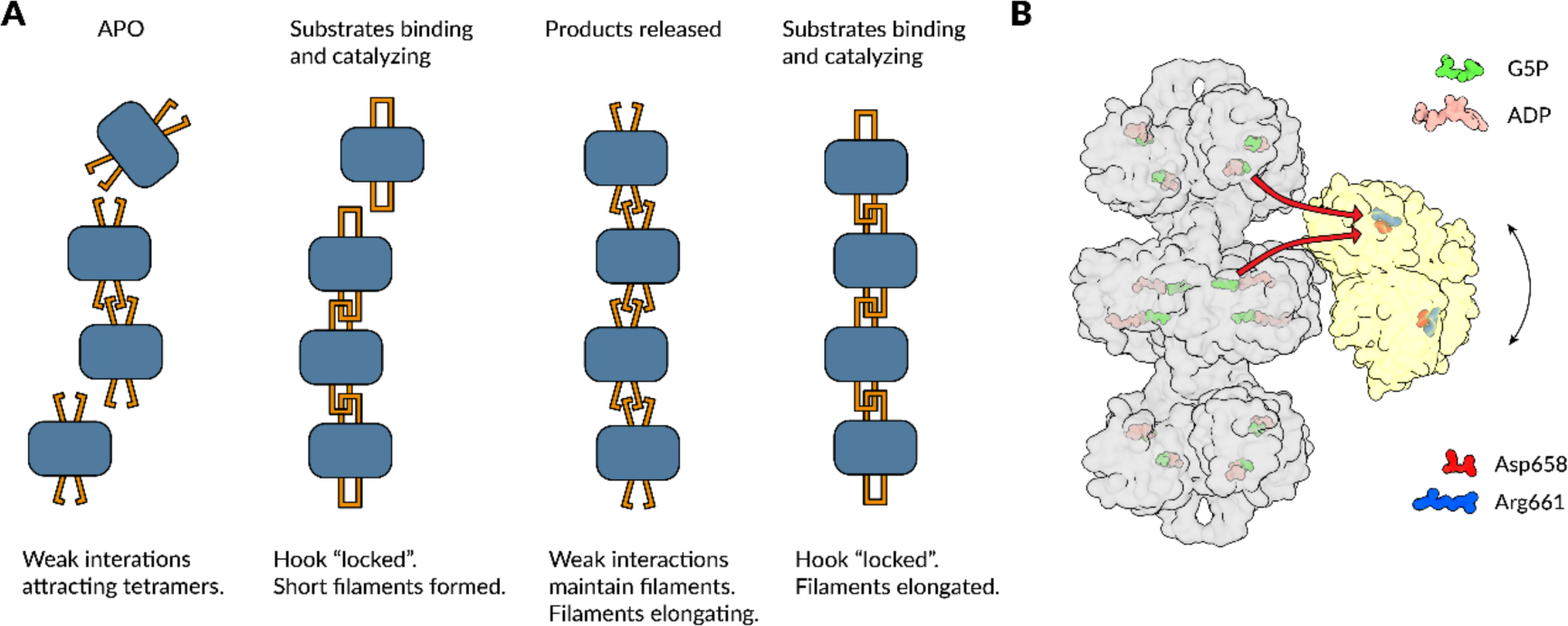
Model of atP5CS2 filament dynamics and contribution to substrates channeling. (A) Model of atP5CS2 filament dynamics. The blue rounded rectangles represent GK tetramers, and the orange sticks represent hooks. Two different states of hooks are shown in this figure: the open one (APO and products released states) and the locked one (substrates binding and catalyzing states). (B) Possible mechanisms of how filament contributes to substrates channeling. Three GK tetramers (gray) and one GPR dimer (yellow) are shown here. GPR dimer can rotates up and down around the central GK tetramer, and one farthest position of GPR dimer is shown here. G5P binding sites in GPR are inferred from dmP5CS model (PDB: 7WXI), two key amino acids involved in GPR binding, Asp658 and Arg661, are displayed. The red arrows show two possible G5P transfer paths.

## Data availability

The structure data accession codes are EMD-35901 and PDB-8J0F.

## Supporting information

Validation report

## Acknowledgments

We thank Binglian Zheng for providing the cDNA of *Arabidopsis thaliana*. EM data were collected at the ShanghaiTech Cryo-EM Imaging Facility. We thank the Molecular and Cell Biology Core Facility (MCBCF) at the School of Life Science and Technology, ShanghaiTech University and Shanghai Frontiers Science Center for Biomacromolecules and Precision Medicine for providing technical support. This work was supported by grants from: Ministry of Science and Technology of China (No. 2021YFA0804700), National Natural Science Foundation of China (No. 31771490), Shanghai Science and Technology Commission (No. 20JC1410500), UK Medical Research Council (grant nos. MC_UU_12021/3 and MC_U137788471) for grants to J.L.L.

## Author contributions

C.J.G initiated the project, cloned and established the expression system for target genes. T.Z performed the functional assays, the phylogenetic analysis, the cryo-EM data processing, model building and structure refinement. C.J.G and T.Z expressed and purified the proteins, prepared samples for EM studies, collected cryo-EM data, analyzed experiments, visualized results and wrote the manuscript. Q.L, X.Z and J.Z assisted the protein purification, functional assays or data analysis. J.L.L revised manuscript, obtained the funding and supervised the project.

## Competing interests

Authors declare that they have no competing interests.

## Methods

### atP5CS2 protein purification

atP5CS2 cDNA is a kind gift from Prof. Binglian Zheng (Fudan University). CDS of atP5CS2 gene was cloned into a modified pET28a vector, where a 6 × His-SUMO-tag was fused at the N terminus of atP5CS2. The expression vector was transformed into *E. coli* strain Rosetta (DE3). For protein expression, cells were first cultured at 37 ℃ to OD600 = 0.8-1.0, then protein expression was induced with 40 μM IPTG at 16 ℃ overnight. Cells were harvested by centrifugation at 4 ℃ and 4000 g, resuspended with lysis buffer (50 mM Tris-HCl pH 8.0, 500 mM NaCl, 10% glycerol, 20 mM imidazole, 1 mM PMSF, 5 mM β-mercaptoethanol, 5 mM benzamidine, 2 μg/ml leupeptin, and 2 μg/ml pepstatin). Then the cell suspension was homogenized with ultrasonic homogenizer (Scientz-IID). The lysate was centrifugated at 4 ℃, 18000 g for 45 min. Supernatant was incubated with Ni-NTA Agarose (QIAGEN) for 1 h and washed with 40 mM imidazole. Then proteins were eluted with elution buffer (50 mM Tris-HCl pH 8.0, 500 mM NaCl, 10% glycerol, 250 mM imidazole, 5 mM β-mercaptoethanol). Yeast ULP1 was used to remove SUMO-tag. Large amount of storage buffer (25 mM Tris-HCl pH 8.0 and 150 mM NaCl) was added together with ULP1 for optimal ULP1 activity. After incubated and concentrated at 4 ℃, proteins were further purified through Superose 6 Increase 10/300 GL (Cytiva) in storage buffer (25 mM Tris-HCl pH 8.0 and 150 mM NaCl). Peak fractions were collected, concentrated, snap-frozen with liquid nitrogen and stored at –80°C before use.

### atP5CS2 activity NADPH assay

The reaction buffer contains 25 mM Tris-HCl pH 8.0, 150 mM NaCl, 20 mM monosodium glutamate, 10 mM MgCl2, 5 mM ATP (Takara, sodium salt, pH 7.0), and 0.5 mM NADPH (Roche, tetrasodium salt). Reaction was started by adding 150 nM atP5CS2 (wild-type or mutant). Concentration of atP5CS2 was determined by BCA kit (Beyotime) using BSA as reference. The reaction was monitored at 25 ℃ in an SpectraMax i3 plate reader (Molecular Devices). Concentration of NADPH was converted from absorbance at 340 nm with molar extinction coefficients = 6.22 L·mol^−1^·cm^−1^. The path length was determined by PathCheck Sensor in SpectraMax i3 plate reader.

### atP5CS2 activity Pi assay

Phosphate is generated from two sources: spontaneous cyclization of G5P and reduction of G5P by NADPH. Taken together, the phosphate assay represents overall G5P generation rate. The reaction buffer is same as NADPH assay. 150 nM enzyme was added to start reaction. The reaction system was incubated at 25 ℃. Reaction was terminated at different times by adding ice-cold 4M perchloric acid to final concentration of 1 M. Then perchloric acid was neutralized and precipitated with 2M ice-cold KOH (final molar ratio perchloric acid : KOH = 1 : 1). The reaction mixture was centrifugated at 4 ℃, 13000 g for 15 min. Supernatant was used to determine Pi concentration with Malachite Green Phosphate Detection Kit (Beyotime).

### Negative staining

Proteins were incubated with different substrates at 25 ℃ before negative staining. All substrates used here had same concentration as in activity assay. Proteins were diluted to 2 – 4 μM. After incubation, protein samples were applied to plasma cleaned carbon-coated EM grids (400 mech, EMCN). The grids were then washed in ddH2O for twice and stained with 1% - 2% uranyl formate. Negative-stain EM grids were imaged with a Talos L120C microscope (FEI).

### Cryo-EM grid preparation and data collection

For cryo-EM, purified atP5CS2 was diluted to approximately 5 μM in reaction buffer. The sample was incubated for about 40 min on ice before vitrification. The sample was applied on H2/O2 plasma cleaned ANTcryo™ holy support film (Au300-R1.2/1.3) and then was immediately blotted for 3.0 s. The application and blot were repeated for twice. Then the grid was plunge-frozen in liquid ethane cooled by liquid nitrogen using Vitrobot (Thermo Fisher) at 8°C with 100% humidity. Images were collected on Titan Krios G3 (FEI) equipped with a K3 Summit direct electron detector (Gatan), operating in counting super-resolution mode at 300 kV with a total dose of 60 e^−^/Å^2^, subdivided into 50 frames in 2.8 s exposure using SerialEM^42^. The images were recorded at a nominal magnification of 22,500 × and a calibrated pixel size of 1.06 Å, with defocus ranging from 0.8 to 2.5 μm.

### Data processing

The image processing and reconstruction were performed with RELION 3.3^43,44^. We used MotionCor2^45^ and CTFFIND4^46^ via RELION GUI for pre-processing. Only micrographs with estimated resolution better than 5 Å were selected for further processing. We first use manual picking and 2D classification to generate 2D templates for atP5CS2 filament. Then atP5CS2 particles were picked with these templates. Particles were cleaned with several rounds of 2D classifications and two round of 3D classifications with C1 symmetry and then D2 symmetry. Finally, 223041 particles were used for 3D reconstruction with D2 symmetry and a filament with three helical units was acquired. Due to the flexibility of atP5CS2 filament, we performed an additional 3D refinement with a mask around GK tetramer to get better resolution. Then we used CTF refinement and Bayesian polishing^47^ to further improve the resolution. For detailed analysis on heterogeneity of GK monomer, we first applied D2 symmetry expansion on refined particles (after CTF refinement and Bayesian polishing). Then we performed local 3D auto-refine on GK tetramer with C1 symmetry. We first used 3D classification without alignment with mask on GK monomer and T = 4 to remove bad aligned particles. Then good particles were 3D local refined with a very soft mask on GK monomer. We used another round of 3D classification without alignment with mask on GK monomer and T = 20 to reveal dynamics on GK monomer. We got 4 best classes and each class was processed with local 3D refinement and sharpening. For all finally refined maps, local resolution estimation was performed using Relion’s own algorithm, and locally filtered maps were used for model building. To visualize the GPR domain, we re-extracted particles with shift along helical axis to re-center particles on the hook. Then after one round of 3D classification, some transient locations of GPR dimers can be visualized.

### Model building

The initial model of atP5CS2 GK domain was acquired through AlpahFold2^48^ database (https://alphafold.com/). Initial model was fitted into maps using UCSF Chimera^49^. Models were iteratively refined using a combination of Coot^50^ and phenix.real_space_refine in Phenix^51,52^. For GPR dimer, model was generated with AlphaFold2 Multimer^53^ on our local server with full-length atP5CS2 dimer as input. To analyze possible interactions of GPR dimers, rotamers of some residues in the fitted models were adjusted in UCSF Chimera using Dunbrack 2010^54^ rotamer library. UCSF ChimeraX^55^ was also used for visualization purpose.

### Evolutional analysis of P5CS

All P5CS protein sequences were downloaded from UniProt database. All multiple sequence alignments (MSA) used MAFFT with method E-INS-i^56^. For phylogenetic tree reconstruction, we downloaded all sequences which were annotated with EC number “2.7.2.11” (GK) or “2.7.2.11” and “1.2.1.41” (P5CS). Only reviewed GK sequences were used. The sequences were filtered with length (to remove too short or too long sequences) and re-mapped to UniRef50 reference sequences. After alignment, gaps were removed by trimAl^57^. Phylogenetic tree was built using IQTREE^58^ and model was auto selected^59^ (Q.pfam+I+I+R7 chosen according to BIC). Phylogenetic tree was visualized using FigTree^60^ and colored with Adobe illustrator. ESPript 3^61^ was used for MSA visualization. Sequence logos were generated using WebLogo 3^62^.

---

**Table 1.** Cryo-EM data collection, refinement, and validation statistics (in a separate Excel file)

## Supplementary information

**Figure S1.**
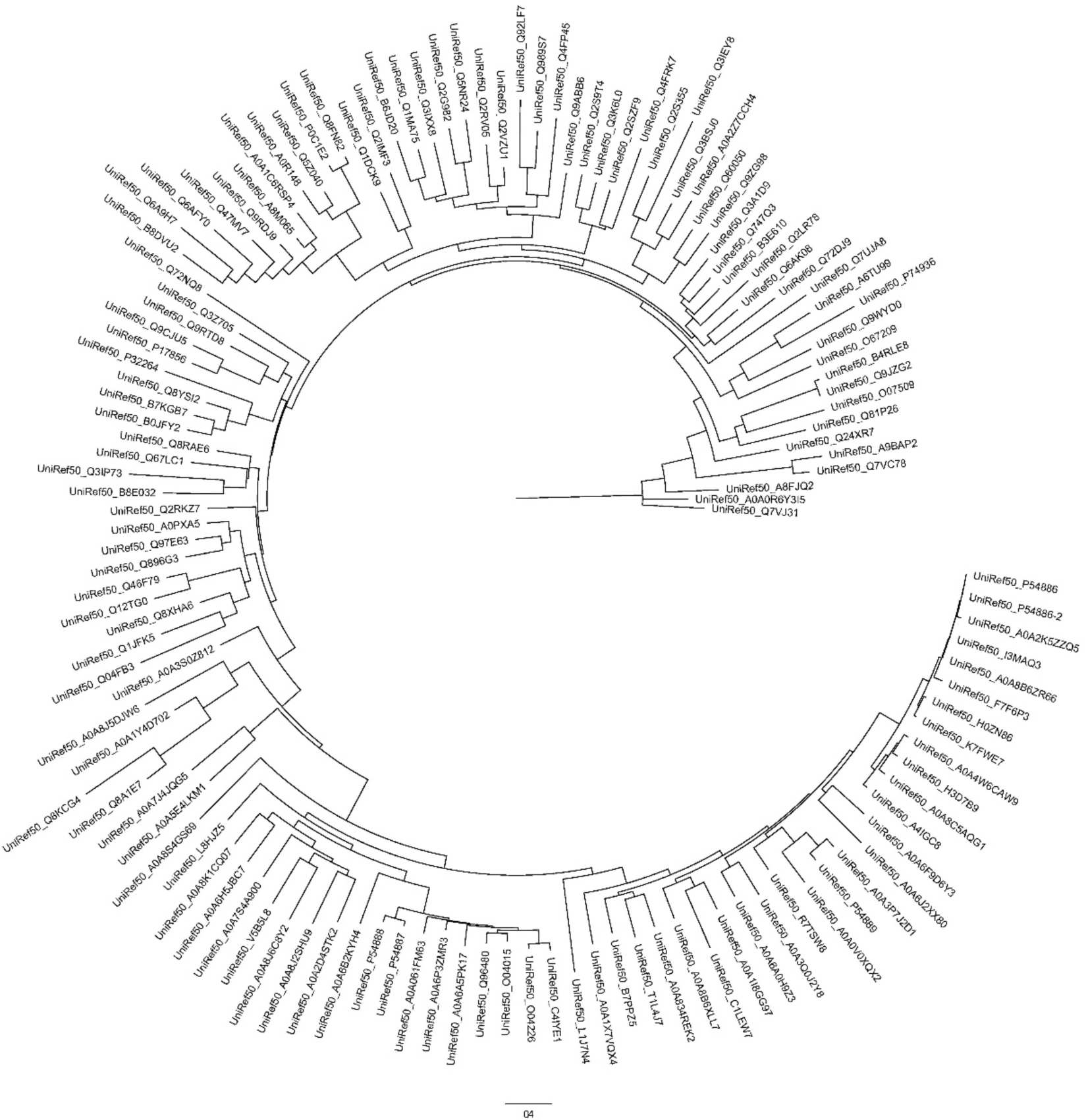
Original phylogenetic tree of GK and P5CS. UniRef50 representative sequences are used for multi sequence alignment and phylogenetic tree reconstruction. Name of each UniRef50 cluster is labeled at the tip of each clade. This original tree has the same orientation as the Figure 1B.

**Figure S2.**
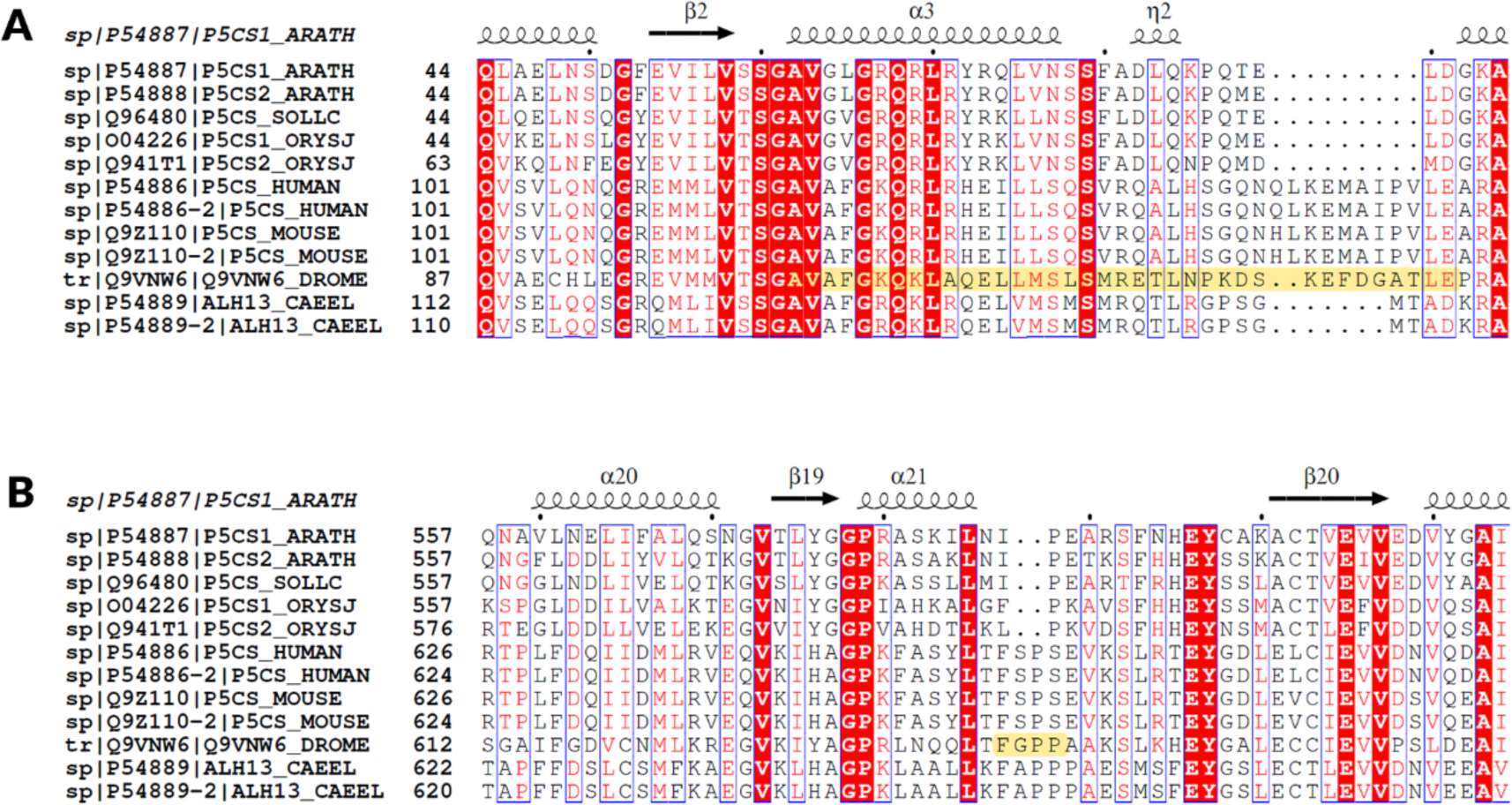
Alignment results of some representative P5CS sequences. (A) sequence alignment result around hook motif. Residues corresponding to hook of dmP5CS are highlighted. (B) sequence alignment result around contact loop. Residues corresponding to the conserved motif “FXPX” in dmP5CS are highlighted.

**Figure S3.**
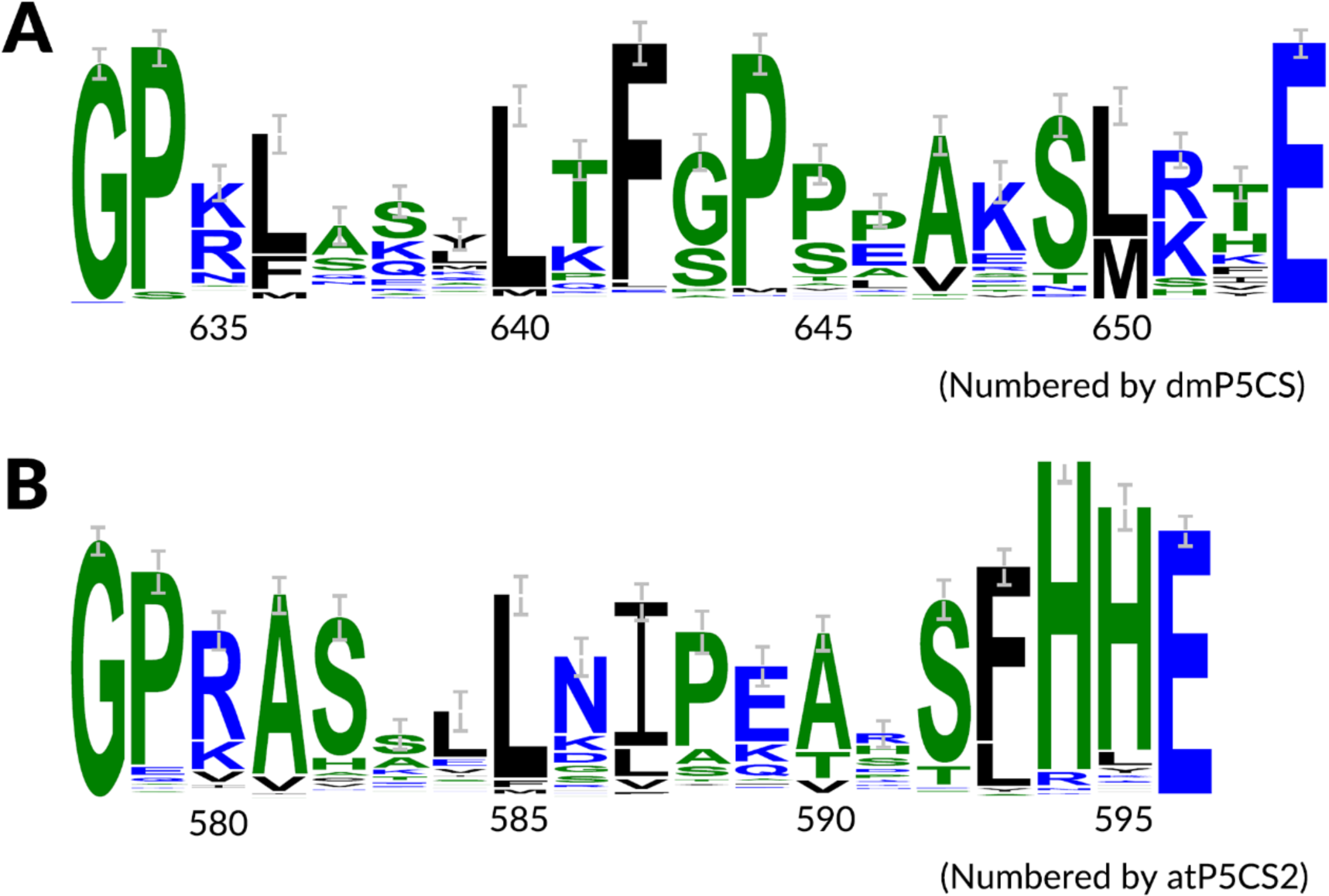
Part of sequence logo of aligned P5CS around contact loop. (A) Sequence logo of aligned animal P5CS. The reference residue number at the bottom is numbered based on dmP5CS. (B) Sequence logo of aligned plant P5CS. The reference residue number at the bottom is numbered based on atP5CS2. Plant P5CS are two residues shorter than animal P5CS in this region.

**Figure S4.**
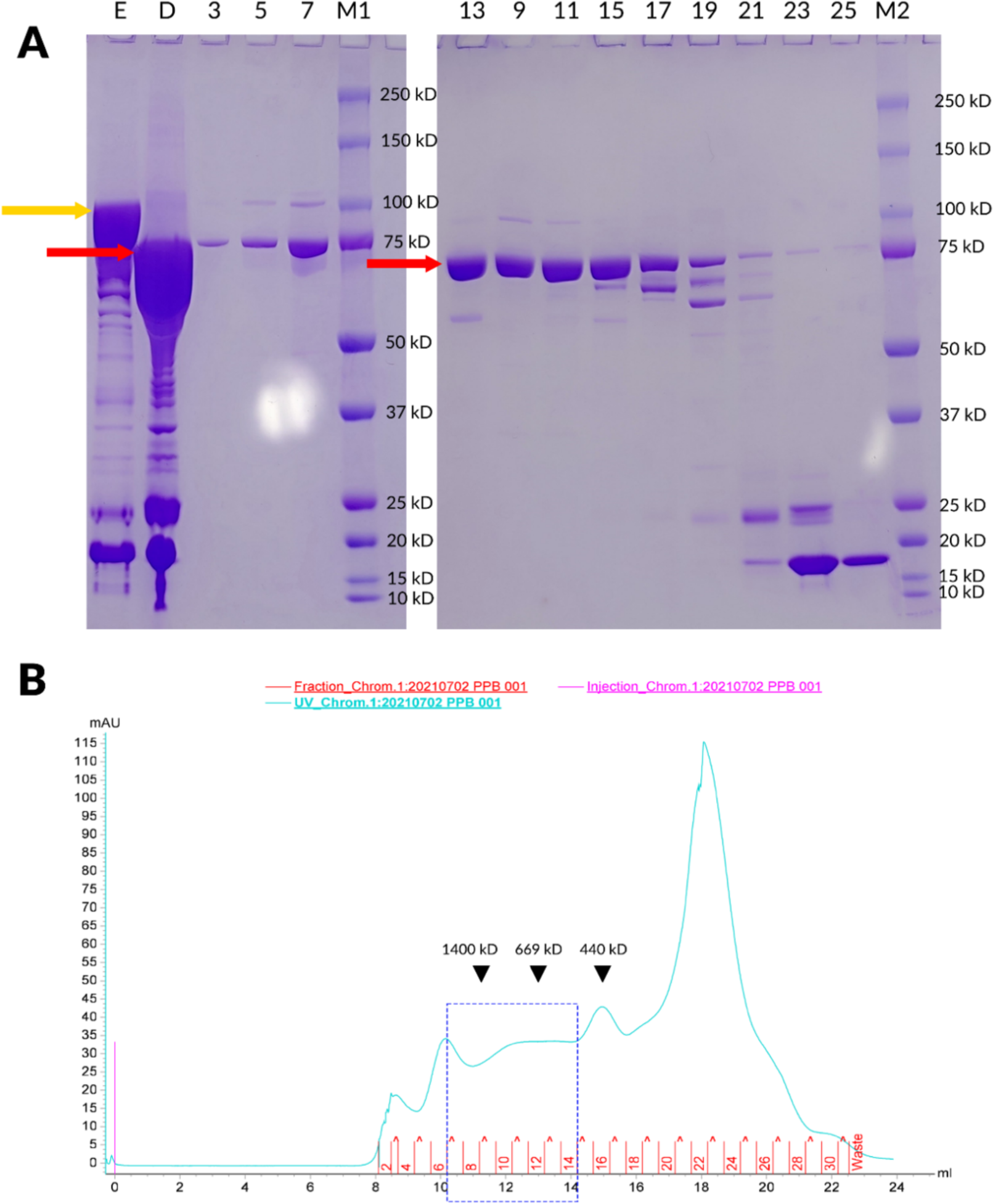
Qualification of atP5CS2 during one purification experiment. (A) SDS-PAGE of atP5CS2 at different purification stages. The band corresponding to SUMO-atP5CS2 fusion protein (about 92.6 kDa) is indicated with yellow arrow. The bands corresponding to atP5CS2 protein (about 78.9 kDa) are indicated with red arrows. Different fractions during purification are represented with characters on the top of panel A. “E”: Elution from Ni^2+^-NTA beads. “D”: Elution after ULP1 digestion. “M1” and “M2”: Standard protein markers. The corresponding molecular weights are labeled at the sides of the bands. “3” to “25”: Different fractions after purification with size exclusion chromatography (Superose™ 6 Increase 10/300 GL). (B) Elution profile from size exclusion chromatography. The fractions are numbered in red above X-axis. The fraction numbers are the same as those in panel A. In this experiment we collected fraction 7-14 (indicated as blue box) as final product. Reference molecular weights are labeled with black triangles.

**Figure S5.**
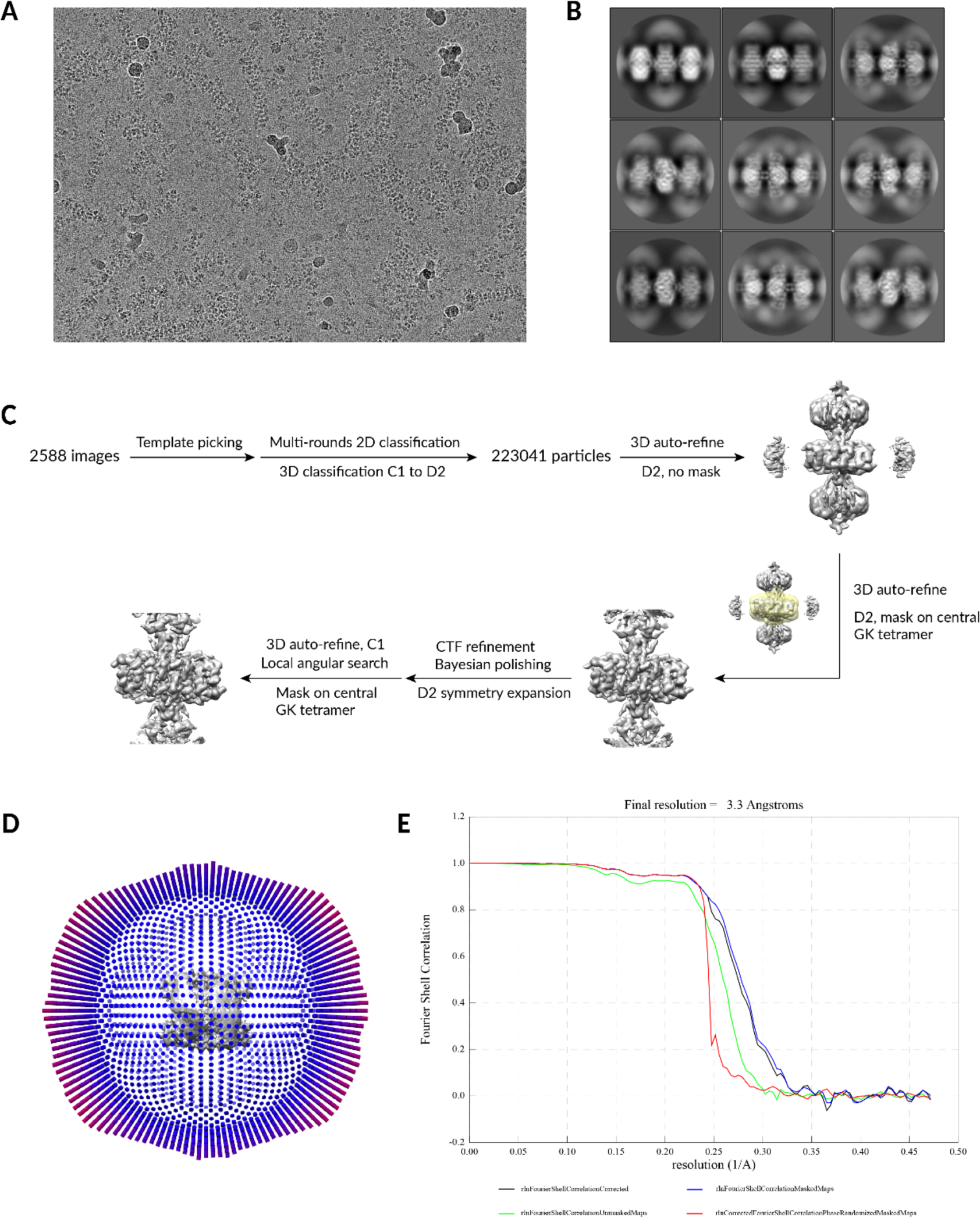
Data processing of atP5CS2. (A) Representative micrograph after motion correction. (B) Representative 2D classification results. (C) Workflow for reconstruction atP5CS2 filament and GK tetramer. (D) Angular distribution (top view) of final GK tetramer reconstruction. Particles had been D2 expanded. (E) FSC curves for focus refined GK tetramer.

**Figure S6.**
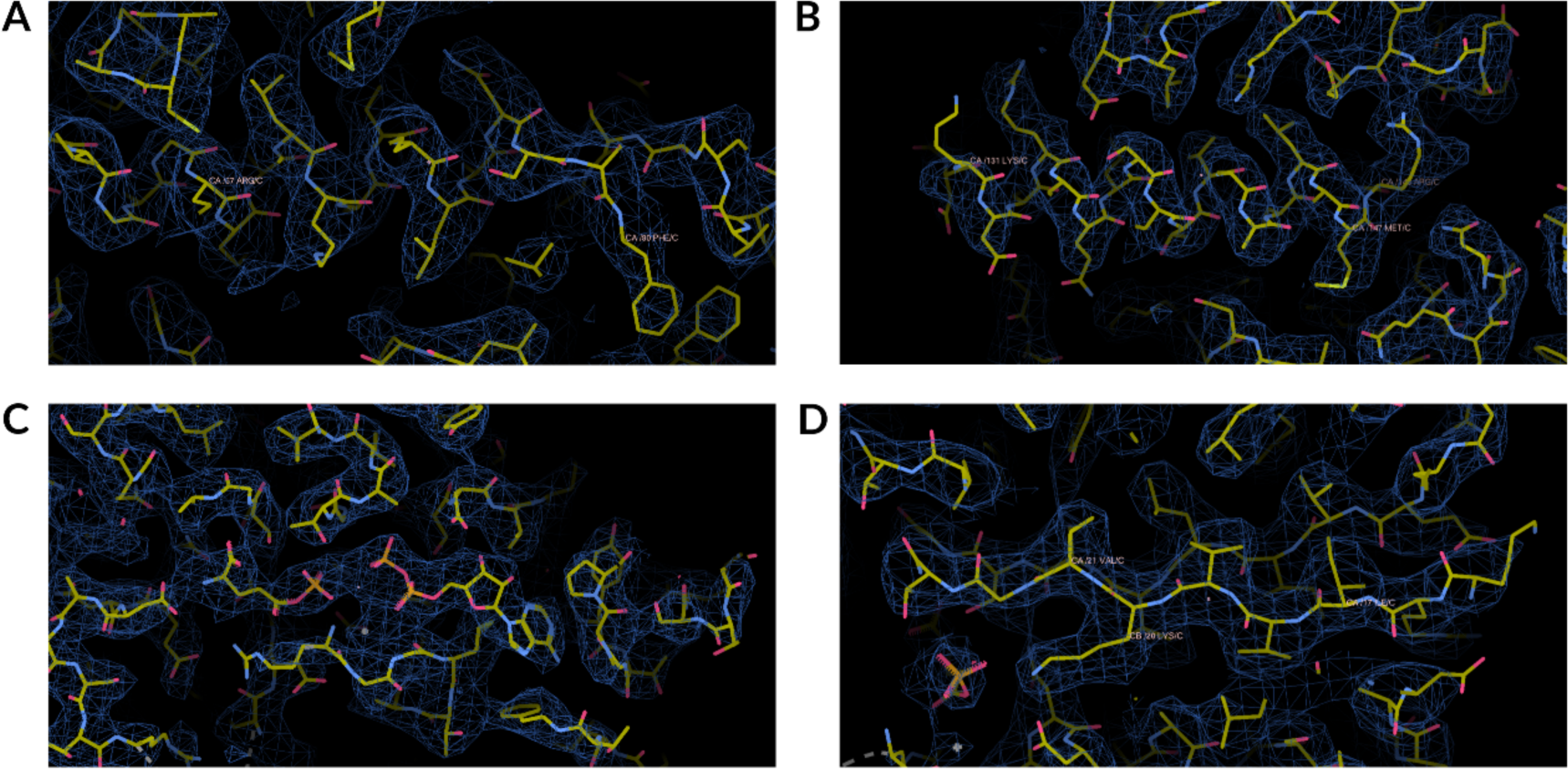
Representative density of focus refined GK tetramer. (A-D) Density around representative helixes, β-sheet, and ligands.

**Figure S7.**
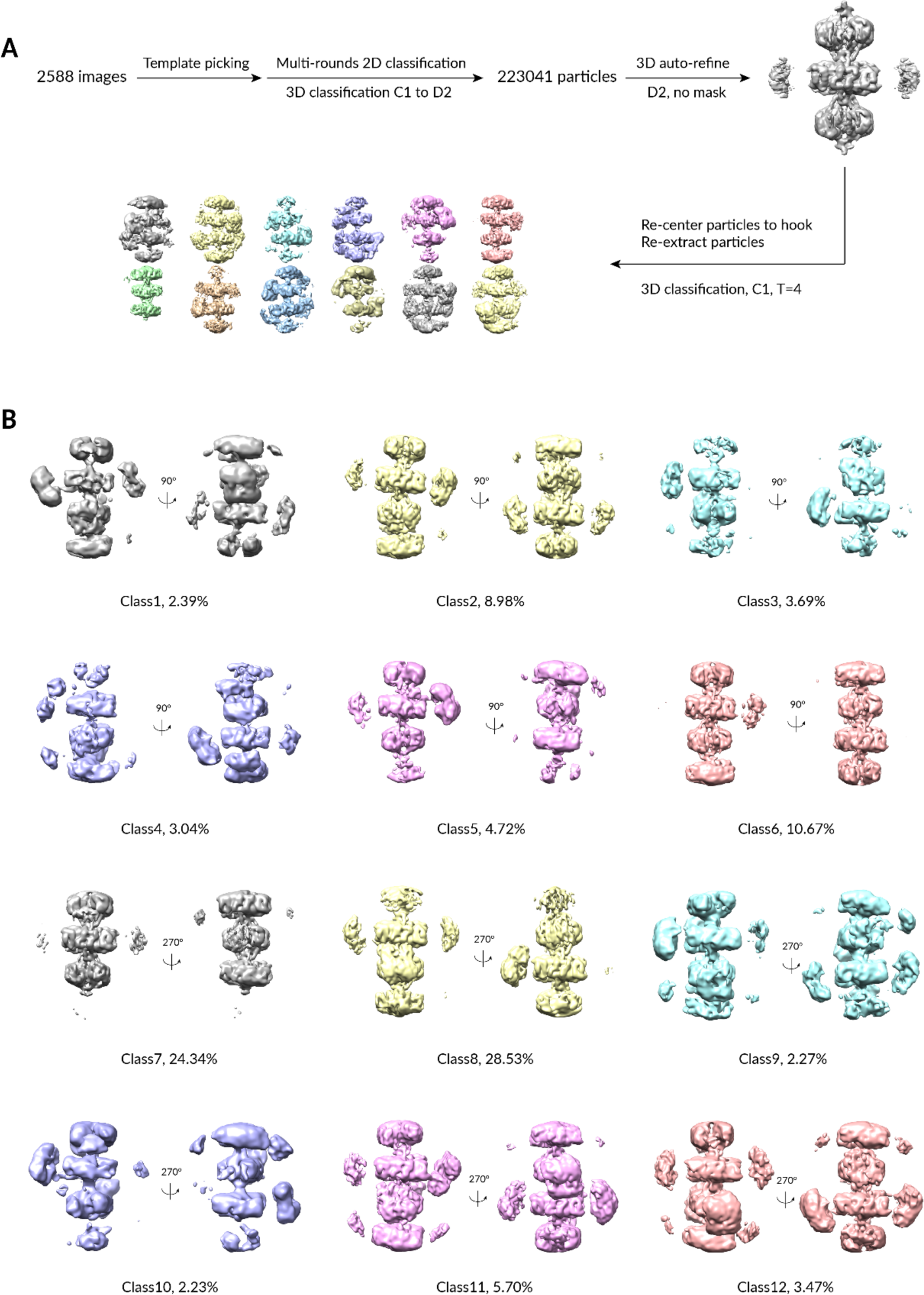
3D classification to visualize GPR. (A) Workflow for GPR 3D classification. After the first reconstruction for atP5CS2, particles were re-centered to hook and re-extracted. (B) Display of 12 classes. GPR can be visualized in most classes with very dynamic positions.

**Figure S8.**
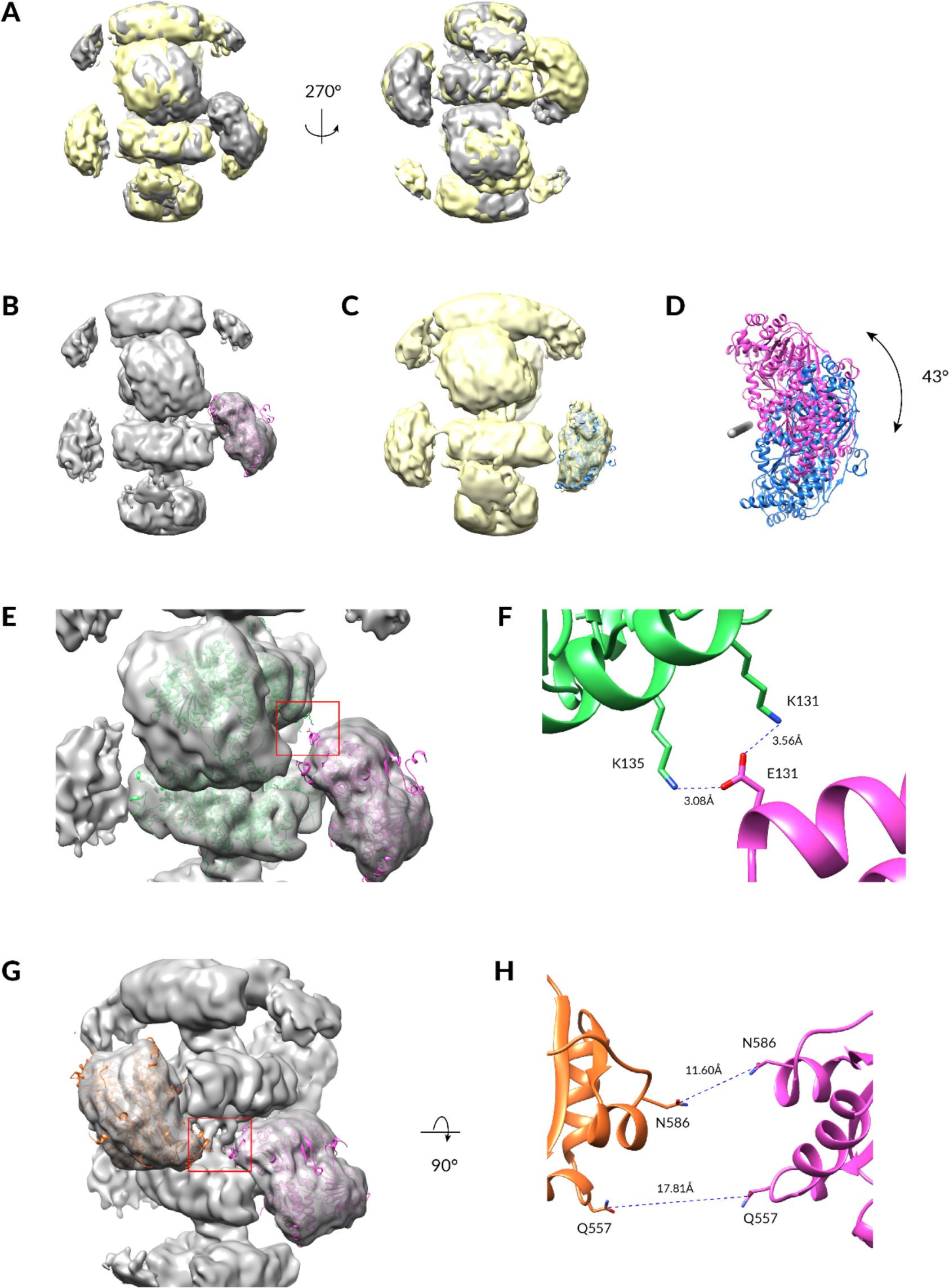
Dynamic positions of GPR dimers. (A) Superposition of class11 (gray) and class12 (yellow) maps. GPR dimers show different locations in these two classes. (B) AlphaFold2 model for GPR dimer is fitted into class 11 map. (C) AlphaFold2 model for GPR dimer is fitted into class 12 map. (D) Relative rotation between GPR dimers in class 11 (magenta) and class 12 (blue). The gray bar represents the rotation axis. (E) One GPR dimer model (AlphaFold2) and two GK tetramer models are fitted into class 11 map. The potential interaction region is label with red box. (F) Zoom in view of the red box in panel E. Rotamers of these displayed residues are manually adjusted in UCSF Chimera with Dunbrack 2010 library. (G) Two GPR dimer models (AlphaFold2) are fitted into class 11 map. The potential interaction region is label with red box. (H) Zoom in view of the red box in panel G (rotated by 90°). Rotamers of these displayed residues are manually adjusted in UCSF Chimera with Dunbrack 2010 library.

**Figure S9.**
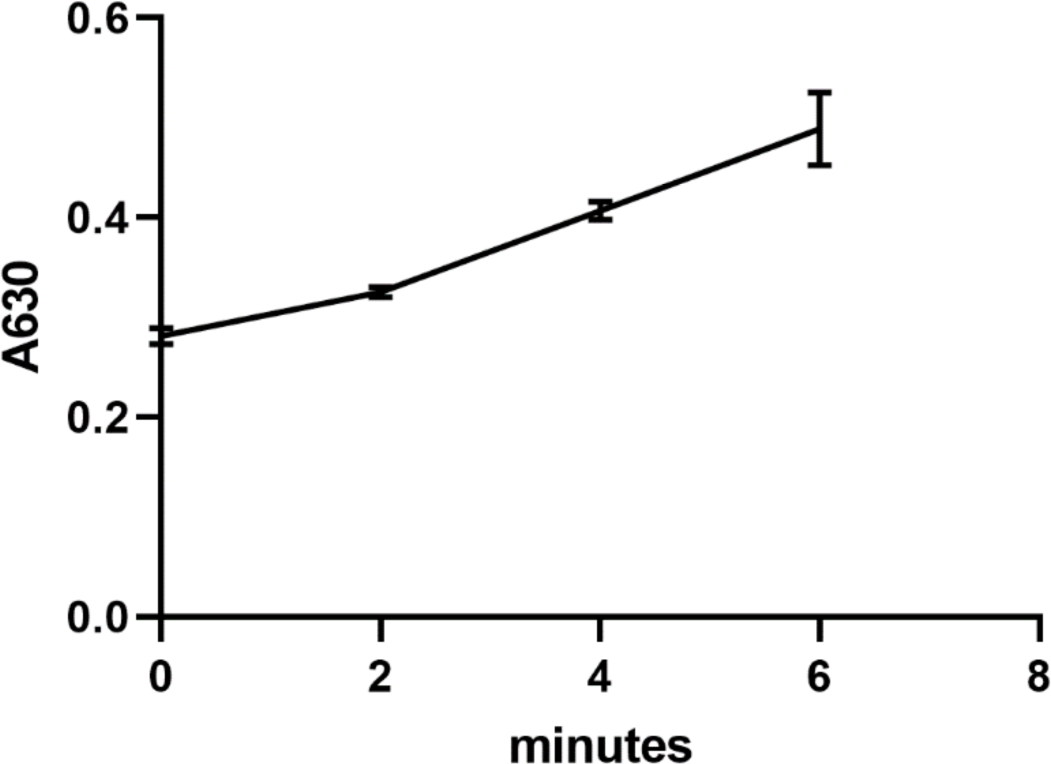
Activity of atP5CS2-GK. A630 is proportional to the Pi produced in reaction buffer. Error bar shows the range of data (n =3).

**Figure S10.**
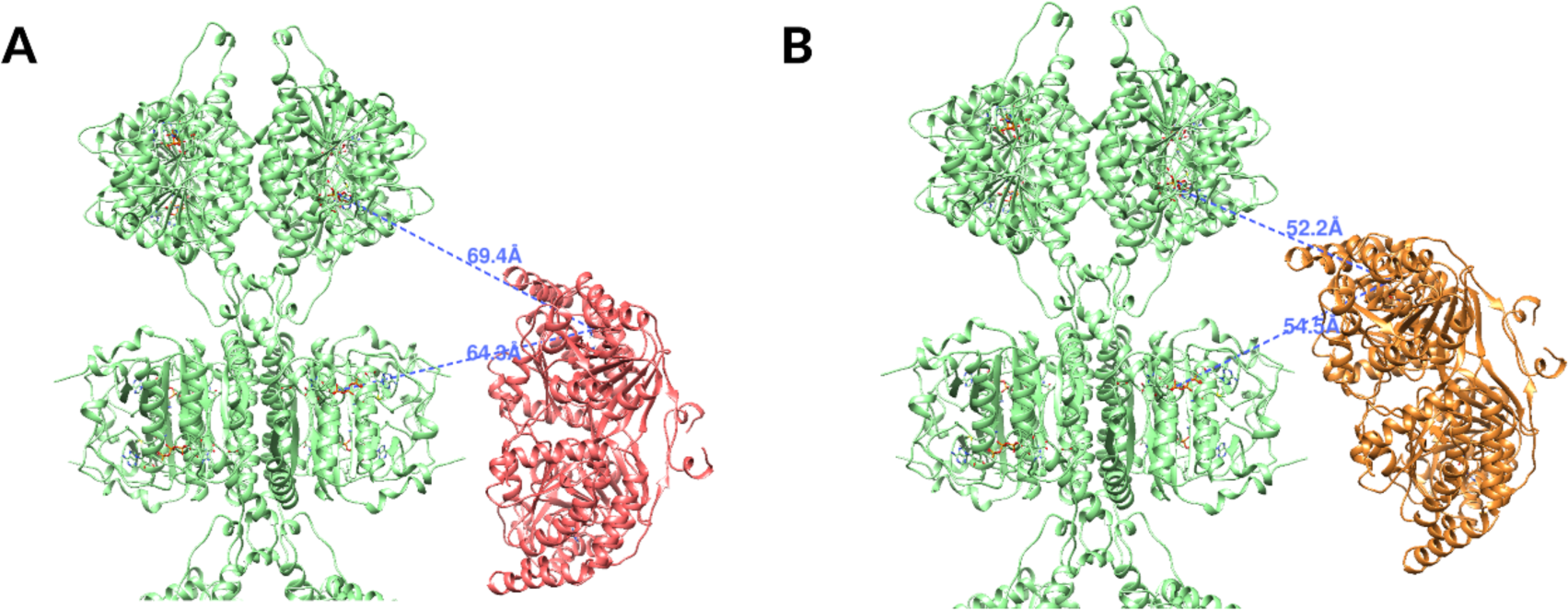
Comparison of distances between GK catalytic center and GPR catalytic center when GPR at different positions. (A) GPR dimer locates at the middle position. (B) GPR dimer rotates to the farthest position (class 11).

## Notes

### Competing Interest Statement

The authors have declared no competing interest.

