## Supplementary material for "Dynamic atP5CS2 Filament Facilitates Substrate Channeling": Validation report

### Full wwPDB EM Validation Report (i)

Apr 26, 2023 – 01:19 PM JST

| | | |
|---|---|---|
| PDB ID | : | 8J0F |
| EMDB ID | : | EMD-35901 |
| Title | : | GK tetramer with adjacent hooks at reaction state |
| Deposited on | : | 2023-04-10 |
| Resolution | : | 3.30 Å(reported) |

**This wwPDB validation report is for manuscript review**

This is a Full wwPDB EM Validation Report.

This report is produced by the wwPDB biocuration pipeline after annotation of the structure.

We welcome your comments at **
A user guide is available at
https://www.wwpdb.org/validation/2017/EMValidationReportHelp
with specific help available everywhere you see the (i) symbol.

The types of validation reports are described at
http://www.wwpdb.org/validation/2017/FAQs#types.

The following versions of software and data (see references (i)) were used in the production of this report:

| | | |
|---|---|---|
| EMDB validation analysis | : | 0.0.1.dev50 |
| Mogul | : | 1.8.5 (274361), CSD as541be (2020) |
| MolProbity | : | 4.02b-467 |
| buster-report | : | 1.1.7 (2018) |
| Percentile statistics | : | 20191225.v01 (using entries in the PDB archive December 25th 2019) |
| MapQ | : | 1.9.9 |
| Ideal geometry (proteins) | : | Engh & Huber (2001) |
| Ideal geometry (DNA, RNA) | : | Parkinson et al. (1996) |
| Validation Pipeline (wwPDB-VP) | : | 2.32.2 |

For Manuscript Review

### 1 Overall quality at a glance (i)

The following experimental techniques were used to determine the structure:
*ELECTRON MICROSCOPY*

The reported resolution of this entry is 3.30 Å.

Percentile scores (ranging between 0-100) for global validation metrics of the entry are shown in the following graphic. The table shows the number of entries on which the scores are based.

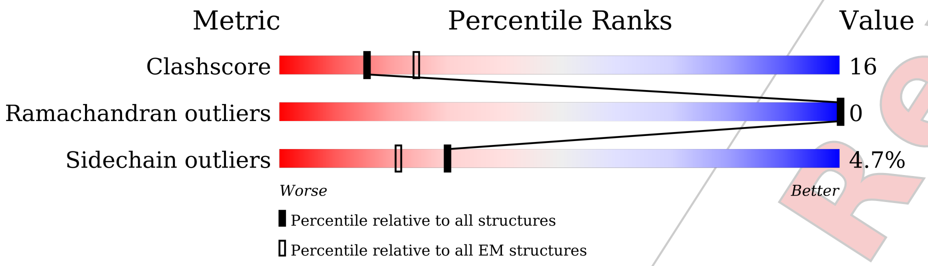


| Metric | Whole archive (#Entries) | EM structures (#Entries) |
|---|---|---|
| Clashscore | 158937 | 4297 |
| Ramachandran outliers | 154571 | 4023 |
| Sidechain outliers | 154315 | 3826 |

The table below summarises the geometric issues observed across the polymeric chains and their fit to the map. The red, orange, yellow and green segments of the bar indicate the fraction of residues that contain outliers for >=3, 2, 1 and 0 types of geometric quality criteria respectively. A grey segment represents the fraction of residues that are not modelled. The numeric value for each fraction is indicated below the corresponding segment, with a dot representing fractions <=5% The upper red bar (where present) indicates the fraction of residues that have poor fit to the EM map (all-atom inclusion < 40%). The numeric value is given above the bar.

| Mol | Chain | Length | Quality of chain |
|---|---|---|---|
| 1 | A | 726 | • / 25% 12% • 63% |
| 1 | B | 726 | • / 25% 11% • 63% |
| 1 | C | 726 | • / 25% 11% • 63% |
| 1 | D | 726 | • / 25% 12% • 63% |
| 1 | E | 726 | •• 98% |
| 1 | F | 726 | •• 98% |
| 1 | G | 726 | •• 98% |

*Continued on next page...*

For Manuscript Review

*Continued from previous page...*

| Mol | Chain | Length | Quality of chain |
|---|---|---|---|
| 1 | H | 726 | 98% |

For Manuscript Review

#### 2 Entry composition (i)

There are 4 unique types of molecules in this entry. The entry contains 8932 atoms, of which 0 are hydrogens and 0 are deuteriums.

In the tables below, the AltConf column contains the number of residues with at least one atom in alternate conformation and the Trace column contains the number of residues modelled with at most 2 atoms.

- Molecule 1 is a protein called Delta-1-pyrroline-5-carboxylate synthase B.

| Mol | Chain | Residues | Atoms | AltConf | Trace |
|---|---|---|---|---|---|
| 1 | C | 270 | Total C N O S<br>2067 1303 365 390 9 | 0 | 0 |
| 1 | F | 15 | Total C N O<br>124 79 22 23 | 0 | 0 |
| 1 | A | 270 | Total C N O S<br>2067 1303 365 390 9 | 0 | 0 |
| 1 | E | 15 | Total C N O<br>124 79 22 23 | 0 | 0 |
| 1 | B | 270 | Total C N O S<br>2067 1303 365 390 9 | 0 | 0 |
| 1 | G | 15 | Total C N O<br>124 79 22 23 | 0 | 0 |
| 1 | D | 270 | Total C N O S<br>2067 1303 365 390 9 | 0 | 0 |
| 1 | H | 15 | Total C N O<br>124 79 22 23 | 0 | 0 |

- Molecule 2 is ADENOSINE-5'-DIPHOSPHATE (three-letter code: ADP) (formula: $C_{10}H_{15}N_5O_{10}P_2$) (labeled as "Ligand of Interest" by depositor).

For Manuscript Review

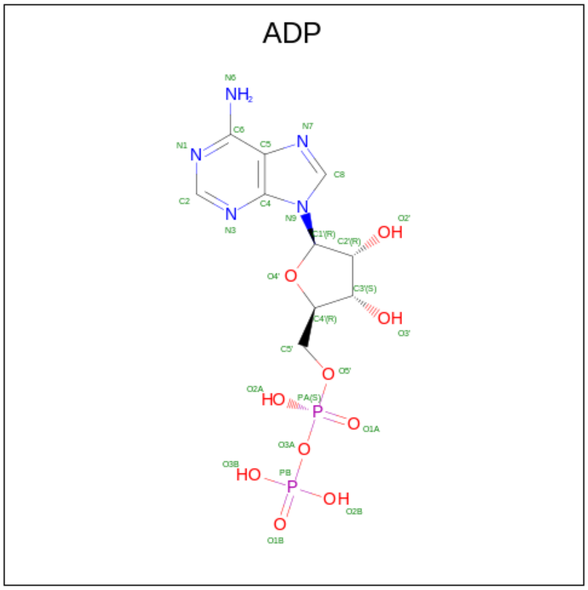


| Mol | Chain | Residues | Atoms | AltConf |
|---|---|---|---|---|
| 2 | C | 1 | Total C N O P<br>27 10 5 10 2 | 0 |
| 2 | A | 1 | Total C N O P<br>27 10 5 10 2 | 0 |
| 2 | B | 1 | Total C N O P<br>27 10 5 10 2 | 0 |
| 2 | D | 1 | Total C N O P<br>27 10 5 10 2 | 0 |

- Molecule 3 is GAMMA-GLUTAMYL PHOSPHATE (three-letter code: RGP) (formula: $C_5H_{10}NO_7P$) (labeled as "Ligand of Interest" by depositor).

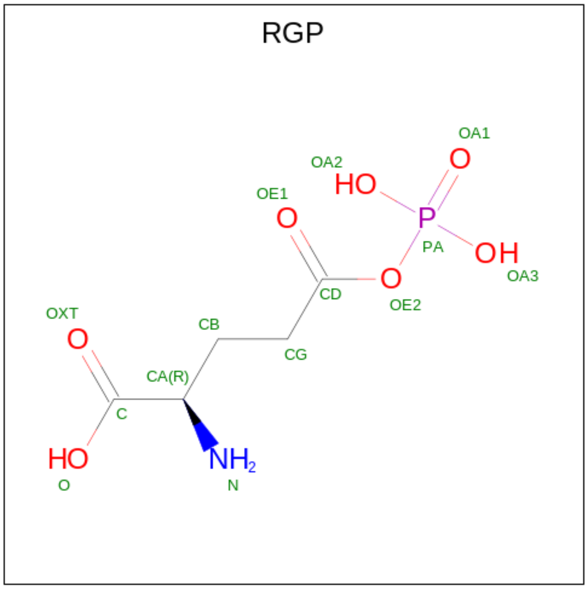


For Manuscript Review

| Mol | Chain | Residues | Atoms | | | | | AltConf |
|---|---|---|---|---|---|---|---|---|
| | | | Total | C | N | O | P | |
| 3 | C | 1 | 14 | 5 | 1 | 7 | 1 | 0 |
| 3 | A | 1 | 14 | 5 | 1 | 7 | 1 | 0 |
| 3 | B | 1 | 14 | 5 | 1 | 7 | 1 | 0 |
| 3 | D | 1 | 14 | 5 | 1 | 7 | 1 | 0 |

- Molecule 4 is MAGNESIUM ION (three-letter code: MG) (formula: Mg) (labeled as "Ligand of Interest" by depositor).

| Mol | Chain | Residues | Atoms | | AltConf |
|---|---|---|---|---|---|
| | | | Total | Mg | |
| 4 | C | 1 | 1 | 1 | 0 |
| 4 | A | 1 | 1 | 1 | 0 |
| 4 | B | 1 | 1 | 1 | 0 |
| 4 | D | 1 | 1 | 1 | 0 |

For Manuscript Review

### 3 Residue-property plots (i)

These plots are drawn for all protein, RNA, DNA and oligosaccharide chains in the entry. The first graphic for a chain summarises the proportions of the various outlier classes displayed in the second graphic. The second graphic shows the sequence view annotated by issues in geometry and atom inclusion in map density. Residues are color-coded according to the number of geometric quality criteria for which they contain at least one outlier: green = 0, yellow = 1, orange = 2 and red = 3 or more. A red diamond above a residue indicates a poor fit to the EM map for this residue (all-atom inclusion < 40%). Stretches of 2 or more consecutive residues without any outlier are shown as a green connector. Residues present in the sample, but not in the model, are shown in grey.

- Molecule 1: Delta-1-pyrroline-5-carboxylate synthase B

Chain C: 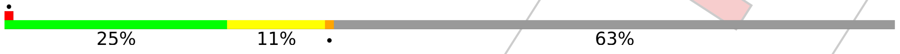


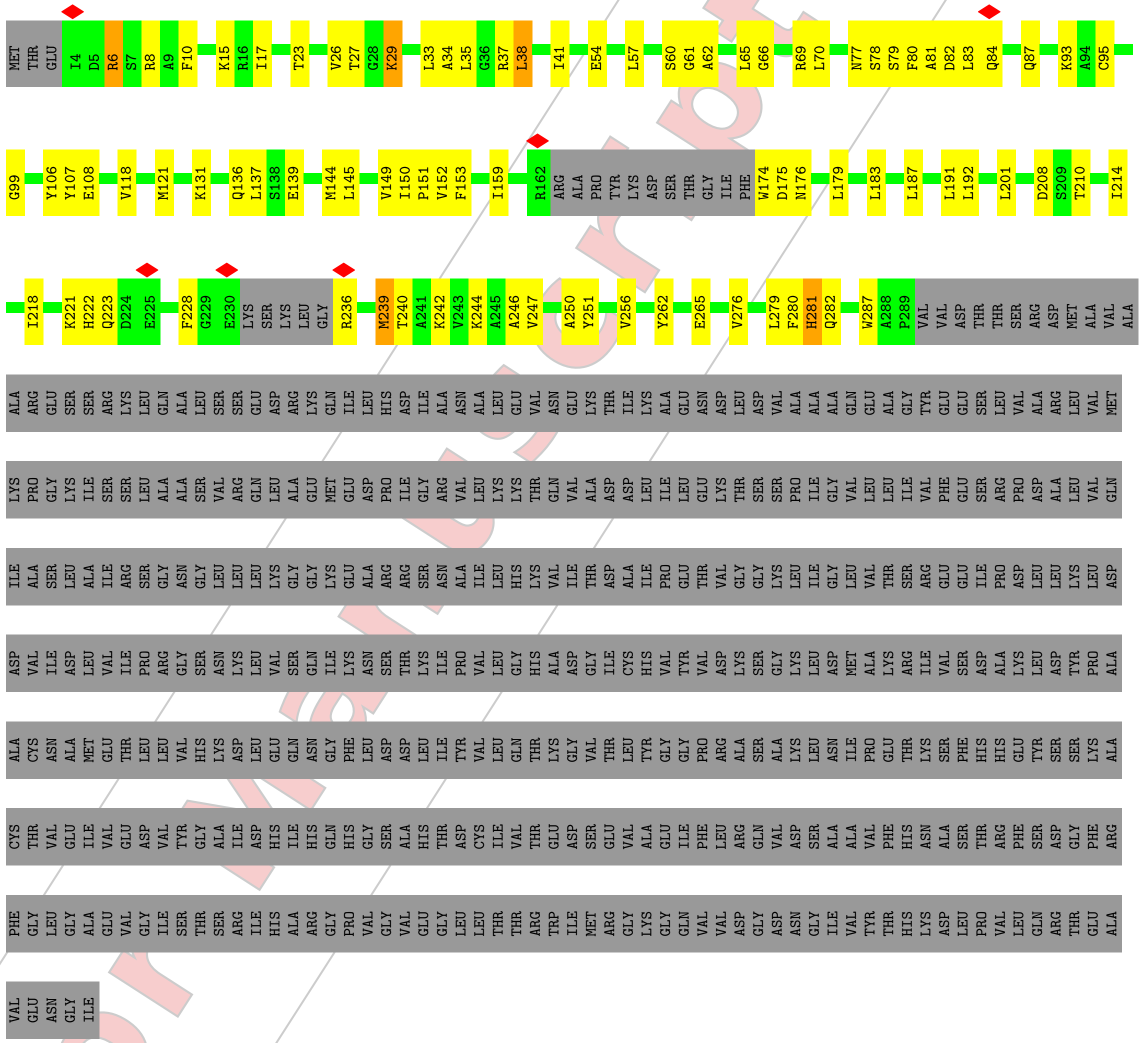

- Molecule 1: Delta-1-pyrroline-5-carboxylate synthase B

Chain F: 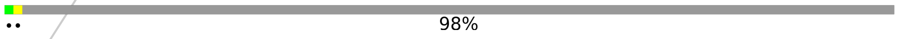


For Manuscript Review

MET THR GLU ILE ASP ARG SER ARG ALA PHE ALA LYS ASP VAL LYS ARG ILE VAL VAL LYS VAL GLY THR ALA VAL VAL THR GLY LYS GLY GLY ARG LEU ALA LEU GLY ARG LEU GLY ALA ILE CYS GLU GLN LEU ALA GLU LEU ASN SER ASP GLY PHE GLU VAL ILE LEU VAL SER SER

GLY ALA VAL GLY LEU GLY ARG GLN ARG LEU ARG Y72 R73 Q74 L75 V76 N77 S78 S79 D82 L83 Q84 K85 P86 GLN MET GLU LEU ASP GLY LYS ALA CYS ALA GLY VAL GLY GLN SER SER LEU MET ALA TYR TYR GLU THR MET PHE ASP GLN LEU ASP VAL THR VAL ALA ALA GLN MET

LEU VAL THR ASP SER SER PHE ARG ASP LYS ASP PHE ARG LYS GLN LEU SER GLU THR VAL LYS ALA MET LEU ARG MET ARG VAL ILE PRO VAL PHE ASN GLU ASN ASP ALA ILE SER THR ARG ARG ALA PRO TYR LYS ASP SER THR GLY ILE PHE TRP ASP ASN ASP SER LEU ALA ALA

LEU LEU SER LEU GLU LEU LYS ALA ASP LEU LEU ILE LEU LEU SER ASP VAL GLU GLY LEU TYR THR GLY PRO PRO SER ASP SER THR SER LYS LEU ILE HIS THR PHE ILE LYS GLU LYS HIS GLN ASP GLU ILE THR PHE GLY GLU LYS SER LYS LEU GLY ARG GLY GLY MET THR ALA

LYS VAL LYS ALA ALA VAL ASN ALA ALA TYR GLY GLY VAL PRO VAL ILE ILE THR SER GLY TYR ALA ALA GLU ASN ILE SER LYS VAL LEU ARG GLY LEU ARG VAL GLY THR LEU PHE HIS GLN ASP ALA HIS LEU TRP ALA PRO VAL VAL ASP THR THR SER ARG ASP MET ALA VAL ALA

ALA ARG GLU SER SER ARG LYS LEU GLN ALA LEU SER SER GLU ASP ARG LYS GLN ILE LEU HIS ASP ILE ALA ASN ALA LEU GLU VAL ASN GLU LYS THR ILE LYS ALA GLU ASN ASP LEU ASP VAL ALA ALA ALA GLN GLU ALA GLY TYR GLU GLU SER LEU VAL ALA ARG LEU VAL MET

LYS PRO GLY LYS ILE SER SER LEU ALA ALA SER VAL ARG GLN LEU ALA GLU MET GLU ASP PRO ILE GLY ARG VAL LEU LYS LYS THR GLN VAL ALA ASP ASP LEU ILE LEU GLU LYS THR SER SER PRO ILE GLY VAL LEU LEU ILE VAL PHE GLU SER ARG PRO ASP ALA LEU VAL GLN

ILE ALA SER LEU ALA ILE ARG SER GLY ASN GLY LEU LEU LEU LYS GLY GLY LYS GLU ALA ARG ARG SER ASN ALA ILE LEU HIS LYS VAL ILE THR ASP ALA ILE PRO GLU THR VAL GLY GLY LYS LEU ILE GLY LEU VAL THR SER ARG GLU GLU ILE PRO ASP LEU LEU LYS LEU ASP

ASP VAL ILE ASP LEU VAL ILE PRO ARG GLY SER ASN LYS LEU VAL SER GLN ILE LYS ASN SER THR LYS ILE PRO VAL LEU GLY HIS ALA ASP GLY ILE CYS HIS VAL TYR VAL ASP LYS SER GLY LYS LEU ASP MET ALA LYS ARG ILE VAL SER ASP ALA LYS LEU ASP TYR PRO ALA

ALA CYS ASN ALA MET GLU THR LEU LEU VAL HIS LYS ASP LEU GLU GLN ASN GLY PHE LEU ASP ASP LEU ILE TYR VAL LEU GLN THR LYS GLY VAL THR LEU TYR GLY GLY PRO ARG ALA ALA SER ALA LYS LEU ASN ILE PRO GLU THR LYS SER PHE HIS HIS GLU TYR SER SER LYS ALA

CYS THR VAL GLU ILE VAL GLU ASP VAL TYR GLY ALA ILE ASP HIS ILE HIS GLN HIS GLY SER ALA HIS THR ASP CYS ILE VAL THR GLU ASP SER GLU VAL ALA GLU ILE PHE LEU ARG GLN VAL ASP SER ALA ALA VAL PHE HIS ASN ALA SER THR ARG PHE SER ASP GLY PHE ARG

PHE GLY LEU GLY ALA GLU VAL GLY ILE SER THR SER ARG ILE HIS ALA ARG GLY PRO VAL GLY VAL GLU GLY LEU LEU THR THR ARG TRP ILE MET ARG GLY LYS GLY GLN VAL VAL ASP GLY ASP ASN GLY ILE VAL TYR THR HIS LYS ASP LEU PRO VAL LEU GLN ARG THR GLU ALA

VAL GLU ASN GLY ILE

- Molecule 1: Delta-1-pyrroline-5-carboxylate synthase B

Chain A: 25% 12% 63%

MET THR GLU I4 D5 R6 S7 R8 A9 F10 A11 K12 K15 R16 I17 T23 V26 T27 G28 K29 L33 A34 L35 G36 R37 L38 I41 E54 L57 S60 G61 A62 L65 G66 R67 Q68 R69 N77 S78 S79 F80 A81 D82 L83 Q84 Q87 K93 A94

C95 G99 Y106 Y107 E108 V118 M121 L122 K131 Q136 L137 S138 E139 M144 L145 V149 I150 P151 V152 F153 R162 ARG ALA PRO TYR LYS ASP SER THR GLY ILE PHE W174 D175 N176 L179 L183 L187 L191 L192 L201 I214 I218

K221 H222 Q223 D224 E225 F228 G229 E230 LYS SER LYS LEU GLY R236 M239 T240 A241 K242 V243 K244 A245 A246 V247 A250 Y251 V256 Y262 E265 V276 L279 F280 H281 Q282 W287 A288 P289 VAL VAL ASP THR THR SER ARG ASP MET ALA VAL ALA ALA ARG GLU

SER SER ARG LYS LEU GLN ALA LEU SER SER GLU ASP ARG LYS GLN ILE LEU HIS ASP ILE ALA ASN ALA LEU GLU VAL ASN GLU LYS THR ILE LYS ALA GLU ASN ASP LEU ASP VAL ALA ALA ALA GLN GLU ALA GLY TYR GLU GLU SER LEU VAL ALA ARG LEU VAL MET LYS PRO GLY

For Manuscript Review

LYS ILE SER SER LEU ALA ALA SER VAL ARG GLN LEU ALA GLU MET GLU ASP PRO ILE GLY ARG VAL LEU LYS LYS THR GLN VAL ALA ASP ASP LEU ILE LEU GLU LYS THR SER SER PRO ILE GLY VAL LEU LEU ILE VAL PHE GLU SER ARG PRO ASP ALA LEU VAL GLN ILE ALA SER

LEU ALA ILE ARG SER GLY ASN GLY LEU LEU LYS GLY GLY LYS GLU ALA ARG ARG SER ASN ALA ILE LEU HIS LYS VAL ILE THR ASP ALA ILE PRO GLU THR VAL GLY GLY LYS LEU ILE GLY LEU VAL THR SER ARG GLU GLU ILE PRO ASP LEU LEU LYS LEU ASP ASP VAL ILE

ASP LEU VAL ILE PRO ARG GLY SER ASN LYS LEU VAL SER GLN ILE LYS ASN SER THR LYS ILE PRO VAL LEU GLY HIS ALA ASP GLY ILE CYS HIS VAL TYR VAL ASP LYS SER GLY LYS LEU ASP MET ALA LYS ARG ILE VAL SER ASP ALA LYS LEU ASP TYR PRO ALA ALA CYS ASN

ALA MET GLU THR LEU LEU VAL HIS LYS ASP LEU GLU GLN ASN GLY PHE LEU ASP ASP LEU ILE TYR VAL LEU GLN THR LYS GLY VAL THR LEU TYR GLY GLY PRO ARG ALA SER ALA LYS LEU ASN ILE PRO GLU THR LYS SER PHE HIS HIS GLU TYR SER SER LYS ALA CYS THR VAL

GLU ILE VAL GLU ASP VAL TYR GLY ALA ILE ASP HIS ILE HIS GLN HIS GLY SER ALA HIS THR ASP CYS ILE VAL THR GLU ASP SER GLU VAL ALA GLU ILE PHE LEU ARG GLN VAL ASP SER ALA ALA VAL PHE HIS ASN ALA SER THR ARG PHE SER ASP GLY PHE ARG PHE GLY LEU

GLY ALA GLU VAL GLY ILE SER THR SER ARG ILE HIS ALA ARG GLY PRO VAL GLY VAL GLU GLY LEU LEU THR THR ARG TRP ILE MET ARG GLY LYS GLY GLN VAL VAL ASP GLY ASP ASN GLY ILE VAL TYR THR HIS LYS ASP LEU PRO VAL LEU GLN ARG THR GLU ALA VAL GLU ASN

GLY ILE

- Molecule 1: Delta-1-pyrroline-5-carboxylate synthase B

Chain E: 98%

MET THR GLU ILE ASP ARG SER ARG ALA PHE ALA LYS ASP VAL LYS ARG ILE VAL VAL LYS VAL GLY THR ALA VAL VAL THR GLY LYS GLY GLY ARG LEU ALA LEU GLY ARG LEU GLY ALA ILE CYS GLU GLN LEU ALA GLU LEU ASN SER ASP GLY PHE GLU VAL ILE LEU VAL SER SER

GLY ALA VAL GLY LEU GLY ARG GLN ARG LEU ARG Y72 R73 Q74 L75 V76 N77 S78 S79 D82 L83 Q84 K85 P86 GLN MET GLU LEU ASP GLY LYS ALA CYS ALA GLY VAL GLY GLN SER SER LEU MET ALA TYR TYR GLU THR MET PHE ASP GLN LEU ASP VAL THR VAL ALA GLN MET

LEU VAL THR ASP SER SER PHE ARG ASP LYS ASP PHE ARG LYS GLN LEU SER GLU THR VAL LYS ALA MET LEU ARG MET ARG VAL ILE PRO VAL PHE ASN GLU ASN ASP ALA ILE SER THR ARG ARG ALA PRO TYR LYS ASP SER THR GLY ILE PHE TRP ASP ASN ASP SER LEU ALA ALA

LEU LEU SER LEU GLU LEU LYS ALA ASP LEU ILE LEU LEU SER ASP VAL GLU GLY LEU TYR THR GLY PRO PRO SER ASP SER THR SER LYS LEU ILE HIS THR PHE ILE LYS GLU LYS HIS GLN ASP GLU ILE THR PHE GLY GLU LYS SER LYS LEU GLY ARG GLY GLY MET THR ALA

LYS VAL LYS ALA ALA VAL ASN ALA ALA TYR GLY GLY VAL PRO VAL ILE ILE THR SER GLY TYR ALA ALA GLU ASN ILE SER LYS VAL LEU ARG GLY LEU ARG VAL GLY THR LEU PHE HIS GLN ASP ALA HIS LEU TRP ALA PRO VAL VAL ASP THR THR SER ARG ASP MET ALA VAL ALA

ALA ARG GLU SER SER ARG LYS LEU GLN ALA LEU SER SER GLU ASP ARG LYS GLN ILE LEU HIS ASP ILE ALA ASN ALA LEU GLU VAL ASN GLU LYS THR ILE LYS ALA GLU ASN ASP LEU ASP VAL ALA ALA ALA GLN GLU ALA GLY TYR GLU GLU SER LEU VAL ALA ARG LEU VAL MET

LYS PRO GLY LYS ILE SER SER LEU ALA ALA SER VAL ARG GLN LEU ALA GLU MET GLU ASP PRO ILE GLY ARG VAL LEU LYS LYS THR GLN VAL ALA ASP ASP LEU ILE LEU GLU LYS THR SER SER PRO ILE GLY VAL LEU LEU ILE VAL PHE GLU SER ARG PRO ASP ALA LEU VAL GLN

ILE ALA SER LEU ALA ILE ARG SER GLY ASN GLY LEU LEU LYS GLY GLY LYS GLU ALA ARG ARG SER ASN ALA ILE LEU HIS LYS VAL ILE THR ASP ALA ILE PRO GLU THR VAL GLY GLY LYS LEU ILE GLY LEU VAL THR SER ARG GLU GLU ILE PRO ASP LEU LEU LYS LEU ASP

ASP VAL ILE ASP LEU VAL ILE PRO ARG GLY SER ASN LYS LEU VAL SER GLN ILE LYS ASN SER THR LYS ILE PRO VAL LEU GLY HIS ALA ASP GLY ILE CYS HIS VAL TYR VAL ASP LYS SER GLY LYS LEU ASP MET ALA LYS ARG ILE VAL SER ASP ALA LYS LEU ASP TYR PRO ALA

ALA CYS ASN ALA MET GLU THR LEU LEU VAL HIS LYS ASP LEU GLU GLN ASN GLY PHE LEU ASP ASP LEU ILE TYR VAL LEU GLN THR LYS GLY VAL THR LEU TYR GLY GLY PRO ARG ALA SER ALA LYS LEU ASN ILE PRO GLU THR LYS SER PHE HIS HIS GLU TYR SER SER LYS ALA

CYS THR VAL GLU ILE VAL GLU ASP VAL TYR GLY ALA ILE ASP HIS ILE HIS GLN HIS GLY SER ALA HIS THR ASP CYS ILE VAL THR GLU ASP SER GLU VAL ALA GLU ILE PHE LEU ARG GLN VAL ASP SER ALA ALA VAL PHE HIS ASN ALA SER THR ARG PHE SER ASP GLY PHE ARG

For Manuscript Review

PHE GLY LEU GLY ALA GLU VAL GLY ILE SER THR SER ARG ILE HIS ALA ARG GLY PRO VAL GLY VAL GLU GLY LEU LEU THR THR ARG TRP ILE MET ARG GLY LYS GLY GLN VAL VAL ASP GLY ASP ASN GLY ILE VAL TYR THR HIS LYS ASP LEU PRO VAL LEU GLN ARG THR GLU ALA

VAL GLU ASN GLY ILE

• Molecule 1: Delta-1-pyrroline-5-carboxylate synthase B

Chain B: 25% 11% 63%

MET THR GLU I4 D5 R6 S7 R8 A9 F10 K15 R16 I17 T23 V26 T27 G28 K29 L33 A34 L35 G36 R37 L38 I41 E54 L57 S60 G61 A62 L65 G66 R69 L70 N77 S78 S79 F80 A81 D82 L83 Q84 Q87 K93 A94 C95

G99 Y106 Y107 E108 V118 M121 K131 Q136 L137 S138 E139 M144 L145 V149 I150 P151 V152 F153 I159 R162 ARG ALA PRO TYR LYS ASP SER THR GLY ILE PHE W174 D175 N176 L179 L183 L187 L191 L192 L201 D208 S209 T210 I214

I218 K221 H222 Q223 D224 E225 F228 G229 E230 LYS SER LYS LEU GLY R236 M239 T240 A241 K242 V243 K244 A245 A246 V247 A250 Y251 V256 Y262 E265 V276 L279 F280 H281 Q282 W287 A288 P289 VAL VAL ASP THR THR SER ARG ASP MET ALA VAL ALA

ALA ARG GLU SER SER ARG LYS LEU GLN ALA LEU SER SER GLU ASP ARG LYS GLN ILE LEU HIS ASP ILE ALA ASN ALA LEU GLU VAL ASN GLU LYS THR ILE LYS ALA GLU ASN ASP LEU ASP VAL ALA ALA ALA GLN GLU ALA GLY TYR GLU GLU SER LEU VAL ALA ARG LEU VAL MET

LYS PRO GLY LYS ILE SER SER LEU ALA ALA SER VAL ARG GLN LEU ALA GLU MET GLU ASP PRO ILE GLY ARG VAL LEU LYS LYS THR GLN VAL ALA ASP ASP LEU ILE LEU GLU LYS THR SER SER PRO ILE GLY VAL LEU LEU ILE VAL PHE GLU SER ARG PRO ASP ALA LEU VAL GLN

ILE ALA SER LEU ALA ILE ARG SER GLY ASN GLY LEU LEU LEU LYS GLY GLY LYS GLU ALA ARG ARG SER ASN ALA ILE LEU HIS LYS VAL ILE THR ASP ALA ILE PRO GLU THR VAL GLY GLY LYS LEU ILE GLY LEU VAL THR SER ARG GLU GLU ILE PRO ASP LEU LEU LYS LEU ASP

ASP VAL ILE ASP LEU VAL ILE PRO ARG GLY SER ASN LYS LEU VAL SER GLN ILE LYS ASN SER THR LYS ILE PRO VAL LEU GLY HIS ALA ASP GLY ILE CYS HIS VAL TYR VAL ASP LYS SER GLY LYS LEU ASP MET ALA LYS ARG ILE VAL SER ASP ALA LYS LEU ASP TYR PRO ALA

ALA CYS ASN ALA MET GLU THR LEU LEU VAL HIS LYS ASP LEU GLU GLN ASN GLY PHE LEU ASP ASP LEU ILE TYR VAL LEU GLN THR LYS GLY VAL THR LEU TYR GLY GLY PRO ARG ALA SER ALA LYS LEU ASN ILE PRO GLU THR LYS SER PHE HIS HIS GLU TYR SER SER LYS ALA

CYS THR VAL GLU ILE VAL GLU ASP VAL TYR GLY ALA ILE ASP HIS ILE HIS GLN HIS GLY SER ALA HIS THR ASP CYS ILE VAL THR GLU ASP SER GLU VAL ALA GLU ILE PHE LEU ARG GLN VAL ASP SER ALA ALA VAL PHE HIS ASN ALA SER THR ARG PHE SER ASP GLY PHE ARG

PHE GLY LEU GLY ALA GLU VAL GLY ILE SER THR SER ARG ILE HIS ALA ARG GLY PRO VAL GLY VAL GLU GLY LEU LEU THR THR ARG TRP ILE MET ARG GLY LYS GLY GLN VAL VAL ASP GLY ASP ASN GLY ILE VAL TYR THR HIS LYS ASP LEU PRO VAL LEU GLN ARG THR GLU ALA

VAL GLU ASN GLY ILE

• Molecule 1: Delta-1-pyrroline-5-carboxylate synthase B

Chain G: 98%

MET THR GLU ILE ASP ARG SER ARG ALA PHE ALA LYS ASP VAL LYS ARG ILE VAL VAL LYS VAL GLY THR ALA VAL VAL THR GLY LYS GLY GLY ARG LEU ALA LEU GLY ARG LEU GLY ALA ILE CYS GLU GLN LEU ALA GLU LEU ASN SER ASP GLY PHE GLU VAL ILE LEU VAL SER SER

GLY ALA VAL GLY LEU GLY ARG GLN ARG LEU ARG Y72 R73 Q74 L75 V76 N77 S78 S79 D82 L83 Q84 K85 P86 GLN MET GLU LEU ASP GLY LYS ALA CYS ALA GLY VAL GLY GLN SER SER LEU MET ALA TYR TYR GLU THR MET PHE ASP GLN LEU ASP VAL THR VAL ALA GLN MET

For Manuscript Review

LEU VAL THR ASP SER SER PHE ARG ASP LYS ASP PHE ARG LYS GLN LEU SER GLU THR VAL LYS ALA MET LEU ARG MET ARG VAL ILE PRO VAL PHE ASN GLU ASN ASP ALA ILE SER THR ARG ARG ALA PRO TYR LYS ASP SER THR GLY ILE PHE TRP ASP ASN ASP SER LEU ALA ALA

LEU LEU SER LEU GLU LEU LYS ALA ASP LEU LEU ILE LEU LEU SER ASP VAL GLU GLY LEU TYR THR GLY PRO PRO SER ASP SER THR SER LYS LEU ILE HIS THR PHE ILE LYS GLU LYS HIS GLN ASP GLU ILE THR PHE GLY GLU LYS SER LYS LEU GLY ARG GLY GLY MET THR ALA

LYS VAL LYS ALA ALA VAL ASN ALA ALA TYR GLY GLY VAL PRO VAL ILE ILE THR SER GLY TYR ALA ALA GLU ASN ILE SER LYS VAL LEU ARG GLY LEU ARG VAL GLY THR LEU PHE HIS GLN ASP ALA HIS LEU TRP ALA PRO VAL VAL ASP THR THR SER ARG ASP MET ALA VAL ALA

ALA ARG GLU SER SER ARG LYS LEU GLN ALA LEU SER SER GLU ASP ARG LYS GLN ILE LEU HIS ASP ILE ALA ASN ALA LEU GLU VAL ASN GLU LYS THR ILE LYS ALA GLU ASN ASP LEU ASP VAL ALA ALA ALA GLN GLU ALA GLY TYR GLU GLU SER LEU VAL ALA ARG LEU VAL MET

LYS PRO GLY LYS ILE SER SER LEU ALA ALA SER VAL ARG GLN LEU ALA GLU MET GLU ASP PRO ILE GLY ARG VAL LEU LYS LYS THR GLN VAL ALA ASP ASP LEU ILE LEU GLU LYS THR SER SER PRO ILE GLY VAL LEU LEU ILE VAL PHE GLU SER ARG PRO ASP ALA LEU VAL GLN

ILE ALA SER LEU ALA ILE ARG SER GLY ASN GLY LEU LEU LEU LYS GLY GLY LYS GLU ALA ARG ARG SER ASN ALA ILE LEU HIS LYS VAL ILE THR ASP ALA ILE PRO GLU THR VAL GLY GLY LYS LEU ILE GLY LEU VAL THR SER ARG GLU GLU ILE PRO ASP LEU LEU LYS LEU ASP

ASP VAL ILE ASP LEU VAL ILE PRO ARG GLY SER ASN LYS LEU VAL SER GLN ILE LYS ASN SER THR LYS ILE PRO VAL LEU GLY HIS ALA ASP GLY ILE CYS HIS VAL TYR VAL ASP LYS SER GLY LYS LEU ASP MET ALA LYS ARG ILE VAL SER ASP ALA LYS LEU ASP TYR PRO ALA

ALA CYS ASN ALA MET GLU THR LEU LEU VAL HIS LYS ASP LEU GLU GLN ASN GLY PHE LEU ASP ASP LEU ILE TYR VAL LEU GLN THR LYS GLY VAL THR LEU TYR GLY GLY PRO ARG ALA SER ALA LYS LEU ASN ILE PRO GLU THR LYS SER PHE HIS HIS GLU TYR SER SER LYS ALA

CYS THR VAL GLU ILE VAL GLU ASP VAL TYR GLY ALA ILE ASP HIS ILE HIS GLN HIS GLY SER ALA HIS THR ASP CYS ILE VAL THR GLU ASP SER GLU VAL ALA GLU ILE PHE LEU ARG GLN VAL ASP SER ALA ALA VAL PHE HIS ASN ALA SER THR ARG PHE SER ASP GLY PHE ARG

PHE GLY LEU GLY ALA GLU VAL GLY ILE SER THR SER ARG ILE HIS ALA ARG GLY PRO VAL GLY VAL GLU GLY LEU LEU THR THR ARG TRP ILE MET ARG GLY LYS GLY GLN VAL VAL ASP GLY ASP ASN GLY ILE VAL TYR THR HIS LYS ASP LEU PRO VAL LEU GLN ARG THR GLU ALA

VAL GLU ASN GLY ILE

- Molecule 1: Delta-1-pyrroline-5-carboxylate synthase B

Chain D: 25% 12% 63%

MET THR GLU I4 D5 R6 S7 R8 A9 F10 A11 K12 K15 R16 I17 T23 V26 T27 G28 K29 L33 A34 L35 G36 R37 L38 I41 E54 L57 S60 G61 A62 L65 G66 R67 Q68 R69 N77 S78 S79 F80 A81 D82 L83 Q84 Q87 K93 A94

C95 G99 Y106 Y107 E108 V118 M121 L122 K131 Q136 L137 S138 E139 M144 L145 V149 I150 P151 V152 F153 R162 ARG ALA PRO TYR LYS ASP SER THR GLY ILE PHE W174 D175 N176 L179 L183 L187 L191 L192 L201 I214 I218

K221 H222 Q223 D224 E225 F228 G229 E230 LYS SER LYS LEU GLY R236 M239 T240 A241 K242 V243 K244 A245 A246 V247 A250 Y251 V256 Y262 E265 V276 L279 F280 H281 Q282 W287 A288 P289 VAL VAL ASP THR THR SER ARG ASP MET ALA VAL ALA ALA ARG GLU

SER SER ARG LYS LEU GLN ALA LEU SER SER GLU ASP ARG LYS GLN ILE LEU HIS ASP ILE ALA ASN ALA LEU GLU VAL ASN GLU LYS THR ILE LYS ALA GLU ASN ASP LEU ASP VAL ALA ALA ALA GLN GLU ALA GLY TYR GLU GLU SER LEU VAL ALA ARG LEU VAL MET LYS PRO GLY

LYS ILE SER SER LEU ALA ALA SER VAL ARG GLN LEU ALA GLU MET GLU ASP PRO ILE GLY ARG VAL LEU LYS LYS THR GLN VAL ALA ASP ASP LEU ILE LEU GLU LYS THR SER SER PRO ILE GLY VAL LEU LEU ILE VAL PHE GLU SER ARG PRO ASP ALA LEU VAL GLN ILE ALA SER

LEU ALA ILE ARG SER GLY ASN GLY LEU LEU LEU LYS GLY GLY LYS GLU ALA ARG ARG SER ASN ALA ILE LEU HIS LYS VAL ILE THR ASP ALA ILE PRO GLU THR VAL GLY GLY LYS LEU ILE GLY LEU VAL THR SER ARG GLU GLU ILE PRO ASP LEU LEU LYS LEU ASP ASP VAL ILE

For Manuscript Review

ASP LEU VAL ILE PRO ARG GLY SER ASN LYS LEU VAL SER GLN ILE LYS ASN SER THR LYS ILE PRO VAL LEU GLY HIS ALA ASP GLY ILE CYS HIS VAL TYR VAL ASP LYS SER GLY LYS LEU ASP MET ALA LYS ARG ILE VAL SER ASP ALA LYS LEU ASP TYR PRO ALA ALA CYS ASN

ALA MET GLU THR LEU LEU VAL HIS LYS ASP LEU GLU GLN ASN GLY PHE LEU ASP ASP LEU ILE TYR VAL LEU GLN THR LYS GLY VAL THR LEU TYR GLY GLY PRO ARG ALA SER ALA LYS LEU ASN ILE PRO GLU THR LYS SER PHE HIS HIS GLU TYR SER SER LYS ALA CYS THR VAL

GLU ILE VAL GLU ASP VAL TYR GLY ALA ILE ASP HIS ILE HIS GLN HIS GLY SER ALA HIS THR ASP CYS ILE VAL THR GLU ASP SER GLU VAL ALA GLU ILE PHE LEU ARG GLN VAL ASP SER ALA ALA VAL PHE HIS ASN ALA SER THR ARG PHE SER ASP GLY PHE ARG PHE GLY LEU

GLY ALA GLU VAL GLY ILE SER THR SER ARG ILE HIS ALA ARG GLY PRO VAL GLY VAL GLU GLY LEU LEU THR THR ARG TRP ILE MET ARG GLY LYS GLY GLN VAL VAL ASP GLY ASP ASN GLY ILE VAL TYR THR HIS LYS ASP LEU PRO VAL LEU GLN ARG THR GLU ALA VAL GLU ASN

GLY ILE

• Molecule 1: Delta-1-pyrroline-5-carboxylate synthase B

Chain H: 98%

MET THR GLU ILE ASP ARG SER ARG ALA PHE ALA LYS ASP VAL LYS ARG ILE VAL VAL LYS VAL GLY THR ALA VAL VAL THR GLY LYS GLY GLY ARG LEU ALA LEU GLY ARG LEU GLY ALA ILE CYS GLU GLN LEU ALA GLU LEU ASN SER ASP GLY PHE GLU VAL ILE LEU VAL SER SER

GLY ALA VAL GLY LEU GLY ARG GLN ARG LEU ARG Y72 R73 Q74 L75 V76 N77 S78 S79 D82 L83 Q84 K85 P86 GLN MET GLU LEU ASP GLY LYS ALA CYS ALA GLY VAL GLY GLN SER SER LEU MET ALA TYR TYR GLU THR MET PHE ASP GLN LEU ASP VAL THR VAL ALA GLN MET

LEU VAL THR ASP SER SER PHE ARG ASP LYS ASP PHE ARG LYS GLN LEU SER GLU THR VAL LYS ALA MET LEU ARG MET ARG VAL ILE PRO VAL PHE ASN GLU ASN ASP ALA ILE SER THR ARG ARG ALA PRO TYR LYS ASP SER THR GLY ILE PHE TRP ASP ASN ASP SER LEU ALA ALA

LEU LEU SER LEU GLU LEU LYS ALA ASP LEU LEU ILE LEU LEU SER ASP VAL GLU GLY LEU TYR THR GLY PRO PRO SER ASP SER THR SER LYS LEU ILE HIS THR PHE ILE LYS GLU LYS HIS GLN ASP GLU ILE THR PHE GLY GLU LYS SER LYS LEU GLY ARG GLY GLY MET THR ALA

LYS VAL LYS ALA ALA VAL ASN ALA ALA TYR GLY GLY VAL PRO VAL ILE ILE THR SER GLY TYR ALA ALA GLU ASN ILE SER LYS VAL LEU ARG GLY LEU ARG VAL GLY THR LEU PHE HIS GLN ASP ALA HIS LEU TRP ALA PRO VAL VAL ASP THR THR SER ARG ASP MET ALA VAL ALA

ALA ARG GLU SER SER ARG LYS LEU GLN ALA LEU SER SER GLU ASP ARG LYS GLN ILE LEU HIS ASP ILE ALA ASN ALA LEU GLU VAL ASN GLU LYS THR ILE LYS ALA GLU ASN ASP LEU ASP VAL ALA ALA ALA GLN GLU ALA GLY TYR GLU GLU SER LEU VAL ALA ARG LEU VAL MET

LYS PRO GLY LYS ILE SER SER LEU ALA ALA SER VAL ARG GLN LEU ALA GLU MET GLU ASP PRO ILE GLY ARG VAL LEU LYS LYS THR GLN VAL ALA ASP ASP LEU ILE LEU GLU LYS THR SER SER PRO ILE GLY VAL LEU LEU ILE VAL PHE GLU SER ARG PRO ASP ALA LEU VAL GLN

ILE ALA SER LEU ALA ILE ARG SER GLY ASN GLY LEU LEU LEU LYS GLY GLY LYS GLU ALA ARG ARG SER ASN ALA ILE LEU HIS LYS VAL ILE THR ASP ALA ILE PRO GLU THR VAL GLY GLY LYS LEU ILE GLY LEU VAL THR SER ARG GLU GLU ILE PRO ASP LEU LEU LYS LEU ASP

ASP VAL ILE ASP LEU VAL ILE PRO ARG GLY SER ASN LYS LEU VAL SER GLN ILE LYS ASN SER THR LYS ILE PRO VAL LEU GLY HIS ALA ASP GLY ILE CYS HIS VAL TYR VAL ASP LYS SER GLY LYS LEU ASP MET ALA LYS ARG ILE VAL SER ASP ALA LYS LEU ASP TYR PRO ALA

ALA CYS ASN ALA MET GLU THR LEU LEU VAL HIS LYS ASP LEU GLU GLN ASN GLY PHE LEU ASP ASP LEU ILE TYR VAL LEU GLN THR LYS GLY VAL THR LEU TYR GLY GLY PRO ARG ALA SER ALA LYS LEU ASN ILE PRO GLU THR LYS SER PHE HIS HIS GLU TYR SER SER LYS ALA

CYS THR VAL GLU ILE VAL GLU ASP VAL TYR GLY ALA ILE ASP HIS ILE HIS GLN HIS GLY SER ALA HIS THR ASP CYS ILE VAL THR GLU ASP SER GLU VAL ALA GLU ILE PHE LEU ARG GLN VAL ASP SER ALA ALA VAL PHE HIS ASN ALA SER THR ARG PHE SER ASP GLY PHE ARG

PHE GLY LEU GLY ALA GLU VAL GLY ILE SER THR SER ARG ILE HIS ALA ARG GLY PRO VAL GLY VAL GLU GLY LEU LEU THR THR ARG TRP ILE MET ARG GLY LYS GLY GLN VAL VAL ASP GLY ASP ASN GLY ILE VAL TYR THR HIS LYS ASP LEU PRO VAL LEU GLN ARG THR GLU ALA

VAL GLU ASN GLY ILE

For Manuscript Review

### 4 Experimental information ⓘ

| Property | Value | Source |
|---|---|---|
| EM reconstruction method | SINGLE PARTICLE | Depositor |
| Imposed symmetry | POINT, D2 | Depositor |
| Number of particles used | 223041 | Depositor |
| Resolution determination method | FSC 0.143 CUT-OFF | Depositor |
| CTF correction method | NONE | Depositor |
| Microscope | FEI TITAN KRIOS | Depositor |
| Voltage (kV) | 300 | Depositor |
| Electron dose $(e^{-}/\text{Å}^2)$ | 1.2 | Depositor |
| Minimum defocus (nm) | 1500 | Depositor |
| Maximum defocus (nm) | 2500 | Depositor |
| Magnification | Not provided | |
| Image detector | GATAN K3 (6k x 4k) | Depositor |
| Maximum map value | 0.022 | Depositor |
| Minimum map value | -0.010 | Depositor |
| Average map value | 0.000 | Depositor |
| Map value standard deviation | 0.001 | Depositor |
| Recommended contour level | 0.005 | Depositor |
| Map size (Å) | 254.4, 254.4, 254.4 | wwPDB |
| Map dimensions | 240, 240, 240 | wwPDB |
| Map angles (°) | 90.0, 90.0, 90.0 | wwPDB |
| Pixel spacing (Å) | 1.06, 1.06, 1.06 | Depositor |

For Manuscript Review

### 5 Model quality (i)

#### 5.1 Standard geometry (i)

Bond lengths and bond angles in the following residue types are not validated in this section: ADP, RGP, MG

The Z score for a bond length (or angle) is the number of standard deviations the observed value is removed from the expected value. A bond length (or angle) with $|Z| > 5$ is considered an outlier worth inspection. RMSZ is the root-mean-square of all Z scores of the bond lengths (or angles).

| Mol | Chain | Bond lengths | | Bond angles | |
|---|---|---|---|---|---|
| | | RMSZ | $\#|Z| > 5$ | RMSZ | $\#|Z| > 5$ |
| 1 | A | 0.39 | 0/2094 | 0.59 | 0/2821 |
| 1 | B | 0.39 | 0/2094 | 0.59 | 0/2821 |
| 1 | C | 0.39 | 0/2094 | 0.59 | 0/2821 |
| 1 | D | 0.39 | 0/2094 | 0.59 | 0/2821 |
| 1 | E | 0.40 | 0/126 | 0.42 | 0/169 |
| 1 | F | 0.40 | 0/126 | 0.42 | 0/169 |
| 1 | G | 0.40 | 0/126 | 0.42 | 0/169 |
| 1 | H | 0.40 | 0/126 | 0.42 | 0/169 |
| All | All | 0.39 | 0/8880 | 0.59 | 0/11960 |

There are no bond length outliers.

There are no bond angle outliers.

There are no chirality outliers.

There are no planarity outliers.

#### 5.2 Too-close contacts (i)

In the following table, the Non-H and H(model) columns list the number of non-hydrogen atoms and hydrogen atoms in the chain respectively. The H(added) column lists the number of hydrogen atoms added and optimized by MolProbity. The Clashes column lists the number of clashes within the asymmetric unit, whereas Symm-Clashes lists symmetry-related clashes.

| Mol | Chain | Non-H | H(model) | H(added) | Clashes | Symm-Clashes |
|---|---|---|---|---|---|---|
| 1 | A | 2067 | 0 | 2110 | 69 | 0 |
| 1 | B | 2067 | 0 | 2110 | 67 | 0 |
| 1 | C | 2067 | 0 | 2110 | 67 | 0 |
| 1 | D | 2067 | 0 | 2110 | 68 | 0 |
| 1 | E | 124 | 0 | 122 | 12 | 0 |

*Continued on next page...*

For Manuscript Review

*Continued from previous page...*

| Mol | Chain | Non-H | H(model) | H(added) | Clashes | Symm-Clashes |
|---|---|---|---|---|---|---|
| 1 | F | 124 | 0 | 122 | 12 | 0 |
| 1 | G | 124 | 0 | 122 | 12 | 0 |
| 1 | H | 124 | 0 | 122 | 12 | 0 |
| 2 | A | 27 | 0 | 12 | 2 | 0 |
| 2 | B | 27 | 0 | 12 | 2 | 0 |
| 2 | C | 27 | 0 | 12 | 2 | 0 |
| 2 | D | 27 | 0 | 12 | 1 | 0 |
| 3 | A | 14 | 0 | 7 | 2 | 0 |
| 3 | B | 14 | 0 | 7 | 2 | 0 |
| 3 | C | 14 | 0 | 7 | 2 | 0 |
| 3 | D | 14 | 0 | 7 | 2 | 0 |
| 4 | A | 1 | 0 | 0 | 0 | 0 |
| 4 | B | 1 | 0 | 0 | 0 | 0 |
| 4 | C | 1 | 0 | 0 | 0 | 0 |
| 4 | D | 1 | 0 | 0 | 0 | 0 |
| All | All | 8932 | 0 | 9004 | 293 | 0 |

The all-atom clashscore is defined as the number of clashes found per 1000 atoms (including hydrogen atoms). The all-atom clashscore for this structure is 16.

All (293) close contacts within the same asymmetric unit are listed below, sorted by their clash magnitude.

| Atom-1 | Atom-2 | Interatomic distance (Å) | Clash overlap (Å) |
|---|---|---|---|
| 1:A:279:LEU:HD21 | 1:A:281:HIS:HB2 | 1.71 | 0.72 |
| 1:D:279:LEU:HD21 | 1:D:281:HIS:HB2 | 1.71 | 0.72 |
| 1:B:279:LEU:HD21 | 1:B:281:HIS:HB2 | 1.71 | 0.71 |
| 1:C:279:LEU:HD21 | 1:C:281:HIS:HB2 | 1.71 | 0.70 |
| 1:C:26:VAL:HG11 | 1:C:60:SER:O | 1.96 | 0.66 |
| 1:B:26:VAL:HG11 | 1:B:60:SER:O | 1.96 | 0.66 |
| 1:C:80:PHE:HA | 1:C:83:LEU:HD23 | 1.77 | 0.65 |
| 1:B:80:PHE:HA | 1:B:83:LEU:HD23 | 1.77 | 0.65 |
| 1:D:80:PHE:HA | 1:D:83:LEU:HD23 | 1.77 | 0.65 |
| 1:A:80:PHE:HA | 1:A:83:LEU:HD23 | 1.77 | 0.64 |
| 1:A:26:VAL:HG11 | 1:A:60:SER:O | 1.96 | 0.64 |
| 1:D:26:VAL:HG11 | 1:D:60:SER:O | 1.96 | 0.64 |
| 1:D:246:ALA:HB1 | 1:D:256:VAL:HG11 | 1.80 | 0.64 |
| 1:A:246:ALA:HB1 | 1:A:256:VAL:HG11 | 1.80 | 0.64 |
| 1:B:246:ALA:HB1 | 1:B:256:VAL:HG11 | 1.80 | 0.63 |
| 1:C:246:ALA:HB1 | 1:C:256:VAL:HG11 | 1.80 | 0.63 |
| 1:A:214:ILE:HD12 | 1:A:222:HIS:CE1 | 2.34 | 0.63 |
| 1:D:214:ILE:HD12 | 1:D:222:HIS:CE1 | 2.34 | 0.63 |

*Continued on next page...*

For Manuscript Review

*Continued from previous page...*

| Atom-1 | Atom-2 | Interatomic distance (Å) | Clash overlap (Å) |
|---|---|---|---|
| 1:C:214:ILE:HD12 | 1:C:222:HIS:CE1 | 2.34 | 0.63 |
| 1:C:80:PHE:CE1 | 1:E:75:LEU:HB3 | 2.34 | 0.62 |
| 1:C:265:GLU:OE1 | 1:C:265:GLU:N | 2.24 | 0.62 |
| 1:B:214:ILE:HD12 | 1:B:222:HIS:CE1 | 2.34 | 0.62 |
| 1:A:15:LYS:NZ | 1:A:54:GLU:OE2 | 2.26 | 0.62 |
| 1:B:80:PHE:CE1 | 1:H:75:LEU:HB3 | 2.34 | 0.62 |
| 1:D:15:LYS:NZ | 1:D:54:GLU:OE2 | 2.26 | 0.62 |
| 1:A:250:ALA:O | 1:A:282:GLN:NE2 | 2.33 | 0.62 |
| 1:B:250:ALA:O | 1:B:282:GLN:NE2 | 2.33 | 0.62 |
| 1:B:265:GLU:OE1 | 1:B:265:GLU:N | 2.24 | 0.62 |
| 1:D:250:ALA:O | 1:D:282:GLN:NE2 | 2.33 | 0.62 |
| 1:C:250:ALA:O | 1:C:282:GLN:NE2 | 2.33 | 0.62 |
| 1:A:62:ALA:O | 1:A:99:GLY:HA3 | 2.00 | 0.61 |
| 1:A:87:GLN:OE1 | 1:A:87:GLN:N | 2.26 | 0.61 |
| 1:D:62:ALA:O | 1:D:99:GLY:HA3 | 2.00 | 0.61 |
| 1:D:87:GLN:OE1 | 1:D:87:GLN:N | 2.26 | 0.61 |
| 1:F:75:LEU:HB3 | 1:A:80:PHE:CE1 | 2.34 | 0.61 |
| 1:C:62:ALA:O | 1:C:99:GLY:HA3 | 2.00 | 0.61 |
| 1:B:62:ALA:O | 1:B:99:GLY:HA3 | 2.00 | 0.61 |
| 1:C:15:LYS:NZ | 1:C:54:GLU:OE2 | 2.26 | 0.61 |
| 1:G:75:LEU:HB3 | 1:D:80:PHE:CE1 | 2.34 | 0.61 |
| 1:B:15:LYS:NZ | 1:B:54:GLU:OE2 | 2.26 | 0.60 |
| 1:D:265:GLU:OE1 | 1:D:265:GLU:N | 2.24 | 0.60 |
| 1:A:265:GLU:OE1 | 1:A:265:GLU:N | 2.24 | 0.60 |
| 1:E:78:SER:OG | 1:E:82:ASP:HB2 | 2.03 | 0.58 |
| 1:A:244:LYS:O | 1:A:247:VAL:HG12 | 2.04 | 0.58 |
| 1:H:78:SER:OG | 1:H:82:ASP:HB2 | 2.03 | 0.58 |
| 1:D:244:LYS:O | 1:D:247:VAL:HG12 | 2.04 | 0.58 |
| 1:F:78:SER:OG | 1:F:82:ASP:HB2 | 2.03 | 0.57 |
| 1:G:78:SER:OG | 1:G:82:ASP:HB2 | 2.03 | 0.57 |
| 1:E:73:ARG:O | 1:E:76:VAL:HG12 | 2.04 | 0.57 |
| 1:H:73:ARG:O | 1:H:76:VAL:HG12 | 2.04 | 0.57 |
| 1:G:73:ARG:O | 1:G:76:VAL:HG12 | 2.04 | 0.57 |
| 1:C:236:ARG:HH21 | 3:C:802:RGP:HB2 | 1.69 | 0.57 |
| 1:F:73:ARG:O | 1:F:76:VAL:HG12 | 2.04 | 0.57 |
| 1:B:236:ARG:HH21 | 3:B:802:RGP:HB2 | 1.69 | 0.57 |
| 1:B:244:LYS:O | 1:B:247:VAL:HG12 | 2.04 | 0.57 |
| 1:C:87:GLN:OE1 | 1:C:87:GLN:N | 2.26 | 0.57 |
| 1:C:244:LYS:O | 1:C:247:VAL:HG12 | 2.04 | 0.57 |
| 1:A:131:LYS:HE3 | 1:A:131:LYS:HA | 1.87 | 0.56 |
| 1:A:236:ARG:HH21 | 3:A:802:RGP:HB2 | 1.69 | 0.56 |

*Continued on next page...*

For Manuscript Review

Continued from previous page...

| Atom-1 | Atom-2 | Interatomic distance (Å) | Clash overlap (Å) |
|---|---|---|---|
| 1:B:87:GLN:OE1 | 1:B:87:GLN:N | 2.26 | 0.56 |
| 1:D:236:ARG:HH21 | 3:D:802:RGP:HB2 | 1.69 | 0.56 |
| 1:D:131:LYS:HA | 1:D:131:LYS:HE3 | 1.87 | 0.56 |
| 1:C:131:LYS:HE3 | 1:C:131:LYS:HA | 1.87 | 0.56 |
| 2:D:801:ADP:H2' | 2:D:801:ADP:N3 | 2.21 | 0.56 |
| 2:A:801:ADP:N3 | 2:A:801:ADP:H2' | 2.21 | 0.55 |
| 1:B:131:LYS:HE3 | 1:B:131:LYS:HA | 1.87 | 0.55 |
| 1:D:240:THR:O | 1:D:244:LYS:HG2 | 2.07 | 0.55 |
| 2:C:801:ADP:H2' | 2:C:801:ADP:N3 | 2.21 | 0.55 |
| 1:A:240:THR:O | 1:A:244:LYS:HG2 | 2.07 | 0.55 |
| 1:C:34:ALA:O | 1:C:38:LEU:HB2 | 2.07 | 0.55 |
| 2:B:801:ADP:H2' | 2:B:801:ADP:N3 | 2.21 | 0.55 |
| 1:E:85:LYS:HD3 | 1:E:85:LYS:C | 2.28 | 0.54 |
| 1:B:34:ALA:O | 1:B:38:LEU:HB2 | 2.07 | 0.54 |
| 1:D:34:ALA:O | 1:D:38:LEU:HB2 | 2.07 | 0.54 |
| 1:F:85:LYS:HD3 | 1:F:85:LYS:C | 2.28 | 0.54 |
| 1:A:34:ALA:O | 1:A:38:LEU:HB2 | 2.07 | 0.54 |
| 1:B:240:THR:O | 1:B:244:LYS:HG2 | 2.07 | 0.54 |
| 1:G:85:LYS:HD3 | 1:G:85:LYS:C | 2.28 | 0.54 |
| 1:H:85:LYS:C | 1:H:85:LYS:HD3 | 2.28 | 0.54 |
| 1:C:240:THR:O | 1:C:244:LYS:HG2 | 2.07 | 0.54 |
| 1:A:26:VAL:HG13 | 1:A:27:THR:HG23 | 1.90 | 0.54 |
| 1:D:26:VAL:HG13 | 1:D:27:THR:HG23 | 1.90 | 0.54 |
| 1:B:26:VAL:HG13 | 1:B:27:THR:HG23 | 1.90 | 0.54 |
| 1:C:26:VAL:HG13 | 1:C:27:THR:HG23 | 1.90 | 0.53 |
| 1:A:54:GLU:HB3 | 1:A:145:LEU:HD22 | 1.91 | 0.53 |
| 1:D:54:GLU:HB3 | 1:D:145:LEU:HD22 | 1.91 | 0.53 |
| 1:C:239:MET:HE3 | 1:C:242:LYS:HB3 | 1.91 | 0.53 |
| 1:A:183:LEU:HD12 | 1:A:187:LEU:HD11 | 1.90 | 0.53 |
| 1:D:183:LEU:HD12 | 1:D:187:LEU:HD11 | 1.90 | 0.53 |
| 1:B:239:MET:HE3 | 1:B:242:LYS:HB3 | 1.91 | 0.53 |
| 1:F:84:GLN:HG2 | 1:F:85:LYS:N | 2.24 | 0.53 |
| 1:G:84:GLN:HG2 | 1:G:85:LYS:N | 2.24 | 0.53 |
| 1:A:33:LEU:HD11 | 1:A:65:LEU:HD11 | 1.91 | 0.52 |
| 1:A:246:ALA:CB | 1:A:256:VAL:HG11 | 2.40 | 0.52 |
| 1:D:246:ALA:CB | 1:D:256:VAL:HG11 | 2.40 | 0.52 |
| 1:B:60:SER:OG | 3:B:802:RGP:HG2 | 2.09 | 0.52 |
| 1:B:183:LEU:HD12 | 1:B:187:LEU:HD11 | 1.90 | 0.52 |
| 1:D:33:LEU:HD11 | 1:D:65:LEU:HD11 | 1.91 | 0.52 |
| 1:C:60:SER:OG | 3:C:802:RGP:HG2 | 2.09 | 0.52 |
| 1:C:183:LEU:HD12 | 1:C:187:LEU:HD11 | 1.90 | 0.52 |

Continued on next page...

For Manuscript Review

Continued from previous page...

| Atom-1 | Atom-2 | Interatomic distance (Å) | Clash overlap (Å) |
|---|---|---|---|
| 1:C:54:GLU:HB3 | 1:C:145:LEU:HD22 | 1.91 | 0.52 |
| 1:E:84:GLN:HG2 | 1:E:85:LYS:N | 2.24 | 0.52 |
| 1:B:54:GLU:HB3 | 1:B:145:LEU:HD22 | 1.91 | 0.52 |
| 1:A:60:SER:OG | 3:A:802:RGP:HG2 | 2.09 | 0.52 |
| 1:D:60:SER:OG | 3:D:802:RGP:HG2 | 2.09 | 0.52 |
| 1:C:33:LEU:HD11 | 1:C:65:LEU:HD11 | 1.91 | 0.51 |
| 1:B:33:LEU:HD11 | 1:B:65:LEU:HD11 | 1.91 | 0.51 |
| 1:C:8:ARG:NH2 | 1:C:276:VAL:O | 2.38 | 0.51 |
| 1:C:77:ASN:O | 1:F:79:SER:HB2 | 2.10 | 0.51 |
| 1:C:256:VAL:HB | 1:C:280:PHE:HB2 | 1.92 | 0.51 |
| 1:B:77:ASN:O | 1:G:79:SER:HB2 | 2.10 | 0.51 |
| 1:B:256:VAL:HB | 1:B:280:PHE:HB2 | 1.92 | 0.51 |
| 1:A:239:MET:HE3 | 1:A:242:LYS:HB3 | 1.91 | 0.51 |
| 1:A:256:VAL:HB | 1:A:280:PHE:HB2 | 1.92 | 0.51 |
| 1:B:8:ARG:NH2 | 1:B:276:VAL:O | 2.38 | 0.51 |
| 1:D:77:ASN:O | 1:H:79:SER:HB2 | 2.10 | 0.51 |
| 1:D:239:MET:HE3 | 1:D:242:LYS:HB3 | 1.91 | 0.51 |
| 1:A:77:ASN:O | 1:E:79:SER:HB2 | 2.10 | 0.51 |
| 1:D:256:VAL:HB | 1:D:280:PHE:HB2 | 1.92 | 0.51 |
| 1:H:84:GLN:HG2 | 1:H:85:LYS:N | 2.24 | 0.51 |
| 1:C:246:ALA:CB | 1:C:256:VAL:HG11 | 2.40 | 0.51 |
| 1:B:246:ALA:CB | 1:B:256:VAL:HG11 | 2.40 | 0.51 |
| 1:C:60:SER:HB2 | 1:C:176:ASN:HD22 | 1.76 | 0.51 |
| 1:B:60:SER:HB2 | 1:B:176:ASN:HD22 | 1.76 | 0.51 |
| 1:A:87:GLN:H | 1:A:87:GLN:CD | 2.10 | 0.51 |
| 1:C:65:LEU:HD21 | 1:C:69:ARG:CZ | 2.41 | 0.50 |
| 1:A:65:LEU:HD21 | 1:A:69:ARG:CZ | 2.41 | 0.50 |
| 1:A:218:ILE:HG22 | 1:A:221:LYS:H | 1.76 | 0.50 |
| 1:B:87:GLN:H | 1:B:87:GLN:CD | 2.10 | 0.50 |
| 1:C:218:ILE:HG22 | 1:C:221:LYS:H | 1.76 | 0.50 |
| 1:D:218:ILE:HG22 | 1:D:221:LYS:H | 1.76 | 0.50 |
| 1:C:87:GLN:H | 1:C:87:GLN:CD | 2.10 | 0.50 |
| 1:B:65:LEU:HD21 | 1:B:69:ARG:CZ | 2.41 | 0.50 |
| 1:D:65:LEU:HD21 | 1:D:69:ARG:CZ | 2.41 | 0.50 |
| 1:D:87:GLN:H | 1:D:87:GLN:CD | 2.10 | 0.50 |
| 1:A:201:LEU:HD23 | 1:A:214:ILE:HG21 | 1.93 | 0.50 |
| 1:B:218:ILE:HG22 | 1:B:221:LYS:H | 1.76 | 0.50 |
| 1:D:201:LEU:HD23 | 1:D:214:ILE:HG21 | 1.93 | 0.50 |
| 1:A:60:SER:HB2 | 1:A:176:ASN:HD22 | 1.76 | 0.50 |
| 1:D:60:SER:HB2 | 1:D:176:ASN:HD22 | 1.76 | 0.49 |
| 1:C:79:SER:O | 1:C:82:ASP:N | 2.36 | 0.49 |

Continued on next page...

For Manuscript Review

*Continued from previous page...*

| Atom-1 | Atom-2 | Interatomic distance (Å) | Clash overlap (Å) |
|---|---|---|---|
| 1:B:79:SER:O | 1:B:82:ASP:N | 2.36 | 0.49 |
| 1:C:93:LYS:HD2 | 1:D:136:GLN:HE21 | 1.77 | 0.49 |
| 1:A:136:GLN:HE21 | 1:B:93:LYS:HD2 | 1.77 | 0.49 |
| 1:C:201:LEU:HD23 | 1:C:214:ILE:HG21 | 1.93 | 0.49 |
| 1:A:93:LYS:HD2 | 1:B:136:GLN:HE21 | 1.77 | 0.49 |
| 1:C:136:GLN:HE21 | 1:D:93:LYS:HD2 | 1.77 | 0.48 |
| 1:A:79:SER:O | 1:A:82:ASP:N | 2.36 | 0.48 |
| 1:B:201:LEU:HD23 | 1:B:214:ILE:HG21 | 1.93 | 0.48 |
| 1:A:80:PHE:HD2 | 1:E:76:VAL:HG23 | 1.78 | 0.48 |
| 1:D:80:PHE:HD2 | 1:H:76:VAL:HG23 | 1.78 | 0.48 |
| 1:B:80:PHE:HD2 | 1:G:76:VAL:HG23 | 1.78 | 0.48 |
| 1:C:80:PHE:HD2 | 1:F:76:VAL:HG23 | 1.78 | 0.48 |
| 1:D:79:SER:O | 1:D:82:ASP:N | 2.36 | 0.48 |
| 1:A:265:GLU:H | 1:A:265:GLU:CD | 2.14 | 0.48 |
| 1:D:57:LEU:O | 1:D:152:VAL:HA | 2.14 | 0.48 |
| 1:C:57:LEU:O | 1:C:152:VAL:HA | 2.14 | 0.48 |
| 1:A:57:LEU:O | 1:A:152:VAL:HA | 2.14 | 0.48 |
| 1:B:208:ASP:OD2 | 1:B:210:THR:OG1 | 2.25 | 0.48 |
| 1:B:57:LEU:O | 1:B:152:VAL:HA | 2.14 | 0.48 |
| 1:D:66:GLY:HA3 | 1:D:95:CYS:O | 2.14 | 0.48 |
| 1:A:66:GLY:HA3 | 1:A:95:CYS:O | 2.14 | 0.48 |
| 1:D:265:GLU:H | 1:D:265:GLU:CD | 2.14 | 0.48 |
| 1:C:80:PHE:CD2 | 1:F:76:VAL:HG23 | 2.50 | 0.47 |
| 1:B:80:PHE:CD2 | 1:G:76:VAL:HG23 | 2.50 | 0.47 |
| 1:C:33:LEU:HD13 | 1:C:106:TYR:CD2 | 2.50 | 0.47 |
| 1:C:137:LEU:HD21 | 1:C:183:LEU:HD11 | 1.97 | 0.47 |
| 1:A:33:LEU:HD13 | 1:A:106:TYR:CD2 | 2.50 | 0.47 |
| 1:B:137:LEU:HD21 | 1:B:183:LEU:HD11 | 1.97 | 0.47 |
| 1:D:23:THR:HG22 | 1:D:61:GLY:HA2 | 1.97 | 0.47 |
| 1:C:23:THR:HG22 | 1:C:61:GLY:HA2 | 1.97 | 0.47 |
| 1:C:66:GLY:HA3 | 1:C:95:CYS:O | 2.14 | 0.47 |
| 1:A:23:THR:HG22 | 1:A:61:GLY:HA2 | 1.97 | 0.47 |
| 1:B:23:THR:HG22 | 1:B:61:GLY:HA2 | 1.97 | 0.47 |
| 1:B:33:LEU:HD13 | 1:B:106:TYR:CD2 | 2.50 | 0.47 |
| 1:D:33:LEU:HD13 | 1:D:106:TYR:CD2 | 2.50 | 0.47 |
| 1:B:66:GLY:HA3 | 1:B:95:CYS:O | 2.14 | 0.47 |
| 1:A:80:PHE:CD2 | 1:E:76:VAL:HG23 | 2.50 | 0.47 |
| 1:D:80:PHE:CD2 | 1:H:76:VAL:HG23 | 2.50 | 0.46 |
| 1:C:265:GLU:H | 1:C:265:GLU:CD | 2.14 | 0.46 |
| 1:A:137:LEU:HD21 | 1:A:183:LEU:HD11 | 1.97 | 0.46 |
| 1:D:137:LEU:HD21 | 1:D:183:LEU:HD11 | 1.97 | 0.46 |

*Continued on next page...*

For Manuscript Review

*Continued from previous page...*

| Atom-1 | Atom-2 | Interatomic distance (Å) | Clash overlap (Å) |
|---|---|---|---|
| 1:C:287:TRP:N | 1:C:287:TRP:CD1 | 2.82 | 0.46 |
| 1:B:265:GLU:H | 1:B:265:GLU:CD | 2.14 | 0.46 |
| 1:C:29:LYS:HD2 | 1:C:29:LYS:C | 2.36 | 0.46 |
| 1:B:29:LYS:HD2 | 1:B:29:LYS:C | 2.36 | 0.46 |
| 1:F:74:GLN:HG2 | 1:F:75:LEU:N | 2.31 | 0.46 |
| 1:B:287:TRP:N | 1:B:287:TRP:CD1 | 2.82 | 0.46 |
| 1:C:208:ASP:OD2 | 1:C:210:THR:OG1 | 2.25 | 0.46 |
| 1:G:74:GLN:HG2 | 1:G:75:LEU:N | 2.31 | 0.46 |
| 1:A:29:LYS:C | 1:A:29:LYS:HD2 | 2.36 | 0.46 |
| 1:B:144:MET:CE | 1:B:151:PRO:HG3 | 2.46 | 0.46 |
| 1:C:144:MET:CE | 1:C:151:PRO:HG3 | 2.46 | 0.45 |
| 1:E:74:GLN:HG2 | 1:E:75:LEU:N | 2.31 | 0.45 |
| 1:D:29:LYS:C | 1:D:29:LYS:HD2 | 2.36 | 0.45 |
| 1:C:35:LEU:HD13 | 1:C:106:TYR:OH | 2.16 | 0.45 |
| 1:B:35:LEU:HD13 | 1:B:106:TYR:OH | 2.16 | 0.45 |
| 1:H:74:GLN:HG2 | 1:H:75:LEU:N | 2.31 | 0.45 |
| 1:D:35:LEU:HD13 | 1:D:106:TYR:OH | 2.16 | 0.45 |
| 1:D:287:TRP:N | 1:D:287:TRP:CD1 | 2.82 | 0.45 |
| 1:A:35:LEU:HD13 | 1:A:106:TYR:OH | 2.16 | 0.45 |
| 1:A:287:TRP:CD1 | 1:A:287:TRP:N | 2.82 | 0.45 |
| 1:D:37:ARG:O | 1:D:41:ILE:HG12 | 2.17 | 0.45 |
| 1:F:75:LEU:HB3 | 1:A:80:PHE:HE1 | 1.81 | 0.45 |
| 1:A:37:ARG:O | 1:A:41:ILE:HG12 | 2.17 | 0.45 |
| 1:A:144:MET:CE | 1:A:151:PRO:HG3 | 2.46 | 0.45 |
| 1:G:75:LEU:HB3 | 1:D:80:PHE:HE1 | 1.81 | 0.45 |
| 1:D:144:MET:CE | 1:D:151:PRO:HG3 | 2.46 | 0.45 |
| 1:C:37:ARG:O | 1:C:41:ILE:HG12 | 2.17 | 0.44 |
| 1:B:37:ARG:O | 1:B:41:ILE:HG12 | 2.17 | 0.44 |
| 1:A:250:ALA:O | 1:A:251:TYR:C | 2.56 | 0.44 |
| 1:D:250:ALA:O | 1:D:251:TYR:C | 2.56 | 0.44 |
| 1:D:192:LEU:HD23 | 1:D:256:VAL:HG22 | 2.00 | 0.44 |
| 1:A:192:LEU:HD23 | 1:A:256:VAL:HG22 | 2.00 | 0.44 |
| 1:A:8:ARG:NH2 | 1:A:276:VAL:O | 2.38 | 0.43 |
| 1:A:17:ILE:HG22 | 1:A:191:LEU:HB3 | 2.00 | 0.43 |
| 1:B:192:LEU:HD23 | 1:B:256:VAL:HG22 | 2.00 | 0.43 |
| 1:D:60:SER:HB2 | 1:D:176:ASN:ND2 | 2.33 | 0.43 |
| 1:D:17:ILE:HG22 | 1:D:191:LEU:HB3 | 2.00 | 0.43 |
| 1:C:174:TRP:CZ2 | 1:C:179:LEU:HD13 | 2.54 | 0.43 |
| 1:C:192:LEU:HD23 | 1:C:256:VAL:HG22 | 2.00 | 0.43 |
| 1:A:60:SER:HB2 | 1:A:176:ASN:ND2 | 2.33 | 0.43 |
| 1:B:174:TRP:CZ2 | 1:B:179:LEU:HD13 | 2.54 | 0.43 |

*Continued on next page...*

For Manuscript Review

*Continued from previous page...*

| Atom-1 | Atom-2 | Interatomic distance (Å) | Clash overlap (Å) |
|---|---|---|---|
| 1:D:8:ARG:NH2 | 1:D:276:VAL:O | 2.38 | 0.43 |
| 1:C:6:ARG:NH1 | 1:C:287:TRP:HB3 | 2.34 | 0.43 |
| 1:C:201:LEU:HB3 | 1:C:214:ILE:HG21 | 2.01 | 0.43 |
| 1:C:250:ALA:O | 1:C:251:TYR:C | 2.56 | 0.43 |
| 1:A:139:GLU:HA | 1:A:139:GLU:OE1 | 2.18 | 0.43 |
| 1:B:6:ARG:NH1 | 1:B:287:TRP:HB3 | 2.34 | 0.43 |
| 1:B:201:LEU:HB3 | 1:B:214:ILE:HG21 | 2.01 | 0.43 |
| 1:D:139:GLU:OE1 | 1:D:139:GLU:HA | 2.18 | 0.43 |
| 1:C:81:ALA:HA | 1:C:84:GLN:HE22 | 1.84 | 0.43 |
| 1:C:139:GLU:HA | 1:C:139:GLU:OE1 | 2.18 | 0.43 |
| 1:B:17:ILE:HG22 | 1:B:191:LEU:HB3 | 2.00 | 0.43 |
| 1:B:60:SER:HB2 | 1:B:176:ASN:ND2 | 2.33 | 0.43 |
| 1:C:17:ILE:HG22 | 1:C:191:LEU:HB3 | 2.00 | 0.43 |
| 1:C:60:SER:HB2 | 1:C:176:ASN:ND2 | 2.33 | 0.43 |
| 1:B:81:ALA:HA | 1:B:84:GLN:HE22 | 1.84 | 0.43 |
| 1:B:139:GLU:OE1 | 1:B:139:GLU:HA | 2.18 | 0.43 |
| 1:B:250:ALA:O | 1:B:251:TYR:C | 2.56 | 0.43 |
| 1:A:201:LEU:HB3 | 1:A:214:ILE:HG21 | 2.01 | 0.43 |
| 1:D:12:LYS:HA | 1:D:12:LYS:HD3 | 1.73 | 0.43 |
| 1:A:12:LYS:HA | 1:A:12:LYS:HD3 | 1.73 | 0.42 |
| 1:D:174:TRP:CZ2 | 1:D:179:LEU:HD13 | 2.54 | 0.42 |
| 1:D:201:LEU:HB3 | 1:D:214:ILE:HG21 | 2.01 | 0.42 |
| 1:A:174:TRP:CZ2 | 1:A:179:LEU:HD13 | 2.54 | 0.42 |
| 1:C:78:SER:O | 1:F:79:SER:HA | 2.20 | 0.42 |
| 1:B:78:SER:O | 1:G:79:SER:HA | 2.20 | 0.42 |
| 1:D:6:ARG:NH1 | 1:D:287:TRP:HB3 | 2.34 | 0.42 |
| 1:A:78:SER:O | 1:E:79:SER:HA | 2.20 | 0.42 |
| 1:B:121:MET:O | 1:B:153:PHE:HA | 2.20 | 0.42 |
| 1:C:121:MET:O | 1:C:153:PHE:HA | 2.20 | 0.42 |
| 1:A:6:ARG:NH1 | 1:A:287:TRP:HB3 | 2.34 | 0.42 |
| 1:D:78:SER:O | 1:H:79:SER:HA | 2.20 | 0.42 |
| 1:D:81:ALA:HA | 1:D:84:GLN:HE22 | 1.84 | 0.42 |
| 1:A:81:ALA:HA | 1:A:84:GLN:HE22 | 1.84 | 0.42 |
| 1:B:70:LEU:HD23 | 1:B:70:LEU:HA | 1.91 | 0.42 |
| 1:B:228:PHE:HE2 | 1:B:240:THR:HA | 1.84 | 0.42 |
| 1:D:228:PHE:HE2 | 1:D:240:THR:HA | 1.84 | 0.42 |
| 1:C:118:VAL:HG12 | 1:C:150:ILE:HB | 2.02 | 0.41 |
| 1:A:149:VAL:O | 1:A:151:PRO:HD3 | 2.20 | 0.41 |
| 1:A:228:PHE:HE2 | 1:A:240:THR:HA | 1.84 | 0.41 |
| 1:C:228:PHE:HE2 | 1:C:240:THR:HA | 1.84 | 0.41 |
| 1:F:79:SER:O | 1:F:82:ASP:N | 2.53 | 0.41 |

*Continued on next page...*

For Manuscript Review

*Continued from previous page...*

| Atom-1 | Atom-2 | Interatomic distance (Å) | Clash overlap (Å) |
|---|---|---|---|
| 1:B:118:VAL:HG12 | 1:B:150:ILE:HB | 2.02 | 0.41 |
| 1:B:149:VAL:O | 1:B:151:PRO:HD3 | 2.20 | 0.41 |
| 1:D:149:VAL:O | 1:D:151:PRO:HD3 | 2.20 | 0.41 |
| 1:C:70:LEU:HD23 | 1:C:70:LEU:HA | 1.91 | 0.41 |
| 1:C:149:VAL:O | 1:C:151:PRO:HD3 | 2.20 | 0.41 |
| 1:B:80:PHE:HE1 | 1:H:75:LEU:HB3 | 1.81 | 0.41 |
| 1:G:79:SER:O | 1:G:82:ASP:N | 2.53 | 0.41 |
| 1:C:80:PHE:HE1 | 1:E:75:LEU:HB3 | 1.81 | 0.41 |
| 1:A:242:LYS:HE2 | 1:A:242:LYS:HB2 | 1.87 | 0.41 |
| 1:C:37:ARG:HD2 | 1:C:262:TYR:CD1 | 2.56 | 0.41 |
| 1:B:37:ARG:HD2 | 1:B:262:TYR:CD1 | 2.56 | 0.41 |
| 1:C:239:MET:HG3 | 2:C:801:ADP:C4 | 2.56 | 0.41 |
| 1:B:239:MET:HG3 | 2:B:801:ADP:C4 | 2.56 | 0.41 |
| 1:A:67:ARG:HA | 1:A:95:CYS:SG | 2.61 | 0.41 |
| 1:A:121:MET:O | 1:A:153:PHE:HA | 2.20 | 0.41 |
| 1:D:37:ARG:HD2 | 1:D:262:TYR:CD1 | 2.56 | 0.41 |
| 1:D:67:ARG:HA | 1:D:95:CYS:SG | 2.61 | 0.41 |
| 1:A:37:ARG:HD2 | 1:A:262:TYR:CD1 | 2.56 | 0.41 |
| 1:D:121:MET:O | 1:D:153:PHE:HA | 2.20 | 0.41 |
| 1:D:242:LYS:HE2 | 1:D:242:LYS:HB2 | 1.87 | 0.41 |
| 1:A:288:ALA:HA | 1:A:289:PRO:HD3 | 1.98 | 0.40 |
| 1:A:118:VAL:HG12 | 1:A:150:ILE:HB | 2.02 | 0.40 |
| 1:E:79:SER:O | 1:E:82:ASP:N | 2.53 | 0.40 |
| 1:C:159:ILE:HD13 | 1:D:122:LEU:O | 2.21 | 0.40 |
| 1:A:122:LEU:O | 1:B:159:ILE:HD13 | 2.21 | 0.40 |
| 1:D:118:VAL:HG12 | 1:D:150:ILE:HB | 2.02 | 0.40 |
| 1:D:288:ALA:HA | 1:D:289:PRO:HD3 | 1.98 | 0.40 |
| 1:H:79:SER:O | 1:H:82:ASP:N | 2.53 | 0.40 |
| 1:A:239:MET:HG3 | 2:A:801:ADP:C4 | 2.56 | 0.40 |

There are no symmetry-related clashes.

#### 5.3 Torsion angles (i)

##### 5.3.1 Protein backbone (i)

In the following table, the Percentiles column shows the percent Ramachandran outliers of the chain as a percentile score with respect to all PDB entries followed by that with respect to all EM entries.

The Analysed column shows the number of residues for which the backbone conformation was analysed, and the total number of residues.

For Manuscript Review

| Mol | Chain | Analysed | Favoured | Allowed | Outliers | Percentiles | |
|---|---|---|---|---|---|---|---|
| 1 | A | 264/726 (36%) | 259 (98%) | 5 (2%) | 0 | 100 | 100 |
| 1 | B | 264/726 (36%) | 259 (98%) | 5 (2%) | 0 | 100 | 100 |
| 1 | C | 264/726 (36%) | 259 (98%) | 5 (2%) | 0 | 100 | 100 |
| 1 | D | 264/726 (36%) | 259 (98%) | 5 (2%) | 0 | 100 | 100 |
| 1 | E | 13/726 (2%) | 13 (100%) | 0 | 0 | 100 | 100 |
| 1 | F | 13/726 (2%) | 13 (100%) | 0 | 0 | 100 | 100 |
| 1 | G | 13/726 (2%) | 13 (100%) | 0 | 0 | 100 | 100 |
| 1 | H | 13/726 (2%) | 13 (100%) | 0 | 0 | 100 | 100 |
| All | All | 1108/5808 (19%) | 1088 (98%) | 20 (2%) | 0 | 100 | 100 |

There are no Ramachandran outliers to report.

##### 5.3.2 Protein sidechains (i)

In the following table, the Percentiles column shows the percent sidechain outliers of the chain as a percentile score with respect to all PDB entries followed by that with respect to all EM entries.

The Analysed column shows the number of residues for which the sidechain conformation was analysed, and the total number of residues.

| Mol | Chain | Analysed | Rotameric | Outliers | Percentiles | |
|---|---|---|---|---|---|---|
| 1 | A | 221/601 (37%) | 211 (96%) | 10 (4%) | 27 | 58 |
| 1 | B | 221/601 (37%) | 211 (96%) | 10 (4%) | 27 | 58 |
| 1 | C | 221/601 (37%) | 211 (96%) | 10 (4%) | 27 | 58 |
| 1 | D | 221/601 (37%) | 211 (96%) | 10 (4%) | 27 | 58 |
| 1 | E | 14/601 (2%) | 13 (93%) | 1 (7%) | 14 | 42 |
| 1 | F | 14/601 (2%) | 13 (93%) | 1 (7%) | 14 | 42 |
| 1 | G | 14/601 (2%) | 13 (93%) | 1 (7%) | 14 | 42 |
| 1 | H | 14/601 (2%) | 13 (93%) | 1 (7%) | 14 | 42 |
| All | All | 940/4808 (20%) | 896 (95%) | 44 (5%) | 30 | 57 |

All (44) residues with a non-rotameric sidechain are listed below:

| Mol | Chain | Res | Type |
|---|---|---|---|
| 1 | C | 6 | ARG |
| 1 | C | 10 | PHE |

*Continued on next page...*

For Manuscript Review

*Continued from previous page...*

| Mol | Chain | Res | Type |
|---|---|---|---|
| 1 | C | 29 | LYS |
| 1 | C | 38 | LEU |
| 1 | C | 107 | TYR |
| 1 | C | 108 | GLU |
| 1 | C | 175 | ASP |
| 1 | C | 223 | GLN |
| 1 | C | 239 | MET |
| 1 | C | 281 | HIS |
| 1 | F | 84 | GLN |
| 1 | A | 6 | ARG |
| 1 | A | 10 | PHE |
| 1 | A | 29 | LYS |
| 1 | A | 38 | LEU |
| 1 | A | 107 | TYR |
| 1 | A | 108 | GLU |
| 1 | A | 175 | ASP |
| 1 | A | 223 | GLN |
| 1 | A | 239 | MET |
| 1 | A | 281 | HIS |
| 1 | E | 84 | GLN |
| 1 | B | 6 | ARG |
| 1 | B | 10 | PHE |
| 1 | B | 29 | LYS |
| 1 | B | 38 | LEU |
| 1 | B | 107 | TYR |
| 1 | B | 108 | GLU |
| 1 | B | 175 | ASP |
| 1 | B | 223 | GLN |
| 1 | B | 239 | MET |
| 1 | B | 281 | HIS |
| 1 | G | 84 | GLN |
| 1 | D | 6 | ARG |
| 1 | D | 10 | PHE |
| 1 | D | 29 | LYS |
| 1 | D | 38 | LEU |
| 1 | D | 107 | TYR |
| 1 | D | 108 | GLU |
| 1 | D | 175 | ASP |
| 1 | D | 223 | GLN |
| 1 | D | 239 | MET |
| 1 | D | 281 | HIS |
| 1 | H | 84 | GLN |

For Manuscript Review

Sometimes sidechains can be flipped to improve hydrogen bonding and reduce clashes. All (9) such sidechains are listed below:

| Mol | Chain | Res | Type |
|---|---|---|---|
| 1 | C | 113 | GLN |
| 1 | C | 281 | HIS |
| 1 | A | 113 | GLN |
| 1 | A | 281 | HIS |
| 1 | B | 113 | GLN |
| 1 | B | 281 | HIS |
| 1 | D | 113 | GLN |
| 1 | D | 222 | HIS |
| 1 | D | 281 | HIS |

##### 5.3.3 RNA (i)

There are no RNA molecules in this entry.

#### 5.4 Non-standard residues in protein, DNA, RNA chains (i)

There are no non-standard protein/DNA/RNA residues in this entry.

#### 5.5 Carbohydrates (i)

There are no monosaccharides in this entry.

#### 5.6 Ligand geometry (i)

Of 12 ligands modelled in this entry, 4 are monoatomic - leaving 8 for Mogul analysis.

In the following table, the Counts columns list the number of bonds (or angles) for which Mogul statistics could be retrieved, the number of bonds (or angles) that are observed in the model and the number of bonds (or angles) that are defined in the Chemical Component Dictionary. The Link column lists molecule types, if any, to which the group is linked. The Z score for a bond length (or angle) is the number of standard deviations the observed value is removed from the expected value. A bond length (or angle) with $|Z| > 2$ is considered an outlier worth inspection. RMSZ is the root-mean-square of all Z scores of the bond lengths (or angles).

| Mol | Type | Chain | Res | Link | Bond lengths | | | Bond angles | | |
|---|---|---|---|---|---|---|---|---|---|---|
| | | | | | Counts | RMSZ | $\#\|Z\| > 2$ | Counts | RMSZ | $\#\|Z\| > 2$ |
| 2 | ADP | B | 801 | 4 | 24,29,29 | 0.96 | 1 (4%) | 29,45,45 | 1.43 | 4 (13%) |
| 2 | ADP | D | 801 | 4 | 24,29,29 | 0.96 | 1 (4%) | 29,45,45 | 1.43 | 4 (13%) |
| 3 | RGP | D | 802 | - | 11,13,13 | 0.88 | 0 | 14,18,18 | 1.09 | 0 |

For Manuscript Review

| Mol | Type | Chain | Res | Link | Bond lengths | | | Bond angles | | |
|---|---|---|---|---|---|---|---|---|---|---|
| | | | | | Counts | RMSZ | #\|Z\| > 2 | Counts | RMSZ | #\|Z\| > 2 |
| 2 | ADP | A | 801 | 4 | 24,29,29 | 0.96 | 1 (4%) | 29,45,45 | 1.43 | 4 (13%) |
| 3 | RGP | A | 802 | - | 11,13,13 | 0.89 | 0 | 14,18,18 | 1.13 | 0 |
| 3 | RGP | B | 802 | - | 11,13,13 | 0.88 | 0 | 14,18,18 | 1.09 | 0 |
| 3 | RGP | C | 802 | - | 11,13,13 | 0.89 | 0 | 14,18,18 | 1.13 | 0 |
| 2 | ADP | C | 801 | 4 | 24,29,29 | 0.96 | 1 (4%) | 29,45,45 | 1.43 | 4 (13%) |

In the following table, the Chirals column lists the number of chiral outliers, the number of chiral centers analysed, the number of these observed in the model and the number defined in the Chemical Component Dictionary. Similar counts are reported in the Torsion and Rings columns. '-' means no outliers of that kind were identified.

| Mol | Type | Chain | Res | Link | Chirals | Torsions | Rings |
|---|---|---|---|---|---|---|---|
| 2 | ADP | B | 801 | 4 | - | 3/12/32/32 | 0/3/3/3 |
| 2 | ADP | D | 801 | 4 | - | 3/12/32/32 | 0/3/3/3 |
| 3 | RGP | D | 802 | - | - | 4/12/14/14 | - |
| 2 | ADP | A | 801 | 4 | - | 4/12/32/32 | 0/3/3/3 |
| 3 | RGP | A | 802 | - | - | 2/12/14/14 | - |
| 3 | RGP | B | 802 | - | - | 4/12/14/14 | - |
| 3 | RGP | C | 802 | - | - | 2/12/14/14 | - |
| 2 | ADP | C | 801 | 4 | - | 4/12/32/32 | 0/3/3/3 |

All (4) bond length outliers are listed below:

| Mol | Chain | Res | Type | Atoms | Z | Observed(Å) | Ideal(Å) |
|---|---|---|---|---|---|---|---|
| 2 | C | 801 | ADP | C5-C4 | 2.42 | 1.47 | 1.40 |
| 2 | A | 801 | ADP | C5-C4 | 2.42 | 1.47 | 1.40 |
| 2 | B | 801 | ADP | C5-C4 | 2.42 | 1.47 | 1.40 |
| 2 | D | 801 | ADP | C5-C4 | 2.42 | 1.47 | 1.40 |

All (16) bond angle outliers are listed below:

| Mol | Chain | Res | Type | Atoms | Z | Observed($^o$) | Ideal($^o$) |
|---|---|---|---|---|---|---|---|
| 2 | C | 801 | ADP | PA-O3A-PB | -3.52 | 120.76 | 132.83 |
| 2 | A | 801 | ADP | PA-O3A-PB | -3.52 | 120.76 | 132.83 |
| 2 | B | 801 | ADP | PA-O3A-PB | -3.52 | 120.76 | 132.83 |
| 2 | D | 801 | ADP | PA-O3A-PB | -3.52 | 120.76 | 132.83 |
| 2 | C | 801 | ADP | N3-C2-N1 | -3.17 | 123.73 | 128.68 |
| 2 | A | 801 | ADP | N3-C2-N1 | -3.17 | 123.73 | 128.68 |
| 2 | B | 801 | ADP | N3-C2-N1 | -3.17 | 123.73 | 128.68 |
| 2 | D | 801 | ADP | N3-C2-N1 | -3.17 | 123.73 | 128.68 |

*Continued on next page...*

For Manuscript Review

*Continued from previous page...*

| Mol | Chain | Res | Type | Atoms | Z | Observed($^o$) | Ideal($^o$) |
|---|---|---|---|---|---|---|---|
| 2 | C | 801 | ADP | C3'-C2'-C1' | 2.95 | 105.42 | 100.98 |
| 2 | A | 801 | ADP | C3'-C2'-C1' | 2.95 | 105.42 | 100.98 |
| 2 | B | 801 | ADP | C3'-C2'-C1' | 2.95 | 105.42 | 100.98 |
| 2 | D | 801 | ADP | C3'-C2'-C1' | 2.95 | 105.42 | 100.98 |
| 2 | C | 801 | ADP | C4-C5-N7 | -2.54 | 106.75 | 109.40 |
| 2 | A | 801 | ADP | C4-C5-N7 | -2.54 | 106.75 | 109.40 |
| 2 | B | 801 | ADP | C4-C5-N7 | -2.54 | 106.75 | 109.40 |
| 2 | D | 801 | ADP | C4-C5-N7 | -2.54 | 106.75 | 109.40 |

There are no chirality outliers.

All (26) torsion outliers are listed below:

| Mol | Chain | Res | Type | Atoms |
|---|---|---|---|---|
| 2 | C | 801 | ADP | PA-O3A-PB-O3B |
| 2 | A | 801 | ADP | PA-O3A-PB-O3B |
| 3 | C | 802 | RGP | OXT-C-CA-N |
| 3 | A | 802 | RGP | OXT-C-CA-N |
| 3 | B | 802 | RGP | OXT-C-CA-N |
| 3 | D | 802 | RGP | OXT-C-CA-N |
| 3 | B | 802 | RGP | O-C-CA-N |
| 3 | D | 802 | RGP | O-C-CA-N |
| 3 | C | 802 | RGP | O-C-CA-N |
| 3 | A | 802 | RGP | O-C-CA-N |
| 3 | B | 802 | RGP | O-C-CA-CB |
| 3 | D | 802 | RGP | O-C-CA-CB |
| 2 | C | 801 | ADP | C4'-C5'-O5'-PA |
| 2 | A | 801 | ADP | C4'-C5'-O5'-PA |
| 2 | B | 801 | ADP | C4'-C5'-O5'-PA |
| 2 | D | 801 | ADP | C4'-C5'-O5'-PA |
| 3 | B | 802 | RGP | OXT-C-CA-CB |
| 3 | D | 802 | RGP | OXT-C-CA-CB |
| 2 | C | 801 | ADP | PA-O3A-PB-O1B |
| 2 | A | 801 | ADP | PA-O3A-PB-O1B |
| 2 | B | 801 | ADP | PA-O3A-PB-O1B |
| 2 | D | 801 | ADP | PA-O3A-PB-O1B |
| 2 | C | 801 | ADP | PA-O3A-PB-O2B |
| 2 | A | 801 | ADP | PA-O3A-PB-O2B |
| 2 | B | 801 | ADP | PA-O3A-PB-O2B |
| 2 | D | 801 | ADP | PA-O3A-PB-O2B |

There are no ring outliers.

8 monomers are involved in 15 short contacts:

For Manuscript Review

| Mol | Chain | Res | Type | Clashes | Symm-Clashes |
|---|---|---|---|---|---|
| 2 | B | 801 | ADP | 2 | 0 |
| 2 | D | 801 | ADP | 1 | 0 |
| 3 | D | 802 | RGP | 2 | 0 |
| 2 | A | 801 | ADP | 2 | 0 |
| 3 | A | 802 | RGP | 2 | 0 |
| 3 | B | 802 | RGP | 2 | 0 |
| 3 | C | 802 | RGP | 2 | 0 |
| 2 | C | 801 | ADP | 2 | 0 |

The following is a two-dimensional graphical depiction of Mogul quality analysis of bond lengths, bond angles, torsion angles, and ring geometry for all instances of the Ligand of Interest. In addition, ligands with molecular weight > 250 and outliers as shown on the validation Tables will also be included. For torsion angles, if less then 5% of the Mogul distribution of torsion angles is within 10 degrees of the torsion angle in question, then that torsion angle is considered an outlier. Any bond that is central to one or more torsion angles identified as an outlier by Mogul will be highlighted in the graph. For rings, the root-mean-square deviation (RMSD) between the ring in question and similar rings identified by Mogul is calculated over all ring torsion angles. If the average RMSD is greater than 60 degrees and the minimal RMSD between the ring in question and any Mogul-identified rings is also greater than 60 degrees, then that ring is considered an outlier. The outliers are highlighted in purple. The color gray indicates Mogul did not find sufficient equivalents in the CSD to analyse the geometry.

**Ligand ADP B 801**

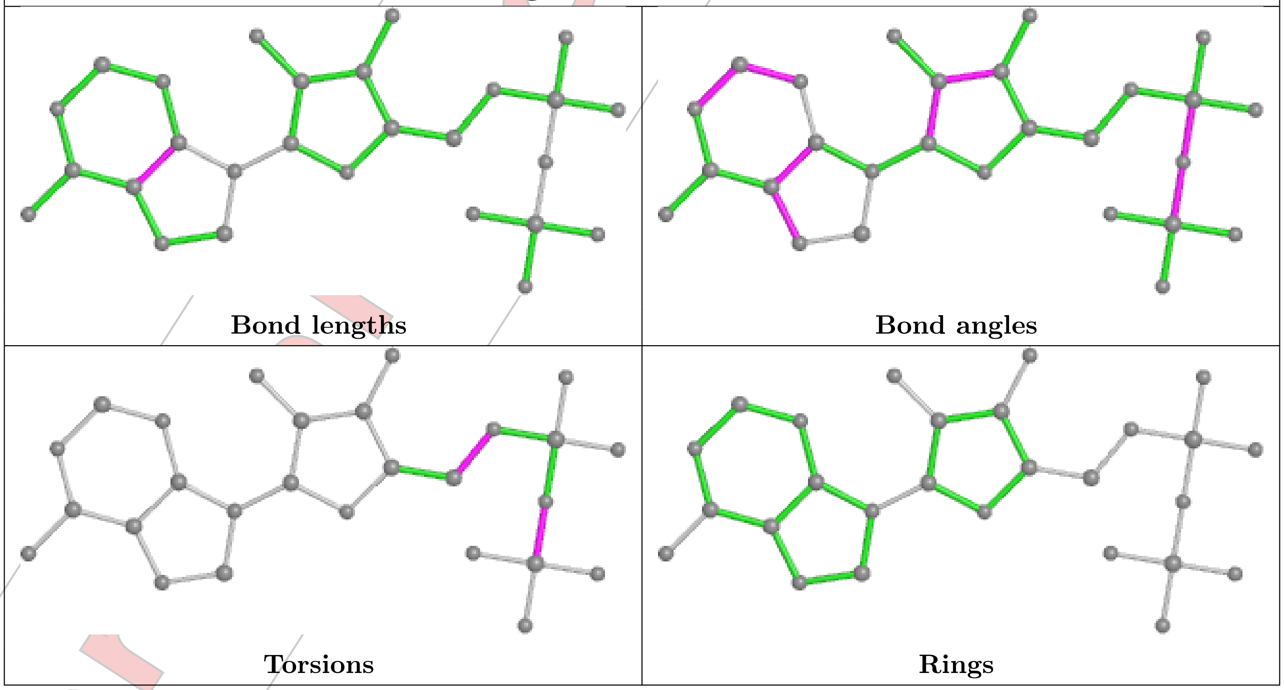


For Manuscript Review

##### Ligand ADP D 801

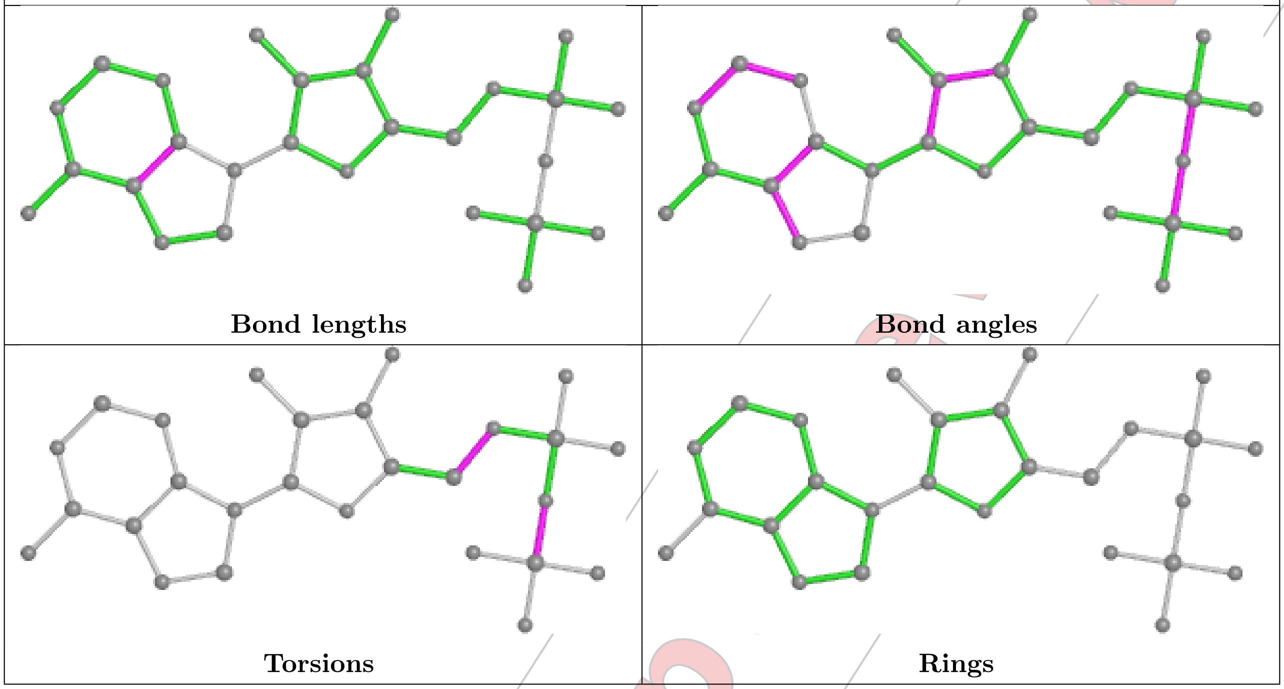


##### Ligand RGP D 802

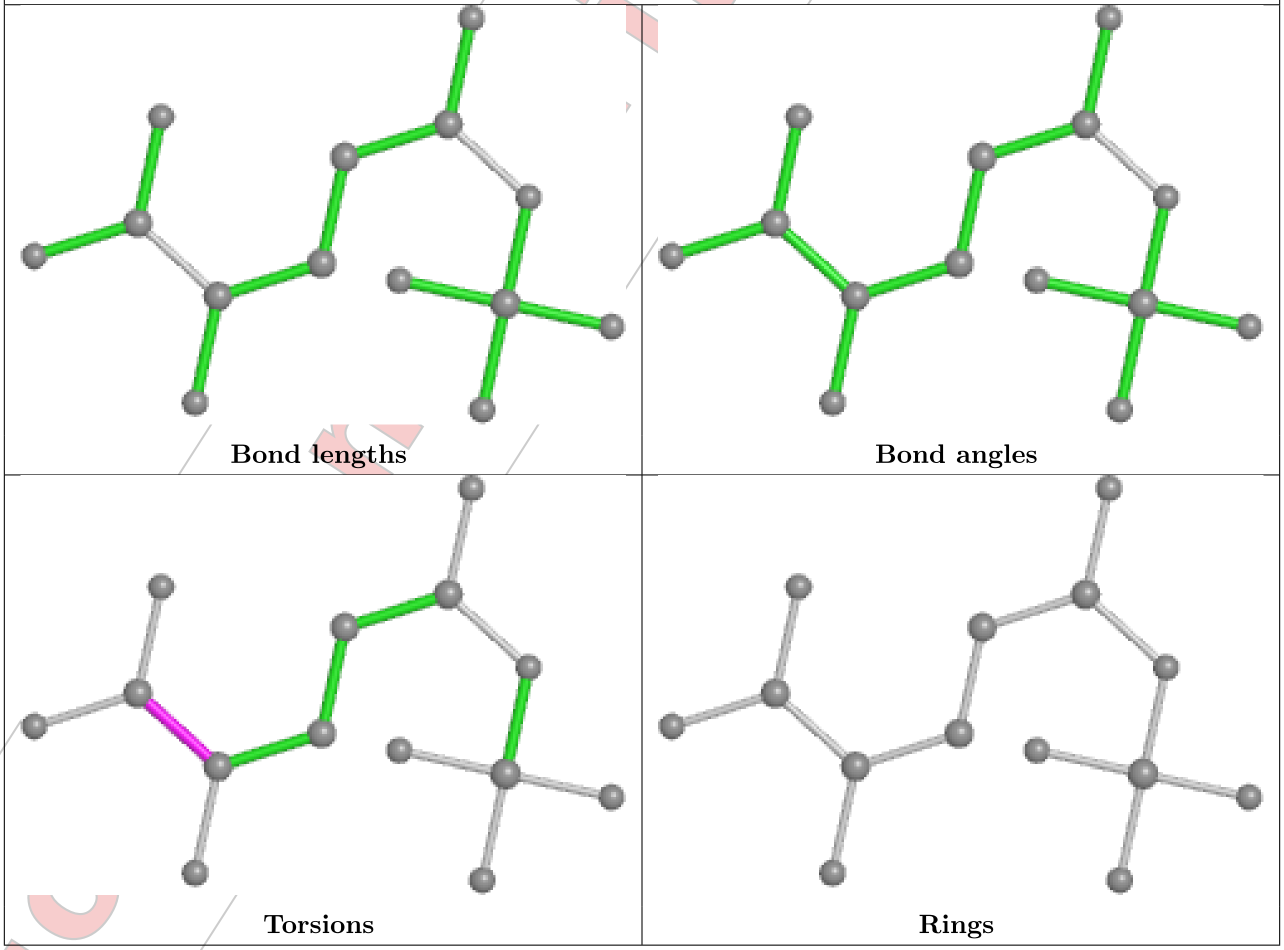

Ligand ADP A 801

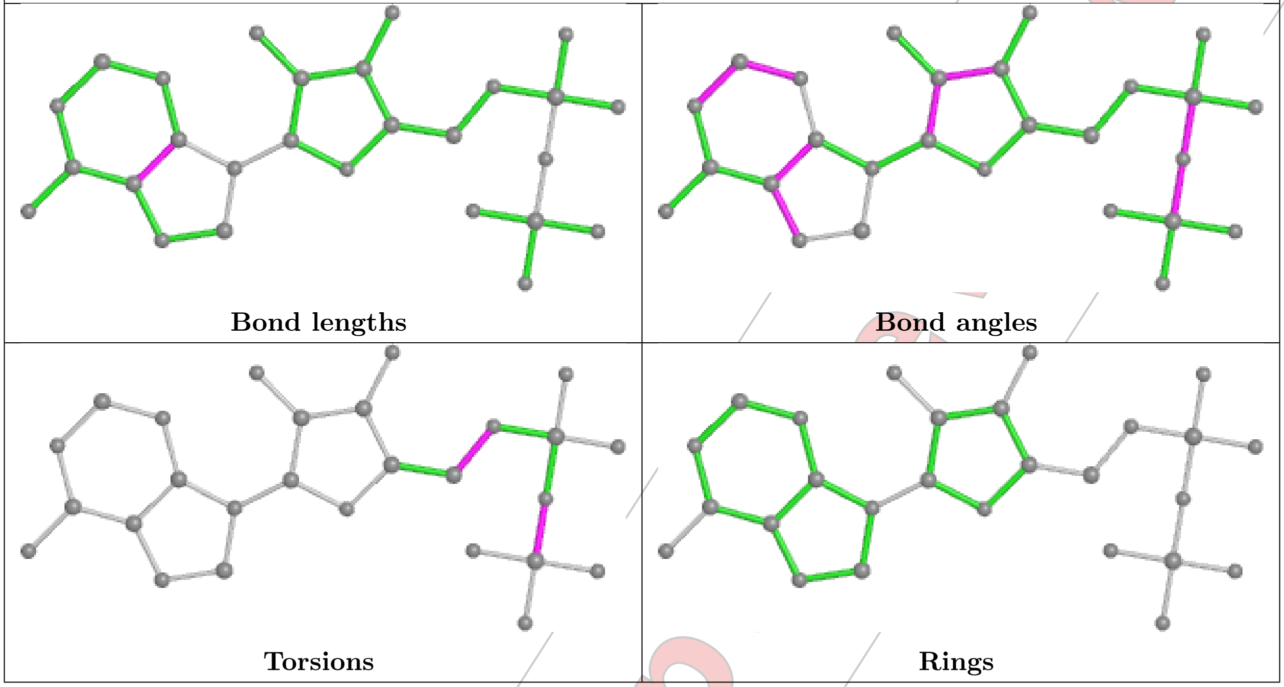


Ligand RGP A 802

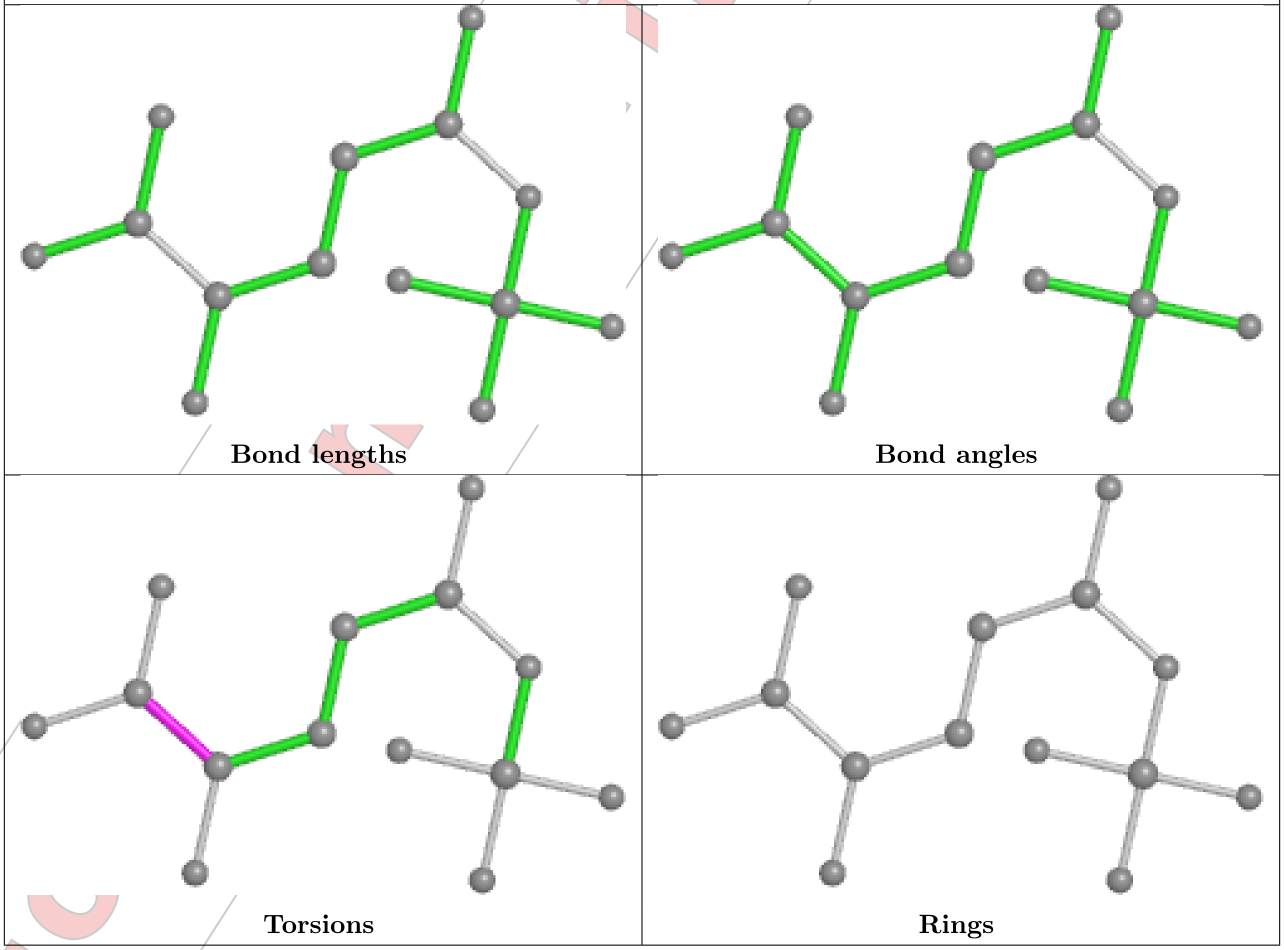

**Ligand RGP B 802**

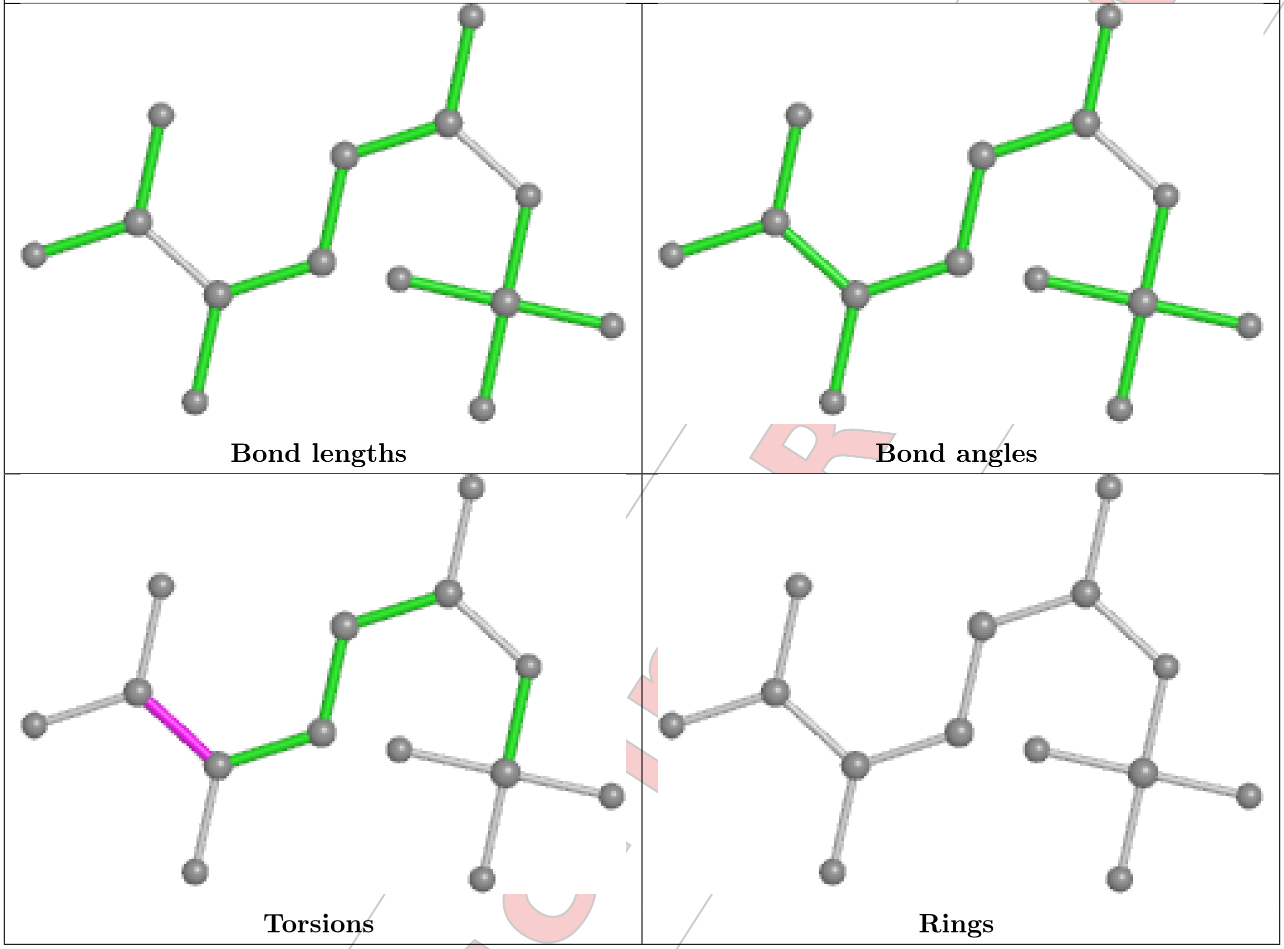


For Manuscript Review

##### Ligand RGP C 802

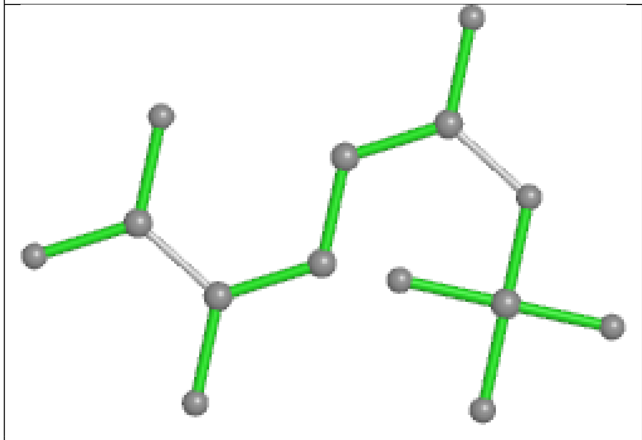
Bond lengths

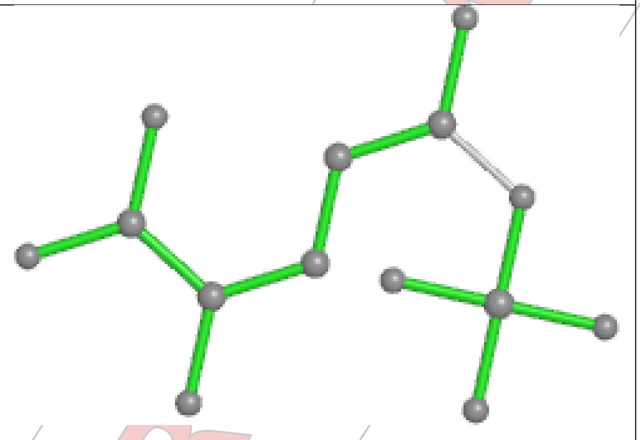
Bond angles

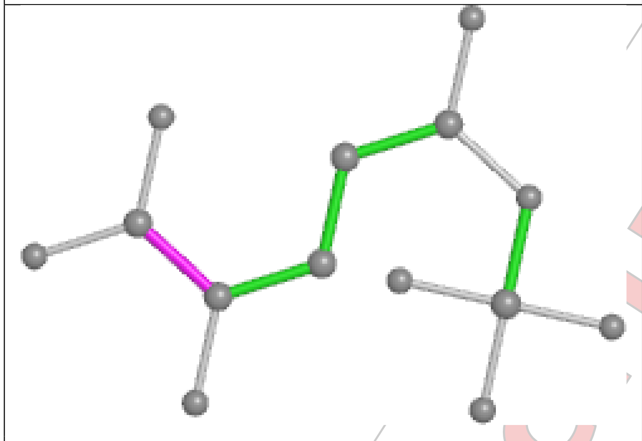
Torsions

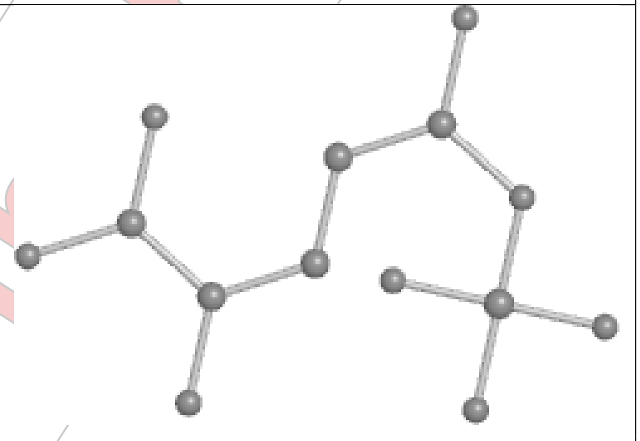
Rings

##### Ligand ADP C 801

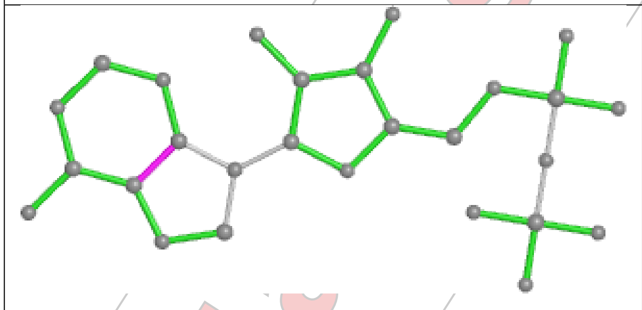
Bond lengths

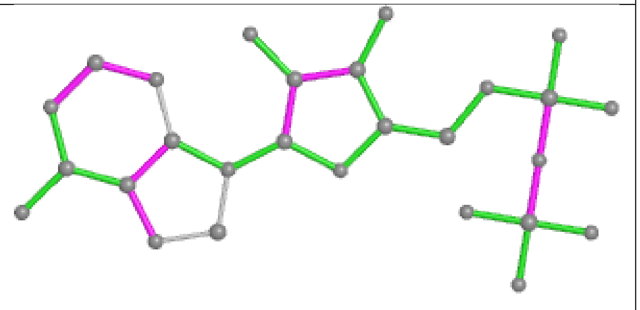
Bond angles

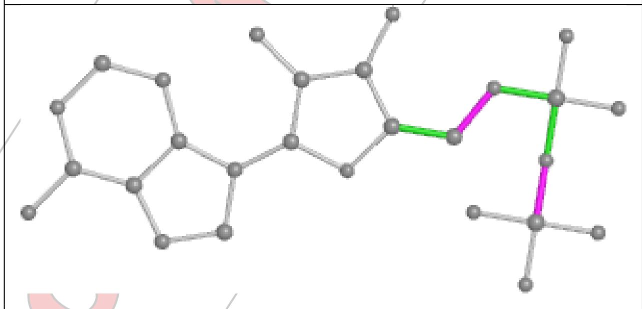
Torsions

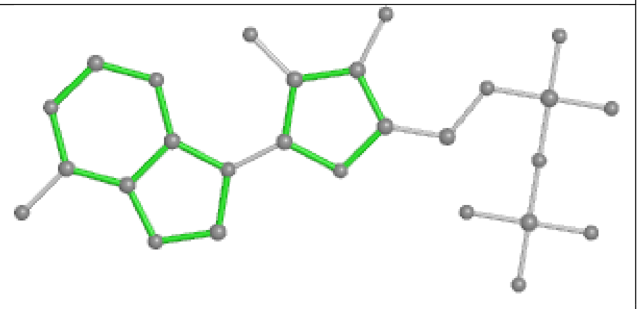
Rings

#### 5.7 Other polymers (i)

There are no such residues in this entry.

#### 5.8 Polymer linkage issues (i)

There are no chain breaks in this entry.

For Manuscript Review

### 6 Map visualisation (i)

This section contains visualisations of the EMDB entry EMD-35901. These allow visual inspection of the internal detail of the map and identification of artifacts.

Images derived from a raw map, generated by summing the deposited half-maps, are presented below the corresponding image components of the primary map to allow further visual inspection and comparison with those of the primary map.

#### 6.1 Orthogonal projections (i)

##### 6.1.1 Primary map

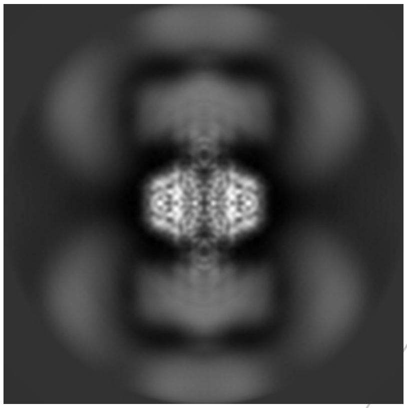
X

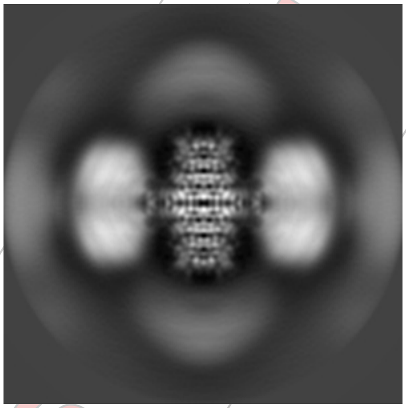
Y

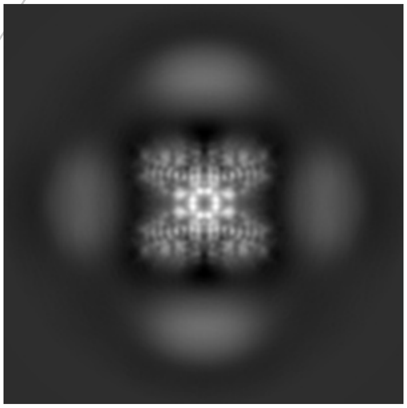
Z

##### 6.1.2 Raw map

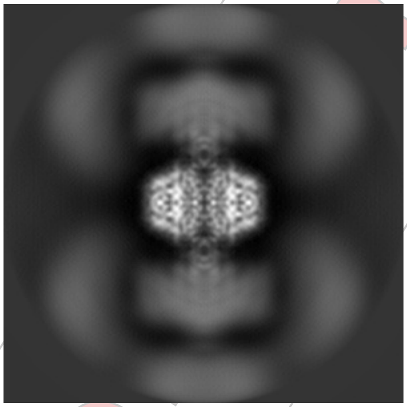
X

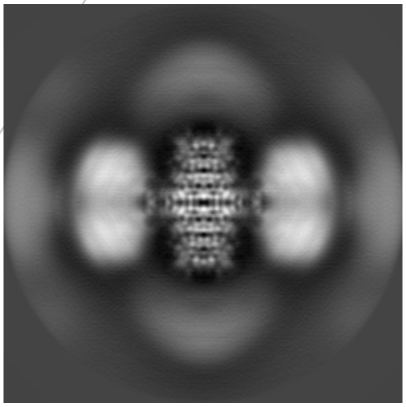
Y

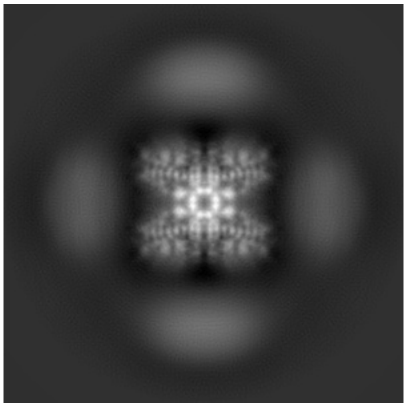
Z

The images above show the map projected in three orthogonal directions.

For Manuscript Review

#### 6.2 Central slices (i)

##### 6.2.1 Primary map

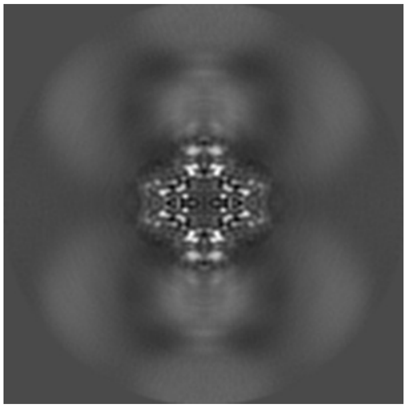

X Index: 120

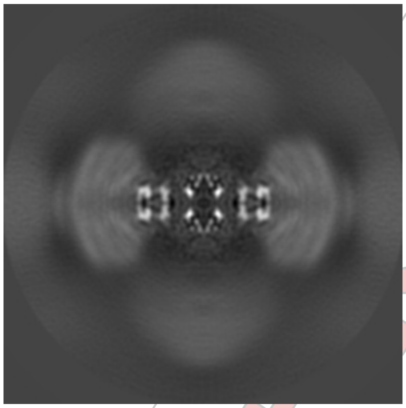

Y Index: 120

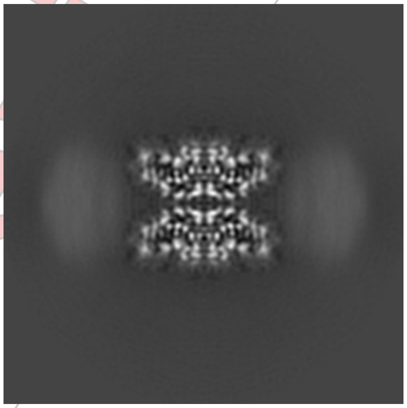

Z Index: 120

##### 6.2.2 Raw map

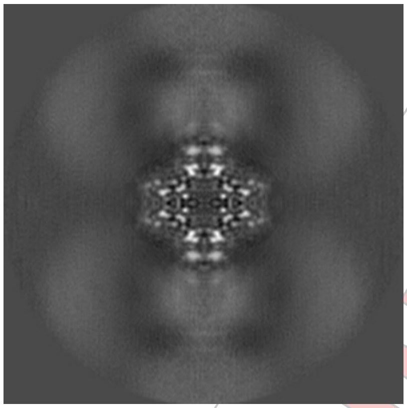

X Index: 120

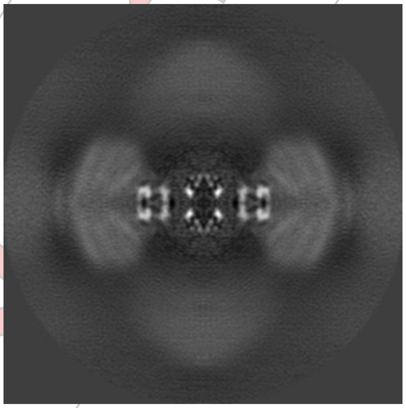

Y Index: 120

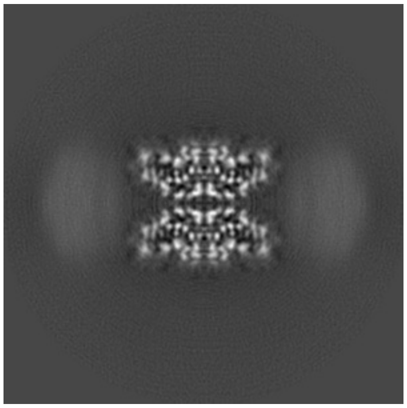

Z Index: 120

The images above show central slices of the map in three orthogonal directions.

For Manuscript Review

#### 6.3 Largest variance slices (i)

##### 6.3.1 Primary map

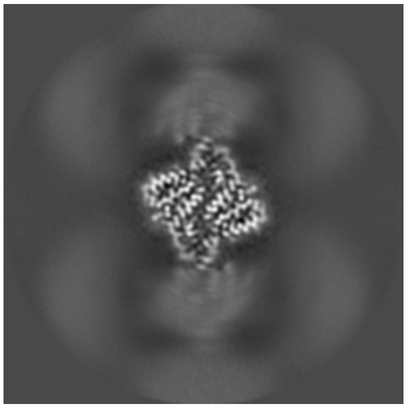
X Index: 116

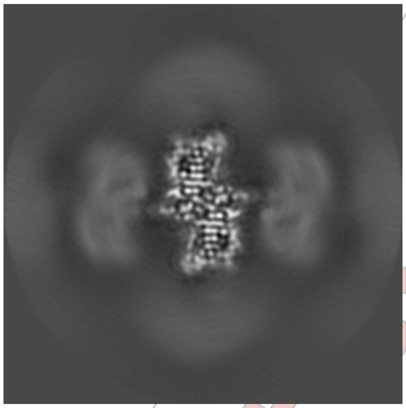
Y Index: 104

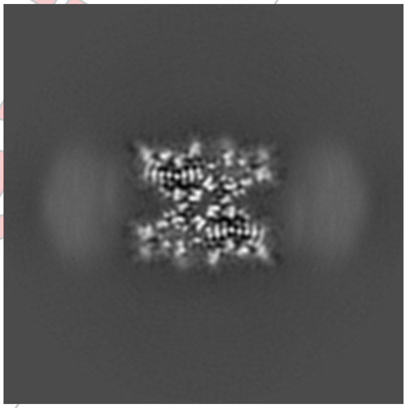
Z Index: 116

##### 6.3.2 Raw map

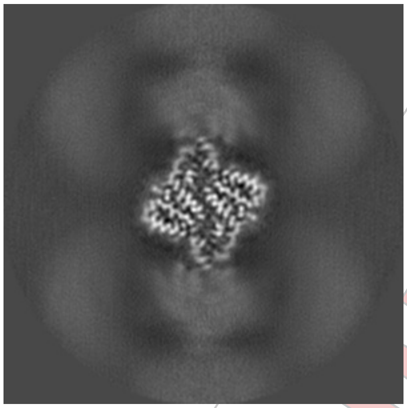
X Index: 124

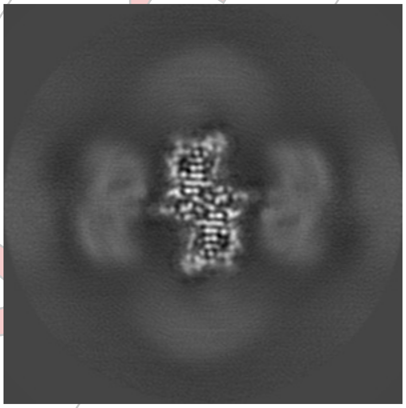
Y Index: 104

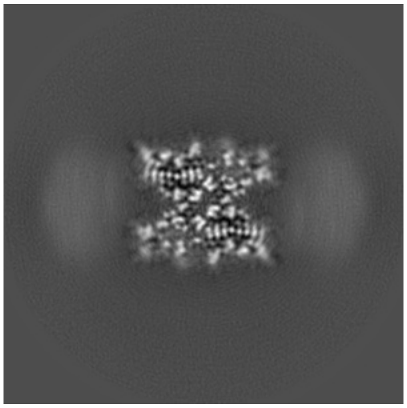
Z Index: 116

The images above show the largest variance slices of the map in three orthogonal directions.

For Manuscript Review

#### 6.4 Orthogonal standard-deviation projections (False-color) (i)

##### 6.4.1 Primary map

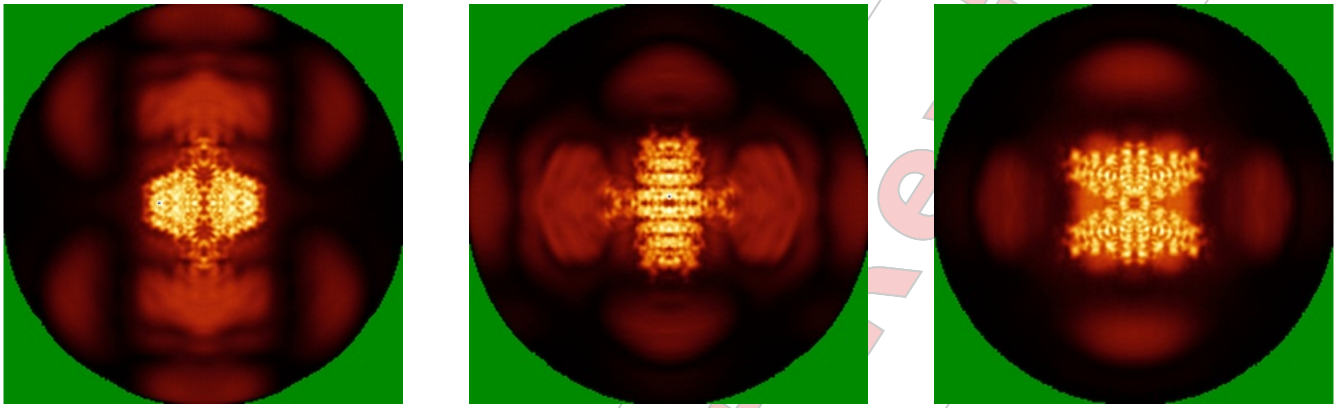

X Y Z

##### 6.4.2 Raw map

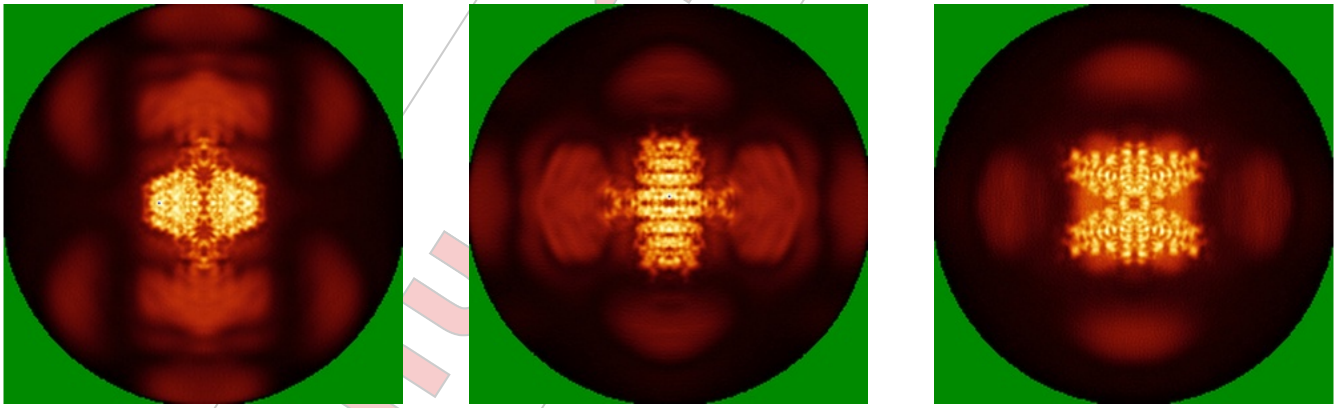

X Y Z

The images above show the map standard deviation projections with false color in three orthogonal directions. Minimum values are shown in green, max in blue, and dark to light orange shades represent small to large values respectively.

For Manuscript Review

#### 6.5 Orthogonal surface views (i)

##### 6.5.1 Primary map

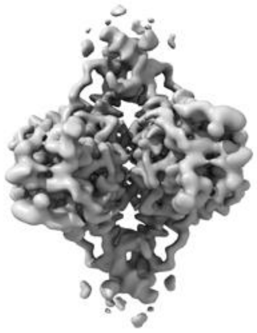

X

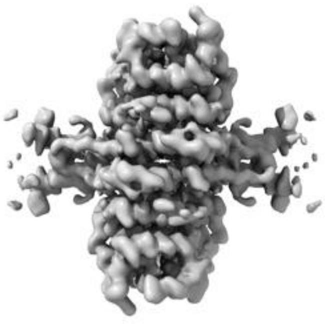

Y

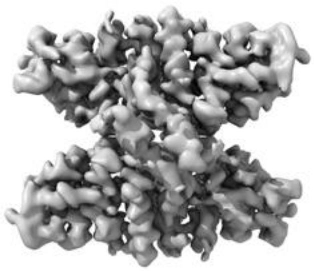

Z

The images above show the 3D surface view of the map at the recommended contour level 0.005. These images, in conjunction with the slice images, may facilitate assessment of whether an appropriate contour level has been provided.

##### 6.5.2 Raw map

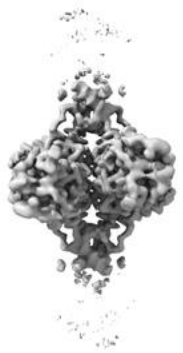

X

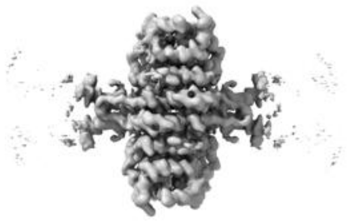

Y

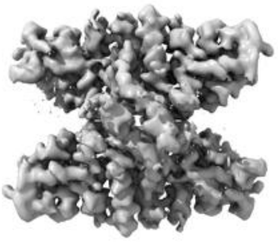

Z

These images show the 3D surface of the raw map. The raw map's contour level was selected so that its surface encloses the same volume as the primary map does at its recommended contour level.

#### 6.6 Mask visualisation (i)

This section was not generated. No masks/segmentation were deposited.

For Manuscript Review

### 7 Map analysis (i)

This section contains the results of statistical analysis of the map.

#### 7.1 Map-value distribution (i)

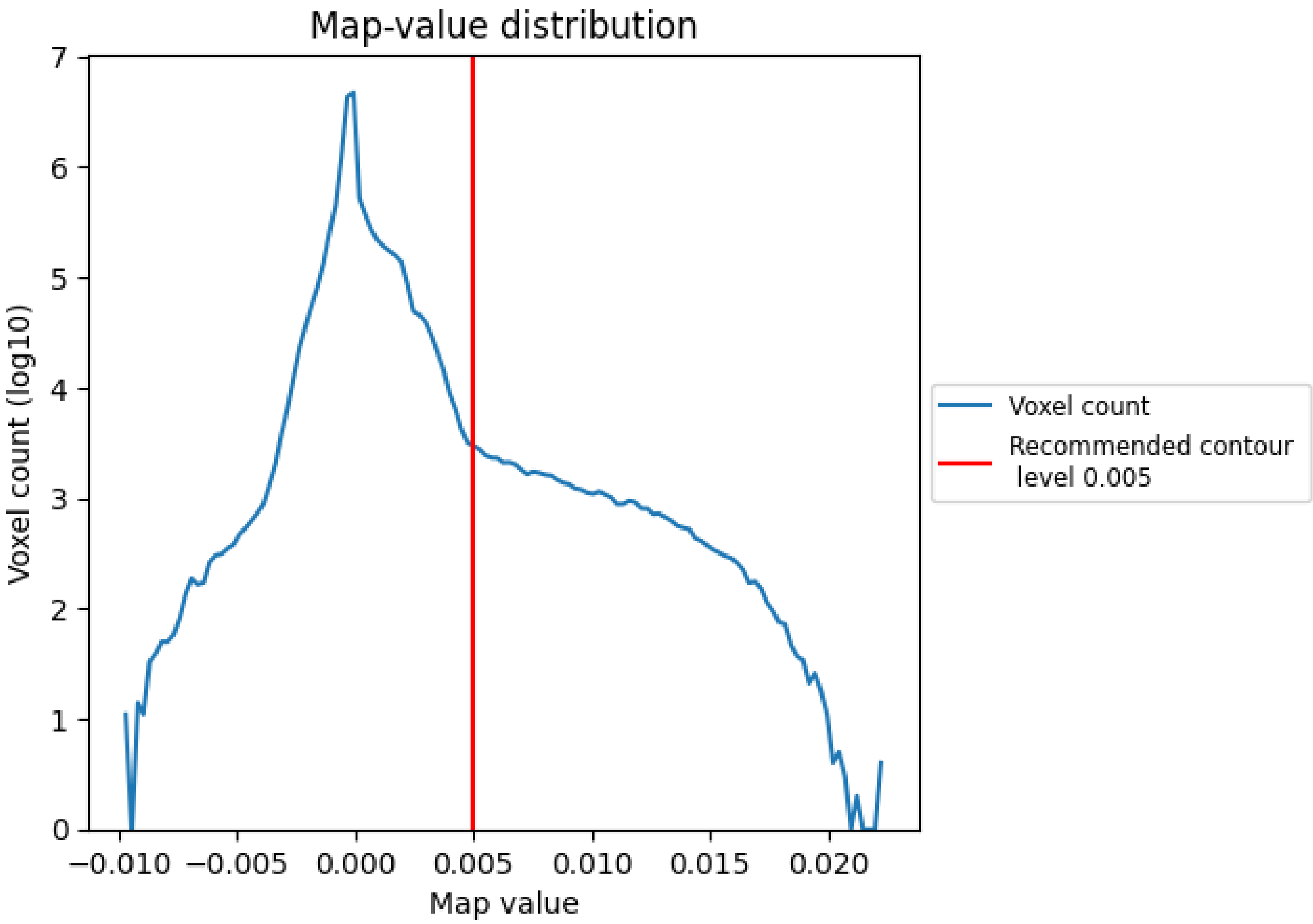


The map-value distribution is plotted in 128 intervals along the x-axis. The y-axis is logarithmic. A spike in this graph at zero usually indicates that the volume has been masked.

For Manuscript Review

#### 7.2 Volume estimate ⓘ

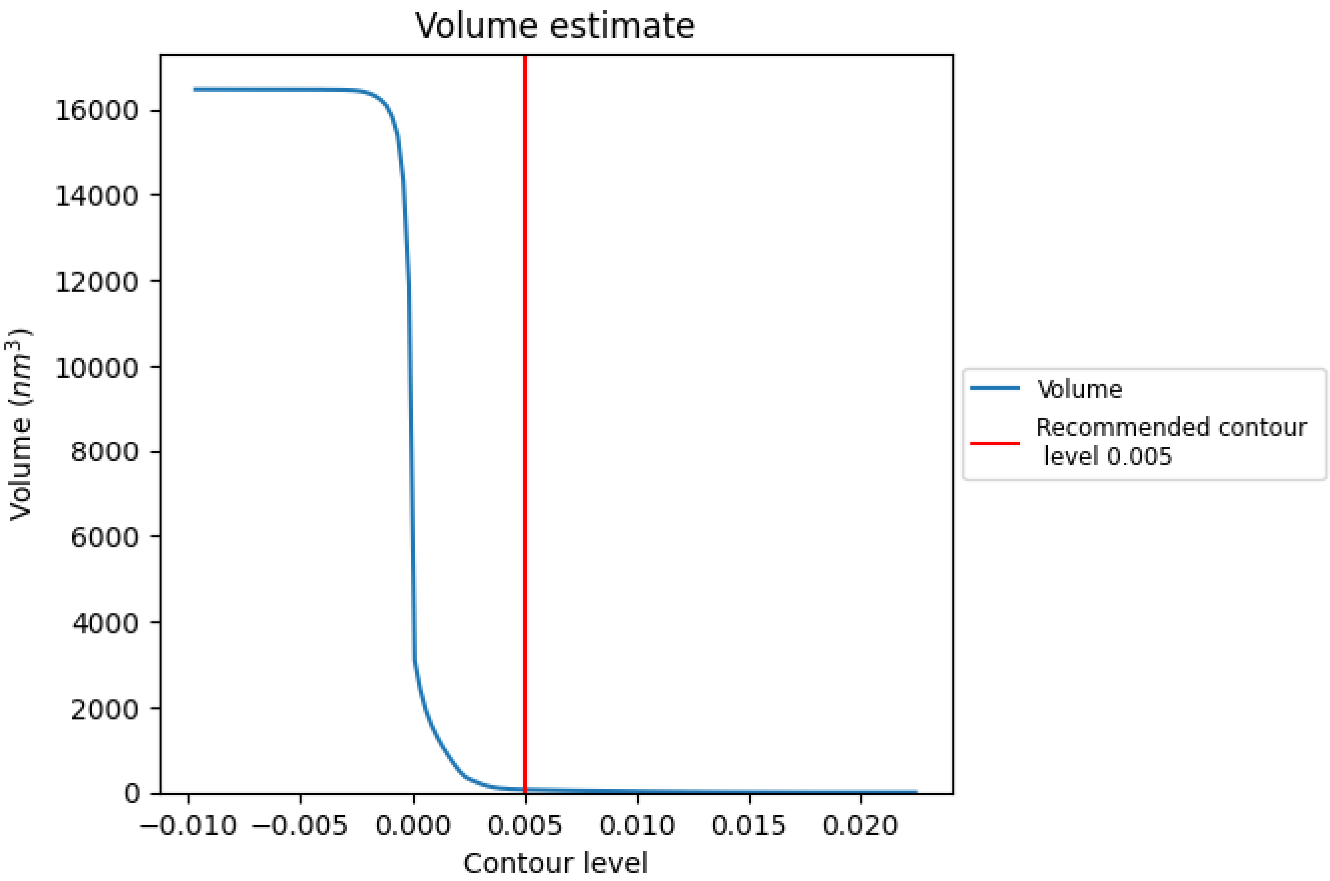


The volume at the recommended contour level is 66 $nm^3$; this corresponds to an approximate mass of 59 kDa.

The volume estimate graph shows how the enclosed volume varies with the contour level. The recommended contour level is shown as a vertical line and the intersection between the line and the curve gives the volume of the enclosed surface at the given level.

For Manuscript Review

#### 7.3 Rotationally averaged power spectrum (i)

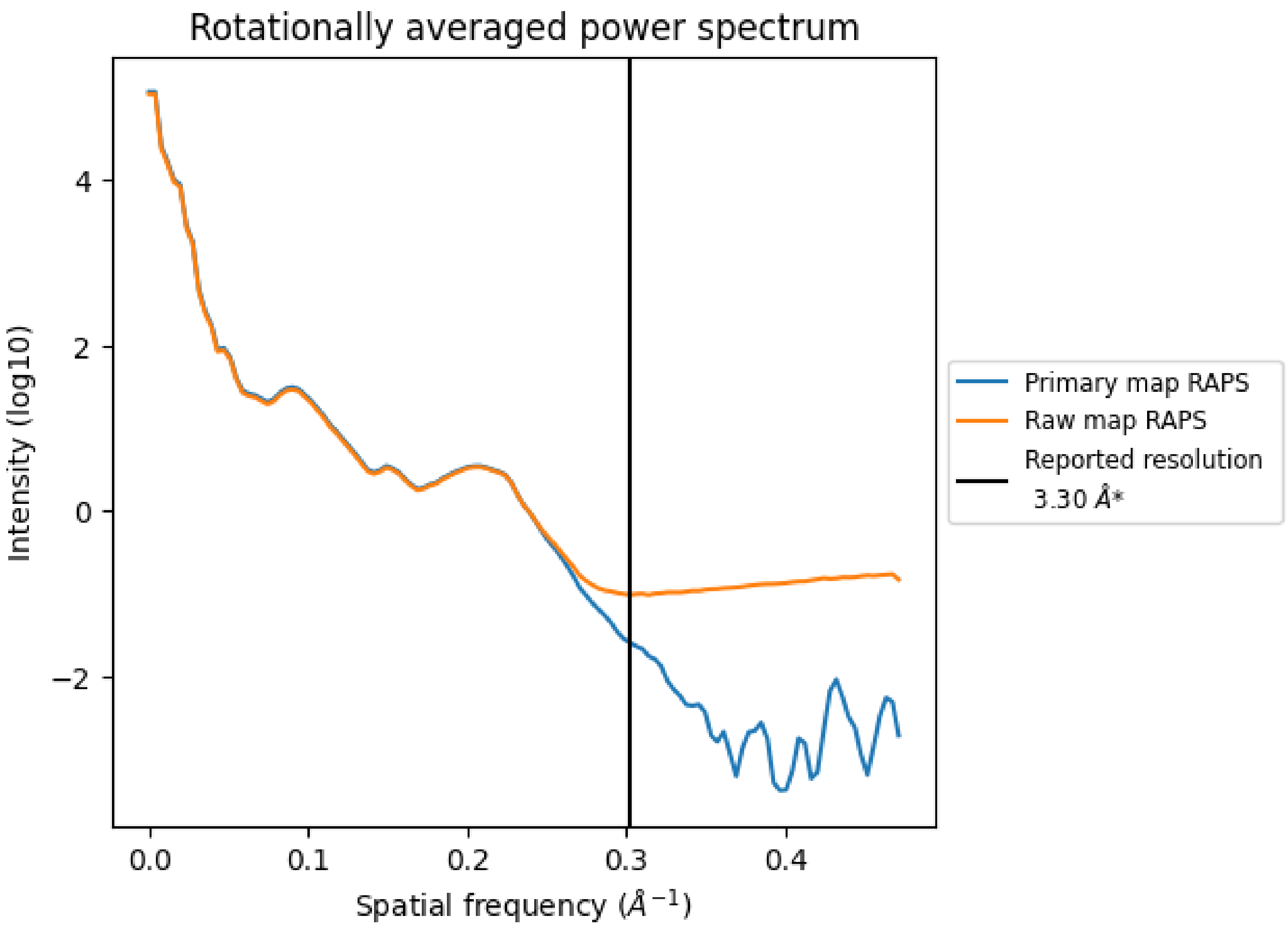


*Reported resolution corresponds to spatial frequency of 0.303 Å$^{-1}$

For Manuscript Review

### 8 Fourier-Shell correlation (i)

Fourier-Shell Correlation (FSC) is the most commonly used method to estimate the resolution of single-particle and subtomogram-averaged maps. The shape of the curve depends on the imposed symmetry, mask and whether or not the two 3D reconstructions used were processed from a common reference. The reported resolution is shown as a black line. A curve is displayed for the half-bit criterion in addition to lines showing the 0.143 gold standard cut-off and 0.5 cut-off.

#### 8.1 FSC (i)

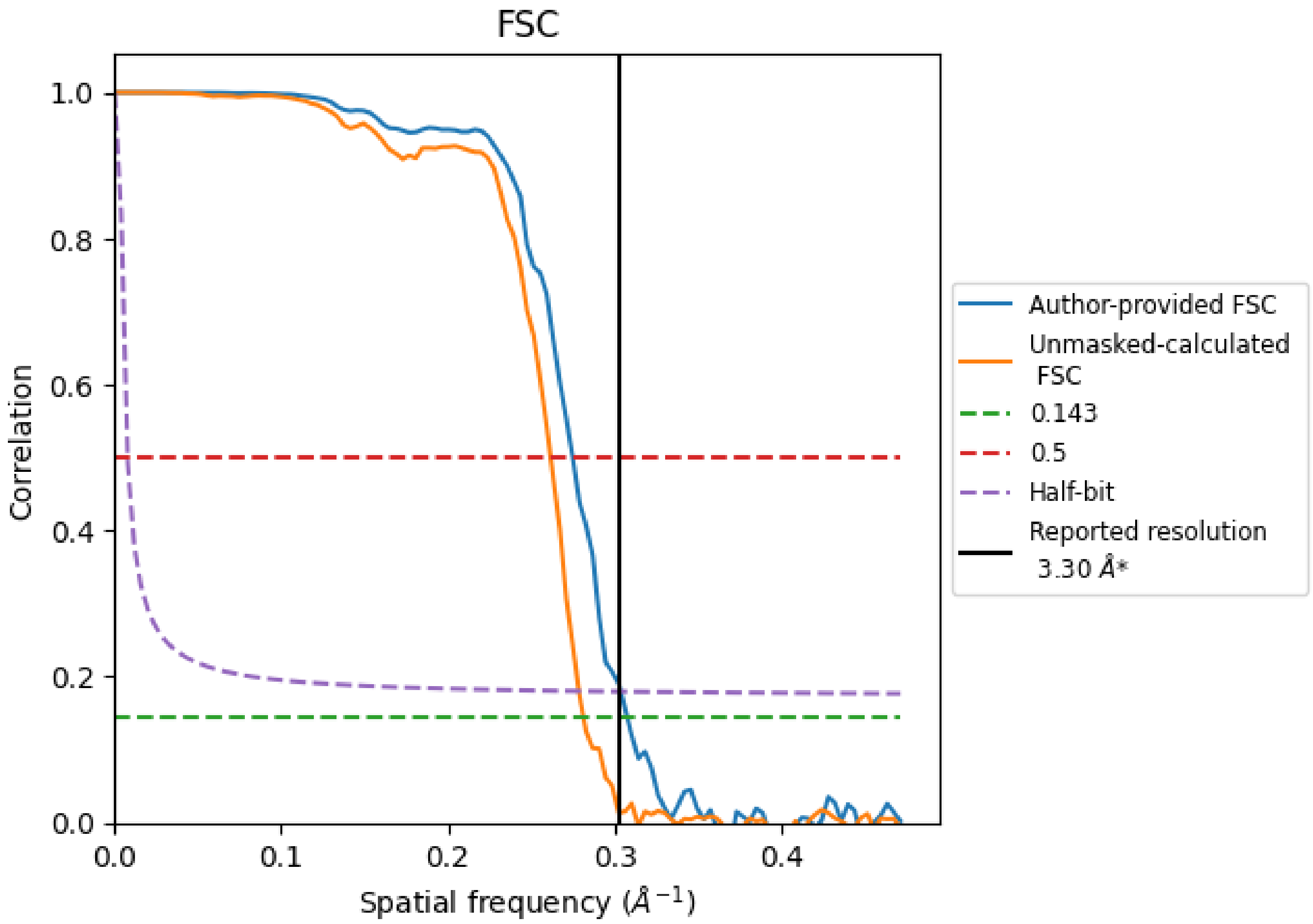


*Reported resolution corresponds to spatial frequency of 0.303 Å$^{-1}$

For Manuscript Review

#### 8.2 Resolution estimates (i)

| Resolution estimate (Å) | Estimation criterion (FSC cut-off) | | |
|---|---|---|---|
| | 0.143 | 0.5 | Half-bit |
| Reported by author | 3.30 | - | - |
| Author-provided FSC curve | 3.25 | 3.64 | 3.29 |
| Unmasked-calculated* | 3.55 | 3.82 | 3.59 |

*Resolution estimate based on FSC curve calculated by comparison of deposited half-maps.

For Manuscript Review

### 9 Map-model fit (i)

This section contains information regarding the fit between EMDB map EMD-35901 and PDB model 8J0F. Per-residue inclusion information can be found in section 3 on page 7.

#### 9.1 Map-model overlay (i)

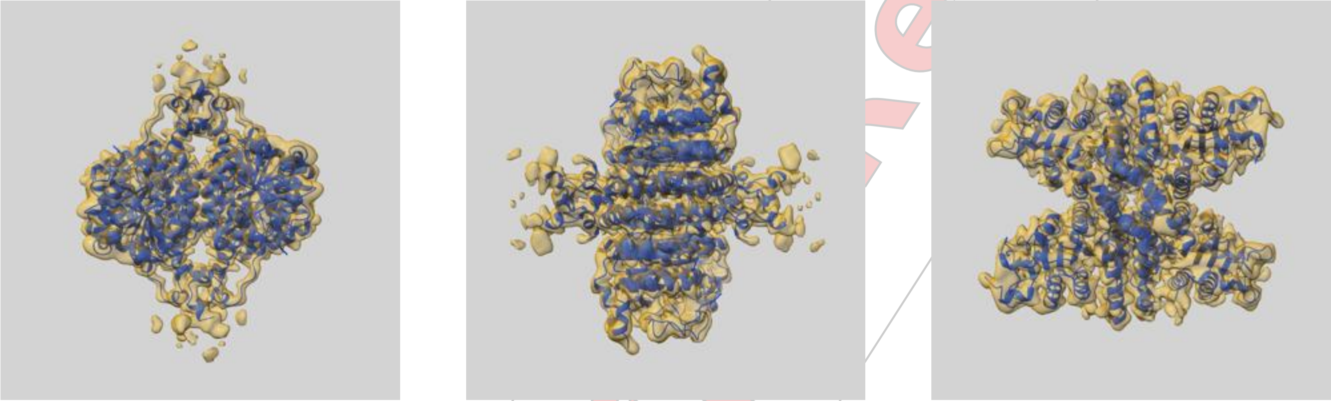

X Y Z

The images above show the 3D surface view of the map at the recommended contour level 0.005 at 50% transparency in yellow overlaid with a ribbon representation of the model coloured in blue. These images allow for the visual assessment of the quality of fit between the atomic model and the map.

For Manuscript Review

#### 9.2 Q-score mapped to coordinate model (i)

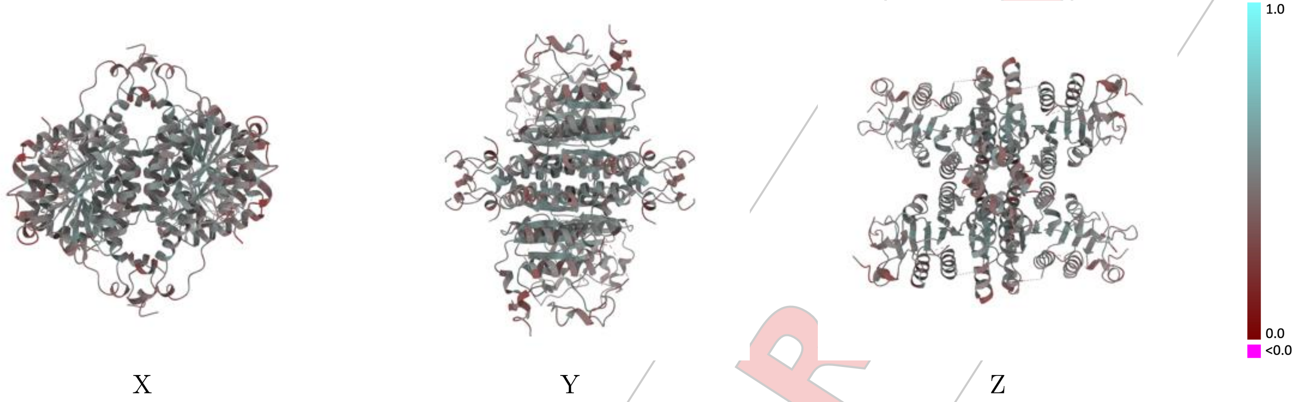


The images above show the model with each residue coloured according its Q-score. This shows their resolvability in the map with higher Q-score values reflecting better resolvability. Please note: Q-score is calculating the resolvability of atoms, and thus high values are only expected at resolutions at which atoms can be resolved. Low Q-score values may therefore be expected for many entries.

#### 9.3 Atom inclusion mapped to coordinate model (i)

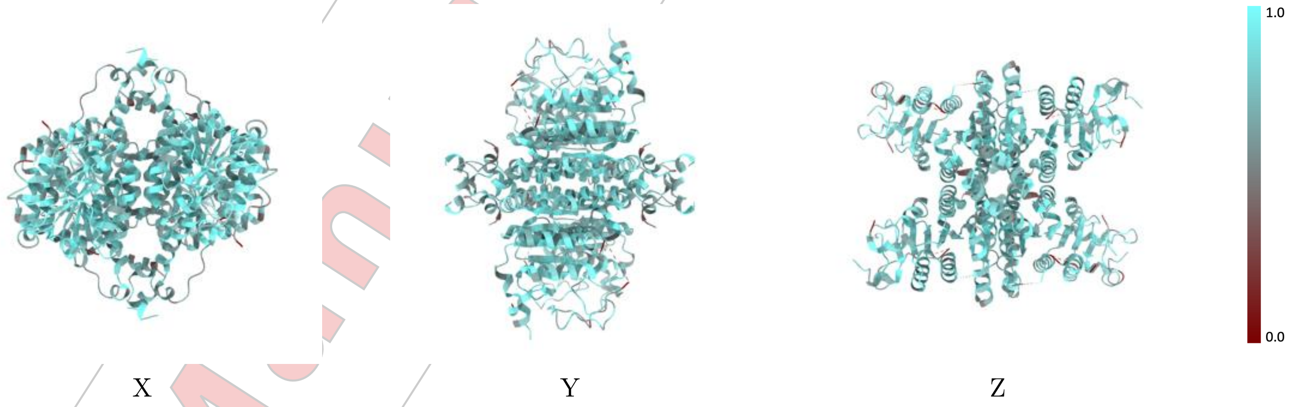


The images above show the model with each residue coloured according to its atom inclusion. This shows to what extent they are inside the map at the recommended contour level (0.005).

For Manuscript Review

#### 9.4 Atom inclusion (i)

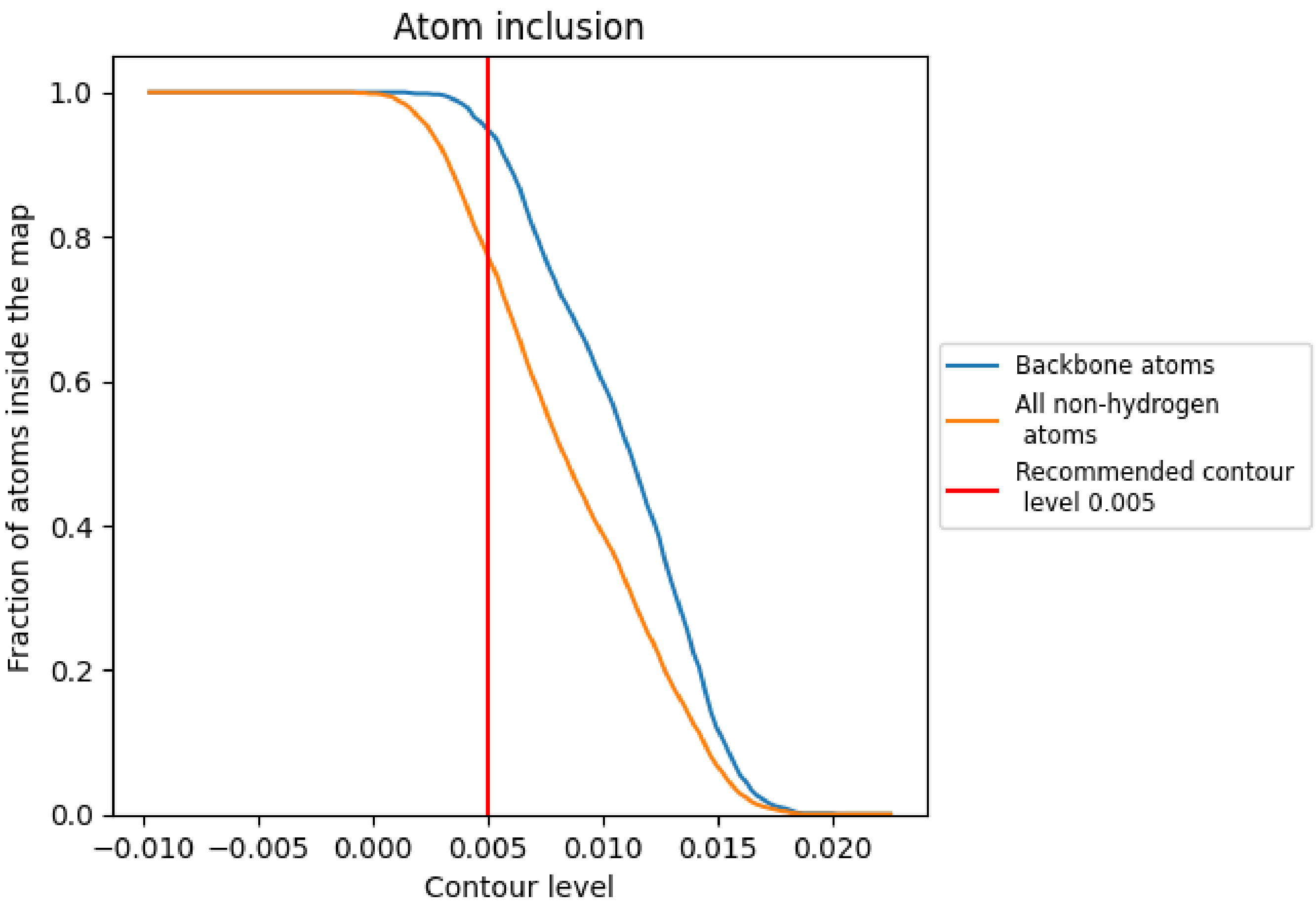


At the recommended contour level, 95% of all backbone atoms, 77% of all non-hydrogen atoms, are inside the map.

For Manuscript Review

#### 9.5 Map-model fit summary (i)

The table lists the average atom inclusion at the recommended contour level (0.005) and Q-score for the entire model and for each chain.

| Chain | Atom inclusion | Q-score |
|---|---|---|
| All | 0.7730 | 0.4560 |
| A | 0.7750 | 0.4600 |
| B | 0.7810 | 0.4600 |
| C | 0.7790 | 0.4590 |
| D | 0.7790 | 0.4600 |
| E | 0.6860 | 0.3880 |
| F | 0.6860 | 0.3920 |
| G | 0.6860 | 0.3980 |
| H | 0.6780 | 0.3900 |

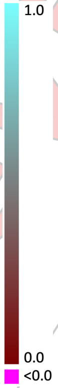


For Manuscript Review
